# Non-muscle myosin II regulates presynaptic actin assemblies and neuronal mechanobiology in *Drosophila*

**DOI:** 10.1101/2023.11.10.566609

**Authors:** Biljana Ermanoska, Jonathan Baets, Avital A. Rodal

## Abstract

Neuromuscular junctions (NMJs) are evolutionarily ancient, specialized contacts between neurons and muscles. They endure mechanical strain from muscle contractions throughout life, but cellular mechanisms for managing this stress remain unclear. Here we identify a novel actomyosin structure at *Drosophila* larval NMJs, consisting of a long-lived, low-turnover presynaptic actin core that co-localizes with non-muscle myosin II (NMII). This core is likely to have contractile properties, as manipulating neuronal NMII levels or activity disrupts its organization. Intriguingly, depleting neuronal NMII triggered changes in postsynaptic muscle NMII levels and organization near synapses, suggesting transsynaptic propagation of actomyosin rearrangements. We also found reduced levels of Integrin adhesion receptors both pre- and postsynaptically upon NMII knockdown, indicating disrupted neuron-muscle connections. Mechanical severing of axons caused similar actin core fragmentation and Integrin loss to NMII depletion, suggesting this structure responds to tension. Our findings reveal a presynaptic actomyosin assembly that maintains mechanical continuity between neurons and muscle, possibly facilitating mechanotransduction at the NMJ via Integrin-mediated adhesion.

## Introduction

Neuromuscular junctions (NMJs) and axons must weather continuous mechanical strain from muscle contractions, while also maintaining contact with significantly more rigid muscle tissue (Tyler, 2012). Thus, the NMJ offers an excellent tissue mechanics model to study evolutionarily conserved mechanisms by which “soft” neurons sustain their shape, function and plasticity while facing substantial mechanical challenges. NMJs are susceptible to aging and neurodegenerative disorders, and retraction of synaptic terminals precedes the degeneration of axons and subsequent loss of neuronal cell bodies in many diseases (Iyer, Shah and Lovering, 2021). Therefore, understanding how mechanical forces are generated, sensed, and transmitted within neurons and between neurons and their surrounding muscle is an important and understudied topic.

The actin cytoskeleton underlies multiple conserved mechanisms by which cells generate and respond to cellular forces (Michelot and Drubin, 2011). For example, endocytosis is supported by submicron-sized (∼200 nm), short-lived (<30s) force-generating actin patches, predominantly containing Arp2/3-nucleated branched filamentous actin (F-actin) (Akamatsu et al., 2020; Del Signore et al., 2021). On the other hand, cell and tissue-scale forces are often mediated by the actin-based motor non-muscle myosin II (NMII) (Murrell et al., 2015). Mammalian NMII is a hexamer composed of two heavy chains (NMII^HC^), two essential light chains (NMII^ELC^), and two regulatory light chains (NMII^LC^), and is present in a folded or extended conformation in cells (Sellers and Heissler, 2019). The transition from folded, auto-inhibited state to an extended filament is tightly controlled via phosphorylation of the NMII^LC^, which triggers the self-assembly of contraction-competent NMII into bipolar filaments, with motor domains at both ends that bind antiparallel actin filaments (Sellers, 1991). In this way, NMII induces sliding of the filaments, resulting in the generation of contractile forces for a variety of cellular needs (Vicente-Manzanares et al., 2009, Garrido-Casado et al., 2024; Quintanilla, Hammer and Beach, 2023). As one example, ventral stress fibers are micron-sized, long-lived assemblies of linear F-actin that recruit NMII and play a key role in cellular mechanics and force sensing in cultured cells (Livne and Geiger, 2016). Thus, depending on the specific force requirements, actin assemblies with variable sizes, lifetime, and molecular composition support different cellular processes.

Numerous actin-based and myosin-associated structures have been observed in axons and presynaptic terminals and could play important roles in the mechanical properties of neurons. An axonal submembranous cytoskeleton (SMC), composed of periodic repeats of spectrin, actin rings and non-muscle myosin II, was described in the past decade (Berger et al., 2018; Costa et al., 2020; Xu, Zhong and Zhuang, 2013). This structure is conserved in mammalian and invertebrate neurons (He et al., 2016; Qu et al., 2017), and supports periodic organization of numerous other molecules important for neurotransmission, cell adhesion and molecular transport (Zhou et al., 2022). A subset of the SMC represents radial actomyosin, which specifically regulates axonal diameter and conduction (Costa et al., 2020), the passage of larger cargos (Costa et al., 2020; Wang et al., 2020), and reversable changes upon mild mechanical stress (Pan et al., 2024). Another population of axonal myosin shows a more longitudinal distribution (Vassilopoulos et al., 2019); however it remains unknown if these longitudinal structures bridge neighboring actin rings,, associate with other axonal linear actin assemblies such as trails (Ganguly et al., 2015) or F-actin bundles (Gallo, 2006; Micinski and Hotulainen, 2024), or have yet another function. SMC periodicity diminishes at synaptic boutons, where actin is found in mesh-like assemblies at the active zone, rails between the active zone and deeper reserve pools, and corrals around the whole presynaptic compartment (Bingham et al., 2023), as well as in transient puncta with features of endocytic sites (Del Signore et al., 2021). Currently it is unknown if any of these assemblies recruit NMII, or how they contribute to the mechanical properties of presynaptic terminals.

Functional studies also implicate actomyosin structures in NMJ physiology. At the *Drosophila* larval NMJ, depletion of neuronal NMII disrupts the organization of synaptic vesicles (Seabrooke, Qiu and Stewart, 2010), suggesting this motor might associate with at least a fraction of presynaptic actin structures. Interestingly, both presynaptic overexpression or knockdown of NMII decreased spontaneous release of vesicles, suggesting that balanced NMMII levels are necessary for presynaptic synaptic transmission (Seabrooke and Stewart, 2011). In line with these findings, at the *Drosophila* embryonic NMJ, changing mechanical tension via axotomy can modulate synaptic vesicle organization and synaptic plasticity (Siechen et al., 2009). Moreover, neuronal depletion of *Drosophila* NMII at the larval NMJ promotes formation of nascent boutons by blebbing (Fernandes et al., 2023). These structures depend on neuronal activity and muscle contraction (Ataman et al., 2008; Fernandes et al., 2023; Piccioli and Littleton, 2014), and can sometimes mature into functional boutons (Ataman et al., 2008; Fernandes et al., 2023; Vasin et al., 2019), representing a form of activity-dependent structural plasticity. Thus, NMII, and by extension actomyosin contractility, are necessary to maintain the integrity of boutons experiencing muscle contractions and to regulate the growth of new synaptic arbors. Taken together, these findings highlight the ability of the NMJ to respond to mechanical forces. However, the subcellular organization and functions of the presynaptic actomyosin assemblies that underlie these events remain largely unknown.

Actomyosin assemblies coordinate with additional multi-protein machineries to control cellular mechanosensing and mechanotransduction. For example, ventral stress fibers are physically coupled to focal adhesion complexes (Livne and Geiger, 2016) along the ventral membrane of adherent cells. At the center of focal adhesion-mediated intra- and extracellular force-tuning are a set of metazoan Integrin receptors that bridge cytoplasmic cytoskeletal components to a large variety of extracellular matrix (ECM) molecules and counter-receptors on other cell types (Chastney, Conway and Ivaska, 2021). In this way, via biochemical and mechanical activation of Integrins, cells sense the mechanical features of their surroundings and transduce these forces into biological signals by inducing assembly and maturation of focal adhesions. The crucial role Integrins play in synaptic biology has been well established in vertebrates (Park and Goda, 2016) and at the *Drosophila* NMJ, where their disruption (Beumer et al., 1999; Orr, Fetter and Davis, 2022) or targeting their interactors (components of the ECM (Tsai et al., 2012; Wang et al., 2018), enzymes involved in ECM glycosylation (Dani, Zhu and Broadie, 2014), an Integrin activator (Lee et al., 2017), or the cytoskeletal linker Talin (Orr, Fetter and Davis, 2022)), induce a range of morphological defects and affect activity-dependent synaptic plasticity. These studies highlight the importance of Integrins in neuronal function and position these molecules as likely candidates to control transsynaptic neuron-to-muscle communication and NMJ mechanobiology.

In this study, we examined presynaptic actin assemblies at the *Drosophila* larval NMJ and identified a long-lived, low-turnover linear actin core traversing the axon terminal. NMII regulates the organization of this core, suggesting that these are actomyosin structures, likely with contractile properties. Strikingly, neuronal depletion of NMII induced decrease of NMII^LC^ levels together with an increase in aggregate-like formations of NMII^HC^ in the postsynaptic muscle, suggesting that neuronal actomyosin rearrangements can propagate their effects transsynaptically. Next, we considered Integrin receptors as mediators of the transsynaptic NMII changes and found a decrease in Integrin foci both pre- and postsynaptically, indicating that neuronal actomyosin defects sensed via local Integrin receptors can propagate to the surrounding muscle tissue. In sum, we demonstrate that a previously unrecognized subpopulation of presynaptic actomyosin maintains the mechanical continuum from neuron to muscle, likely regulating the mechanobiology of the neuromuscular junction.

## Results

### Discrete presynaptic actin assemblies at the Drosophila NMJ

At the *Drosophila* larval NMJ, presynaptic terminals innervate the mechanically active postsynaptic muscle, making this synapse an ideal model to study biological solutions to mechanical challenges at the neuron-muscle interface (**Fig. 1A**). In a previous study, we demonstrated that a subpopulation of patch-like, submicron sized, presynaptic actin is amenable to live imaging via confocal microscopy and subsequent quantitative analysis (Del Signore et al., 2021). To improve the visualization of these small and transient actin structures, we re-designed a broadly used, genetically encoded F-actin marker GMA (<u>G</u>FP <u>M</u>oesin <u>A</u>ctin-binding domain) (Bloor and Kiehart, 2001; Del Signore et al., 2021), by tagging it with the bright mNeonGreen (Shaner et al., 2013) fluorescent protein (mNGMA). To assess the structure and function of presynaptic actin we generated QF-driven, QUAS-mNGMA (QmNGMA) transgenes. This allowed us to genetically combine two binary expression systems (UAS/Gal4 and QUAS/QF) (Brand and Perrimon, 1993; Potter et al., 2010) in a single fly and to visualize presynaptic actin, while independently manipulating different actin-associated proteins via UAS-RNAi lines, without affecting the expression levels of the F-actin marker. Finally, to improve the resolution of the presynaptic actin structures of interest, we used Airyscan microscopy.

**Fig. 1.**
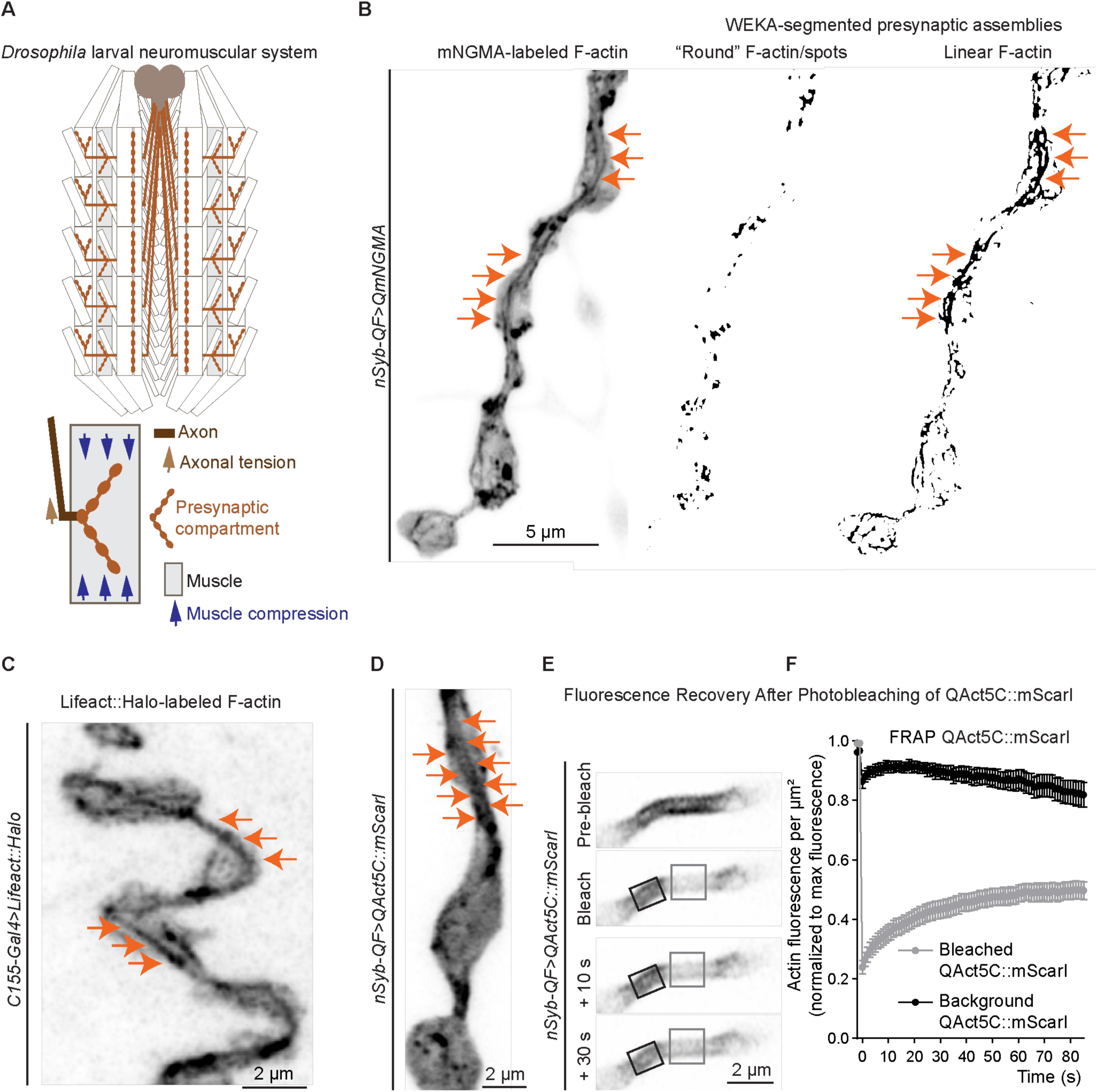
Presynaptic actin cytoskeleton at the *Drosophila* larval NMJ. **A**) Diagram of the *Drosophila* larval neuromuscular system as a tissue mechanics model. The NMJ at muscle 4 (gray) was predominantly assessed in this study. The presynaptic compartment (light brown) is subjected to multiple mechanical forces including axonal tension (brown arrow) and compression due to muscle contractions (blue arrows). **B**) A single frame from live imaging of presynaptic actin assemblies visualized via neuronal expression of the genetically encoded actin marker mNeonGreen-GMA (mNGMA) in larval preparations with intact brains (see also **Supp. Movie S1**). WEKA segmentation was used to distinguish different actin subpopulations (round (spot-like), and linear, (cable-like or core) F-actin structures) and to facilitate their subsequent quantitative analysis (see also **Fig. S1**). Orange arrows indicate a linear actin core traversing the NMJ. **C**) A single frame from live imaging of NMJs expressing an independent genetically encoded actin marker Lifeact::Halo delineates similar spot-like and cable-like (core) actin assemblies. Orange arrows indicate the linear actin core. **D**) A single frame from live imaging of the *Drosophila* actin isoform 5C, tagged with the red fluorescent protein mScarlet-I (QAct5C::mScarI), similarly delineates the actin core in addition to round F-actin assemblies. **E**) Image sequence of FRAP of QAct5C::mScarI-labeled actin core. **F**) FRAP curve of QAct5C::mScarI (N – NMJs, error bars represent SEM). Scale bars – 5 µm in A, 2 µm in other images. **See Table 1 for detailed genotypes.**

With these tools in hand, we imaged presynaptic actin in live dissected larvae with intact brains and axons, at rest. Interestingly, along with numerous spot-like F-actin assemblies throughout the NMJ, the improved signal-to-noise and resolution provided by our approach also revealed a previously unrecognized linear, cable-like actin structure traversing the NMJ **(Fig. 1B**, **Supp. Movie S1**). To enable unbiased analysis of these structures we built a WEKA machine learning-based classifier (Arganda-Carreras et al., 2017) in the image analysis software FIJI (Schindelin et al., 2012) that allowed us to specifically segment round from linear F-actin structures at the NMJ (**Fig. 1B**, **Fig. S1**). Next, we verified the existence of the linear actin core with the broadly used F-actin marker, Lifeact (Riedl et al., 2008), tagged with the Halo self-labeling protein tag (*UAS-Lifeact::Halo*). Neuronally expressed Lifeact::Halo (*C155-Gal4>Lifeact::Halo*) labeled the linear actin core as well as the previously observed spot-like structures at the NMJ (**Fig. 1C**). Thus, we confirmed that the newly identified linear actin assemblies can be visualized with at least two independent, bright actin markers in live larval preparations. We used Lifeact and GMA interchangeably in our experiments, as both markers label comparable F-actin structures at the NMJ.

In the course of our live imaging, we noticed that the presynaptic actin core is stable (in space and time) over minutes (**Fig. S2A**), unlike the transient, spot-like structures, which have lifetimes in the tens of seconds and likely represent endocytic events (Del Signore et al., 2021) (**Fig. S2B**). In addition, fluorescent recovery after photobleaching (FRAP) of the QmNGMA marker demonstrated an immediate relabeling of the same, practically unchanged presynaptic actin core structure, further supporting the idea that these are long-lived assemblies on which the F-actin marker can readily bind and unbind (**Fig. S2C-D**). To explore the dynamics of actin directly, we generated a QUAS transgene carrying the Actin5C *Drosophila* isoform, tagged with the bright, red fluorescent protein mScarlet-I (QAct5C::mScarI) (Bindels et al., 2017). Live imaging of QAct5C::mScarI showed that tagged actin integrated into spot-like and linear F-actin structures along the core, albeit less continuous compared to QmNGMA (**Fig. 1D**, **Fig. S2E**). FRAP analysis of QAct5C::mScarI demonstrated that the F-actin assemblies constituting the presynaptic actin core have slow but detectable recovery (**Fig. 1E-F**), suggesting that they do not turn over quickly. To independently test dynamics, we evaluated the sensitivity of the linear actin core to treatment with the actin-sequestering drug Latrunculin A/LatA. Incubation of dissected larvae for 30 minutes in 1 µM LatA led to the disappearance of distinct actin structures at the NMJ, including the core, while incubation with the DMSO solvent alone did not lead to loss of actin structures (**Fig. S2F-G**). These results indicate that the core does undergo dynamic turnover. In sum, we identified different presynaptic actin assemblies that vary in size and lifetime and are probably involved in distinct functions at synaptic terminals.

Next, we asked if the linear F-actin structures at the NMJ are similar to previously described axonal linear F-actin structures (D’Este et al., 2015; Gallo, 2006; Ganguly et al., 2015; Micinski and Hotulainen, 2024). We assessed QmNGMA-labeled actin in axonal bundles proximal to cell bodies in the ventral ganglion, and at distal axons exiting the nerve bundle to connect to NMJs (**Fig. S3A-B**). In proximal axon bundles the distributions of fluorescent QmNGMA and free GFP were indistinguishable both visually or by analysis of WEKA-segmented structures, possibly due to the narrow diameter of these axons, precluding any conclusion about actin structures in this region (**Fig. S3C**). By contrast, in distal axons, we observed stretches of linear F-actin structures aligned with the axonal shaft, in contrast to neuronally-expressed GFP which filled the entire shaft and NMJ without labeling specific structures (**Fig. S3B**). Thus, linear F-actin structures exist in axons but it is inconclusive whether they exist throughout the axon or represent a continuum from the cell body to the actin core at NMJs.

### Actin-associated proteins at the presynaptic actin core

A large number of actin-associated proteins (AAPs) define the assembly, shape, and stability of actin structures (Pollard, 2016). For example, actin nucleators such as the Arp2/3 complex and formins facilitate the generation of branched and linear actin filaments, respectively. Previously, we demonstrated that most of the endocytic actin patches at the NMJ co-localize with components of the Arp2/3 actin filament nucleation complex (Del Signore et al., 2021). On the other hand, pharmacological inhibition of formins disrupted presynaptic protein composition, vesicle cycling, and endocytosis in induced synapses and hippocampal neurons (Bingham et al., 2023; Wen et al., 2016). Together, these studies suggest that distinct actin subpopulations supporting synaptic processes might be generated via different F-actin nucleators. We assessed the presence of nucleation machinery along the presynaptic actin core by imaging NMJs in larvae co-expressing Lifeact::Halo (to label F-actin) and GFP-tagged Arp3 or the formin Dia. We observed that a fraction of both Arp3::GFP (**Fig. S4A**; **Supp. Movie S2**) and Dia::eGFP (**Fig. S4B**; **Supp. Movie S3**) punctate signal were visible along the linear actin core, unlike the diffuse distribution of control cytosolic GFP (**Fig. S4C**; **Supp. Movie S4**). Thus, we did not detect distinct enrichment of Arp3 and Dia on the presynaptic F-actin structures, suggesting that both nucleation mechanisms could contribute to their assembly.

To further evaluate the molecular constituents of the linear presynaptic actin structures, we investigated the localization of non-muscle tropomyosins. Tropomyosins are important F-actin stabilizers, which co-polymerize with actin in linear structures such as yeast actin cables (Alioto et al., 2016) and stress fibers in cultured mammalian cells (Tojkander et al., 2011), and are also integral components of the periodic axonal cytoskeleton and determinants of neuronal morphology and function (Abouelezz et al., 2020; Brettle, Patel and Fath, 2016). To ask if *Drosophila* non-muscle Tropomyosin 1/Tm1 associates with the linear presynaptic actin core, we generated transgenic flies containing QUAS-driven, mNeonGreen-tagged versions of two of the 18 predicted *Drosophila* Tm1 isoforms (QmN-Tm1-A and QmN-Tm1-L). When expressed in neurons (with *nSyb-QF*) and imaged live, both QTm1 isoforms labeled linear structures traversing the NMJ (**Fig. S5A-B**). Compared to the control cytosolic GFP, Q-Tm1-A co-localized significantly more strongly with the actin marker Lifeact::Halo along the actin core (**Fig. S5C-D**). In addition, Q-Tm1-L signal was sensitive to the same LatA treatment that depolymerized Q-mNGMA-labeled actin structures (**Fig. S5E-F**), consistent with Tm1 association with the F-actin core. Finally, we performed immunostaining with an antibody that recognizes Tm1-A and Tm1-L (Cho et al., 2016), and observed a distribution resembling the presynaptic actin core in control animals (**Fig. S5G**). The Tm1 signal was strongly decreased at NMJs of larvae expressing RNAi targeting all Tm1 isoforms (*panTm1^43542^*) or an independent line targeting Tm1-A (*Tm1^56869^*) (**Fig. S5G-H**), confirming its specificity. Importantly, this result indicates that the core is an endogenous structure and does not depend on expression of an F-actin marker. In sum, the linear presynaptic actin core is decorated by the F-actin stabilizer Tropomyosin, giving this structure a molecular signature resembling stress fibers (Tojkander et al., 2011).

Ventral stress fibers are actomyosin structures that sense and coordinate the adhesion of cultured cells to their substrates (Livne and Geiger, 2016). We asked if the linear presynaptic actin core would similarly recruit NMII. *Drosophila* NMII consists of NMII^HC^, encoded by Zipper/Zip, and NMII^LC^, encoded by Spaghetti squash/Sqh. We used a well-established Sqh::GFP transgene, expressed from its own promoter (Royou et al., 2004) to evaluate its co-localization with neuronally driven Lifeact::Halo. In live larvae, we observed Sqh::GFP as punctate submicron-sized structures localized partially along F-actin and dispersed throughout the bouton, in contrast to the diffuse GFP control signal, which was uniformly distributed and lacked specific localization, suggesting that Sqh::GFP associates with discrete domains within the NMJ (**Fig. S4D**, **Supp. Movie S5**).

Next, we examined the distribution of endogenous NMII^HC^/Zip via immunostaining of control NMJs (**Fig. 2A**) with a previously described antibody (Sokac and Wieschaus, 2008). We further validated antibody specificity by confirming a decrease in Zip signal in neurons and muscles upon expression of Zip RNAi (*Zip^65947^*) in each of these tissues (**Fig. 2B-C, Fig. S6E-G**). We detected Zip as puncta with a range of sizes enriched in and around the HRP-labeled neuronal terminals, some of which aligned along the NMJ core, where we observe the F-actin linear structures (**Fig. 2A**). Interestingly, we also identified a subpopulation of larger aggregate-like Zip structures with smooth edges, localized predominantly in muscles (**Fig. 2A**). Finally, we examined if Zip associates with the presynaptic F-actin assemblies by Supplementary Movie 11neuronally co-expressing Zip::GFP and Lifeact::Halo (*C155-Gal4>Lifeact::Halo>Zip::GFP^OE^*) (**Fig. 2D-E, Supp. Movie S6**). Similar to endogenous NMII^HC^/Zip staining, Zip::GFP labeled puncta of different sizes throughout boutons, and largely colocalized with Lifeact::Halo in both round and linear F-actin assemblies (**Fig. 2D-F)**. Interestingly, some of the Zip::GFP puncta were organized as doublets, resembling the organization of NMII bipolar filaments (**Fig. 2E**). These Zip doublets were spaced with an average distance of ∼260 nm in maximum intensity projection images (**Fig. 2E, G)**, slightly lower than the bipolar filament range of 280 – 400 nm in stress fibers (Svitkina et al., 1989). This tighter spacing may reflect different orientations of bipolar filaments along the *z* axis, or a contracted state (Russell et al., 2011). Some Zip::GFP doublets were detectable along linear F-actin structures, suggesting that they could form actomyosin assemblies, while others were not obviously correlated with F-actin and might represent constitutive NMII minifilament assembly (Vasquez, Tworoger and Martin, 2014). These results suggest that the linear actin structure and NMII together might form a presynaptic actomyosin core with contractile properties.

**Fig. 2.**
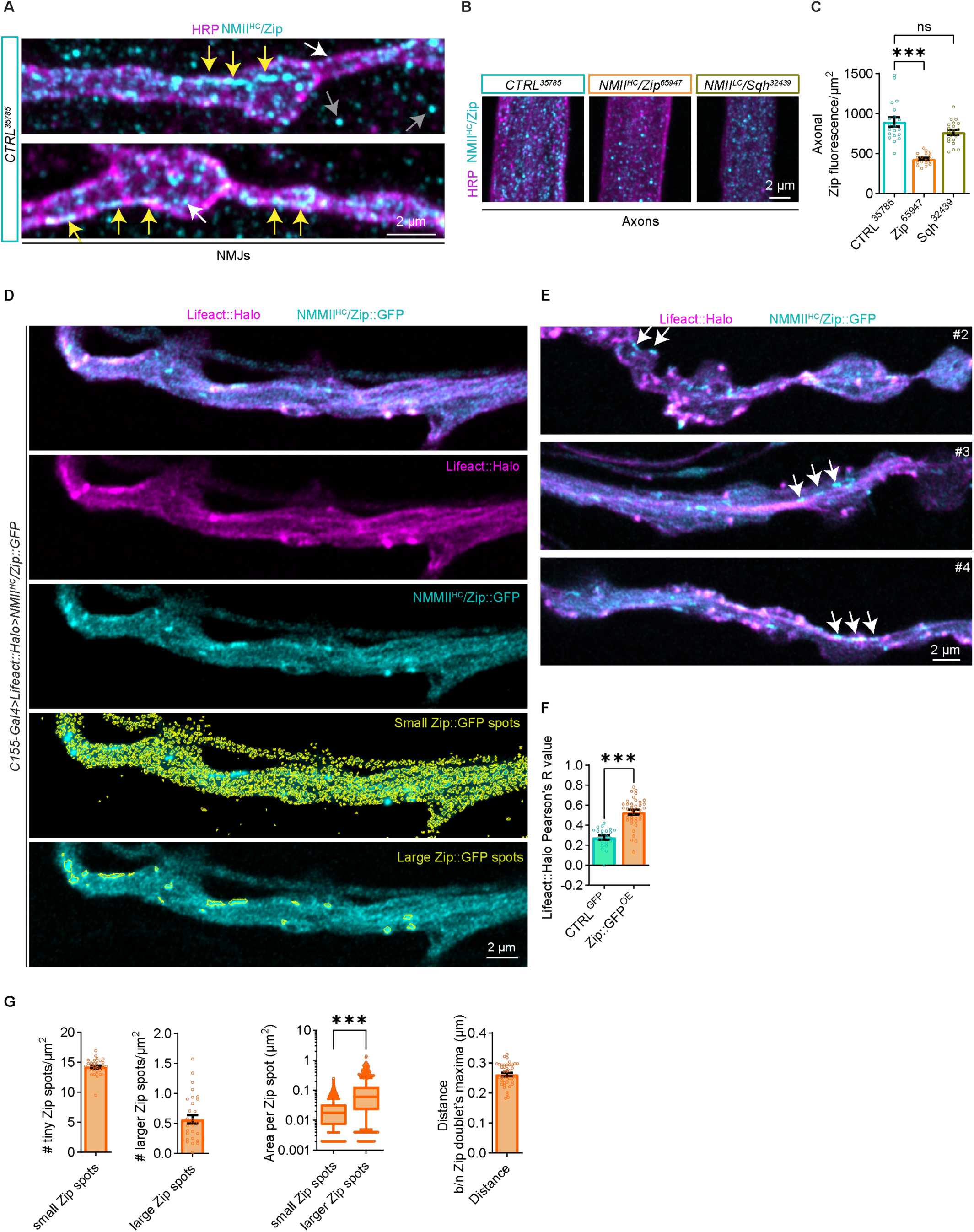
Distribution of endogenous and neuronally expressed NMII^HC^/Zip at *Drosophila* larval NMJs. **A**) Immunostaining of endogenous NMII^HC^/Zip (cyan) at NMJs (HRP, magenta) of wild-type larvae labels large and small puncta at the NMJ and in muscles (yellow arrows – large Zip, white arrows – small Zip structures). Smooth, aggregate-like Zip puncta are visible predominantly in muscles (gray arrows). **B**) Zip puncta (cyan) in distal axonal bundles (HRP in magenta) in animals neuronally (*C155-Gal4*) expressing *CTRL^35785^*-, *NMII^HC^/Zip^65947^*and *NMII^LC^/Sqh^32439^* RNAi lines. **C**) Quantification of Zip fluorescence in axonal bundles. **D**) Live imaging of neuronally co-expressed NMII^HC^/Zip::GFP (cyan) and Lifeact::Halo (magenta) similarly reveals Zip puncta with different size at the NMJ, some of which are enriched at spot-like and linear F-actin assemblies (see also **Supp. Movie S6**). **E**) Independent examples of live-imaged NMJs of larvae expressing Zip::GFP and Lifeact::Halo. White arrows point to putative bipolar Zip filaments. **F**) Pearson’s R correlation between Lifeact::Halo and Zip::GFP. **G**) Quantification of the abundance and size of smaller and larger Zip puncta at the NMJ, as well as the average distance between adjacent Zip puncta in putative bipolar Zip filaments. In bar graphs with linear Y-axis, error bars represent SEM. N in bar graphs – NMJs; N in box and whiskers – individual Zip assemblies. *** p<0.001 upon unpaired, non-parametric Mann-Whitney test. Scale bars – 2 µm. **See Tables 1 for detailed genotypes and N.**

### Non-muscle myosin II regulates the integrity of the presynaptic actin core

Ventral stress fiber contractility and structural integrity depend on both NMII recruitment and mechanical tension (Livne and Geiger, 2016). To assess how NMII might regulate the presynaptic actin cytoskeleton, we imaged QmNGMA-labeled F-actin in control- and Zip RNAi/*Zip^65947^*-expressing larvae. We observed reorganization of both round and linear presynaptic actin assemblies (**Fig. 3A-B**; **Supp. Movies S7-8**). We implemented our WEKA classifier to segment and analyze these assemblies as separate categories (see **Fig. S1A-C** for details). While overall F-actin levels were comparable in both genotypes (**Fig. 3C**), we detected a decrease in the number of actin spots (**Fig. 3D**) and in the percentage of NMJ area covered with linear F-actin (**Fig. 3E**). The remaining round F-actin structures in *Zip^65947^* mutants were comparable in size and brightness to controls (**Fig. 3D**). On the other hand, the linear actin assemblies were smaller and dimmer in mutant NMJs as compared to controls (**Fig. 3E**, **Supp. Movies S7-8**). We observed and quantified a comparable disorganization of presynaptic actin assemblies in fixed *Zip^65947^*larvae (**Fig. S7A-B**). Interestingly, neuronal overexpression of Zip::GFP similarly reduced the area and brightness of both round and linear presynaptic F-actin structures visualized via Lifeact::Halo, compared to overexpression of GFP alone (**Fig. S7C-E**), suggesting that the organization of presynaptic actin is dependent on regulated levels of NMII^HC^/Zip.

**Fig. 3.**
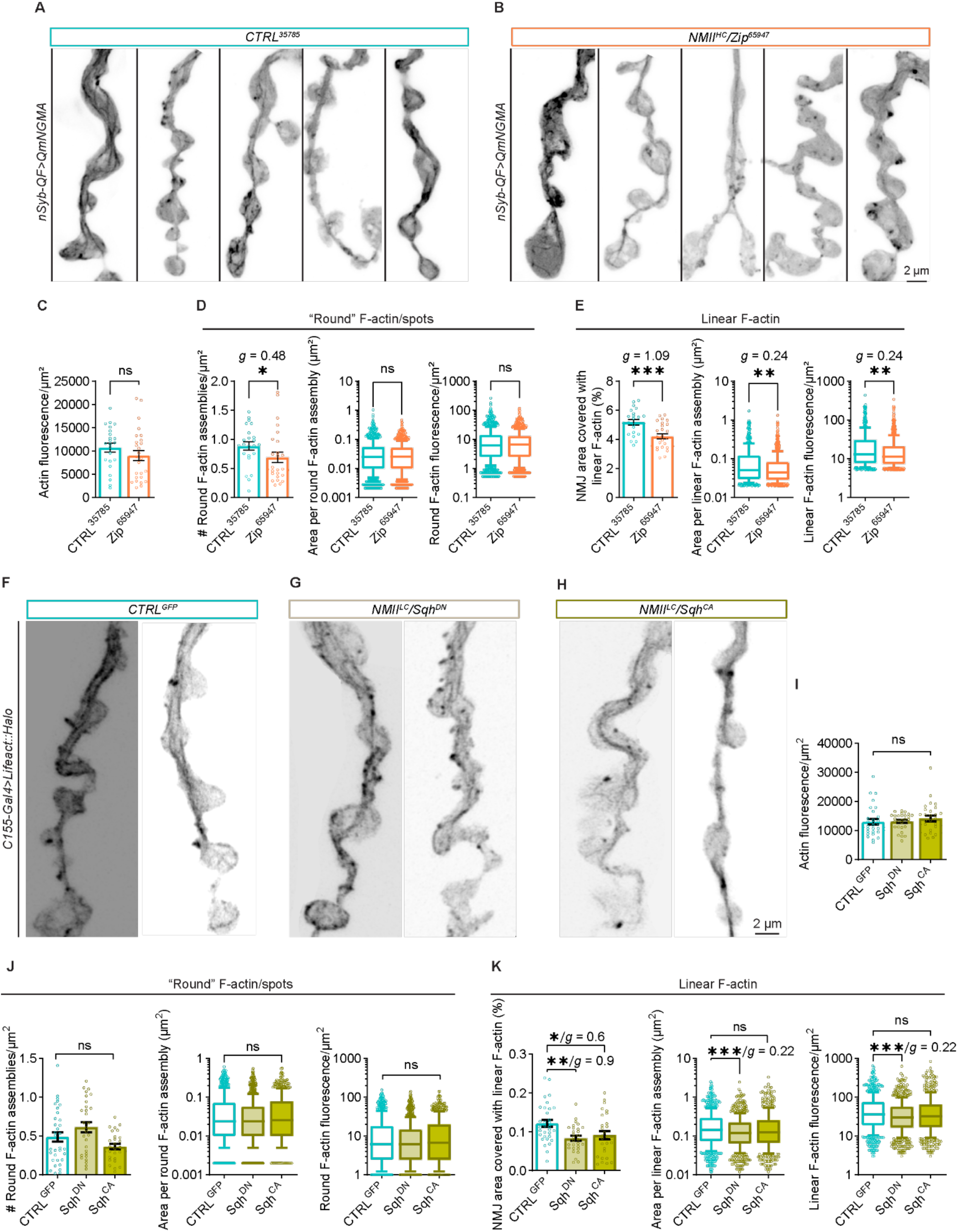
Non-muscle Myosin II regulates the integrity of presynaptic actin assemblies. **A**) QmNGMA-labeled presynaptic actin in larvae co-expressing *CTRL^35785^*- or **B**) *NMII^HC^/Zip^65947^*RNAi (see also **Supp. Movies S7-8**). Representative NMJs are from independent larvae. **C**) Quantification of fluorescence per µm^2^ for the QmNGMA actin marker in whole NMJs. **D**) Quantifications of the number of round F-actin assemblies per µm^2^, as well as their area and fluorescence. **E**) Graphs representing the percentage of NMJ area covered with linear F-actin, as well as the area and fluorescence of the individual assemblies. **F-H**) Lifeact::Halo-labeled presynaptic F-actin in *GFP*-expressing control, or larvae expressing unphosphorylatable Sqh^DN^ and constitutively phosphorylated Sqh^CA^, respectively. **I**) Quantification of fluorescence per µm^2^ for the Lifeact::Halo F-actin marker in whole NMJs. **J**) Quantifications of the number of round F-actin assemblies per µm^2^, as well as their area and fluorescence. **K**) Graphs representing the percentage of NMJ area covered with linear F-actin, as well as the area and fluorescence of the individual assemblies. Box and whiskers graphs were used to represent the results of the area and fluorescence intensity of the individual presynaptic actin structures. Whiskers represent 10^th^ to 90^th^ percentile, while the rest of the data points are shown as individual values. The Y-axis in these graphs represents log10, to capture the broad distribution of the individual values. In bar graphs with linear Y-axis, error bars represent SEM. N in bar graphs – NMJs; N in box and whiskers – individual actin assemblies. ns – not significant, * p<0.05, ** p<0.01, and *** p<0.001 upon unpaired, non-parametric Mann-Whitney test. *g* – Hedges’ *g* represents effect size. Scale bars – 2 µm. **See Table 1 for detailed genotypes and N.**

We next examined how NMII activation state affects presynaptic actin organization by manipulating phosphorylation of NMII^LC^/Sqh at conserved residues Thr20/Ser21, which regulate the transition from folded, contraction-incompetent to unfolded, contraction-competent NMII (Jordan and Karess, 1997; Sellers, 1991). Using Lifeact::Halo to visualize F-actin, we expressed either non-phosphorylatable (Sqh^A20A21^/Sqh^DN^) or phosphomimetic (Sqh^E20E21^/Sqh^CA^) Sqh mutants to alter the balance between inactive and active NMII (**Fig. 3F-H**). Total F-actin levels at mutant NMJs remained comparable to GFP-expressing controls (**Fig. 3I**). The Sqh phosphomutants did not change the size and intensity of round F-actin structures, and led to a minor, not statistically significant variation in their abundance (**Fig. 3J**). Both mutants also significantly reduced the area covered by linear F-actin, with Sqh^DN^ causing an additional decrease in the size and intensity of remaining linear structures (**Fig. 3K**). Together with our Zip knockdown results, these data demonstrate that both NMII levels and activation state regulate presynaptic actin organization, particularly the linear actin core.

### Presynaptic NMII- and actin rearrangements can be sensed and propagated transsynaptically

While exploring presynaptic actomyosin structures, we detected dynamic Sqh::GFP puncta at NMJs and in the postsynaptic muscle proximal to boutons (**Fig. S4D, Supp. Movie S5**). This area contains the subsynaptic reticulum, a postsynaptic membrane specialization enriched with ion channels, cell adhesion molecules, and receptors that facilitate neuron-to-muscle contact (Ataman, Budnik and Thomas, 2006). Postsynaptic Sqh signal was also visible in fixed controls (**Fig. 4A**). Neuron-specific expression of two independent Zip RNAi lines (*Zip^65947^* and *Zip^36727^*) decreased Sqh at the NMJ terminals (**Fig. 4A- B**), as well as in axons (**Fig. S6A-B**). Surprisingly, we also observed a decrease in postsynaptic Sqh::GFP signal, both in the region proximal to the bouton as well as in general in the muscle (**Fig. 4A-B**, **Fig. S6C**). We ruled out the possibility that the postsynaptic Sqh decrease is due to leaky expression of the transgenic *UAS-RNAi* lines in the muscle in the absence of the neuronal driver *C155-Gal4* (**Fig. S6D**). WEKA-based analysis of segmented Sqh::GFP-positive particles in the α- HRP-delineated- and postsynaptic NMJ area (**Fig. 4C)** confirmed a significant decrease in the number, size, and brightness of the NMII^LC^/Sqh structures in *Zip* mutants compared to controls (**Fig. 4D-E**). These data demonstrate that neuronal NMII^HC^/Zip depletion results in transsynaptic decrease of the regulatory light chain subunit Sqh of the motor in the muscle.

**Fig. 4.**
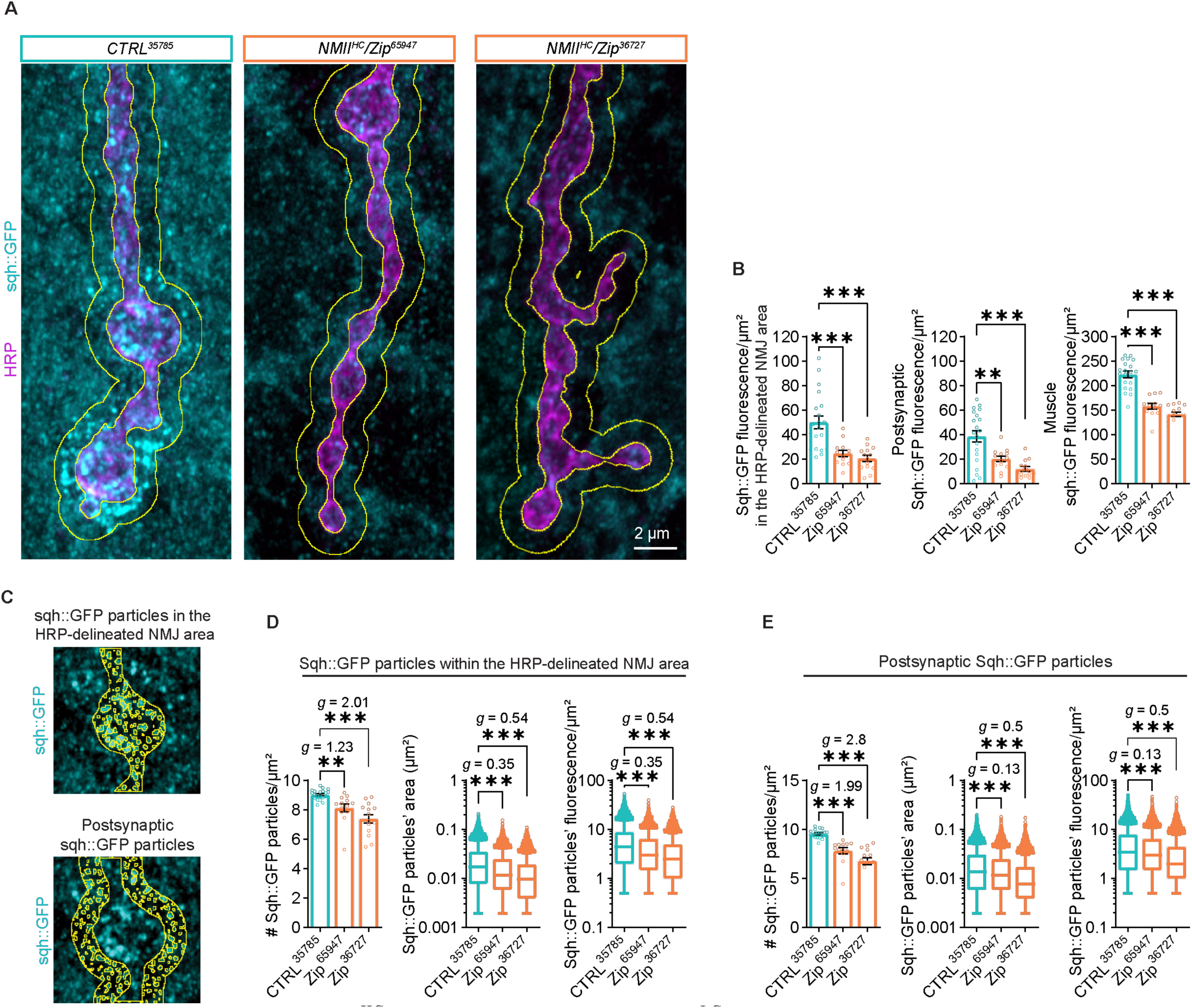
Depletion of neuronal NMII^HC^ reduces the levels of the NMII^LC^ subunit in the muscle. **A**) Distribution of Sqh::GFP at NMJs in fixed larvae expressing *CTRL^35785^*, *Zip^65947^*, and *Zip^36727^* RNAi in neurons (*C155-Gal4*). Sqh::GFP fluorescence was analyzed within the α-HRP-delineated NMJ area (inner yellow outline) as well as 1 µm outside the neuron in the postsynaptic area (between inner and outer yellow outline). **B**) Analysis of the Sqh::GFP fluorescence in the HRP-delineated-, postsynaptic-, and muscle ROIs. **C**) Example of WEKA-segmented individual sqh::GFP particles in the HRP-delineated- and postsynaptic compartments. **D**) Analysis of the number, area and fluorescence WEKA-segmented sqh::GFP particles within the HRP-delineated NMJ area. **E**) Analysis of the number, area and fluorescence of postsynaptic WEKA-segmented sqh::GFP particles. Box and whiskers graphs were used to represent the results of the area and fluorescence intensity of the individual Sqh particles. Whiskers represent 10^th^ to 90^th^ percentile, while the rest of the data point are shown as individual values. The Y-axis in these graphs represents log10, to capture the broad distribution of the individual values. In bar graphs with linear Y-axis, error bars represent SEM. N in bar graphs – NMJs and muscle ROIs; N in box and whiskers – individual Sqh particles. ** p<0.01, *** p<0.001 after one-way ANOVA with Šídák’s multiple comparisons test. *g* – Hedges’ *g* represents effect size. Scale bar – 2 µm. **See Table 1 for detailed genotypes and N.**

We next examined if neuronal NMII depletion affects endogenous NMII^HC^/Zip levels and distribution in muscles (**Fig. 5A**), with our validated Zip antibody and Zip RNAi lines (**Fig. 2B-C; Fig. S6E-G**). We also validated RNAi lines targeting Sqh (**Fig. S6A-B**). The high density of Zip structures near the α-HRP-delineated NMJ region made it impossible to definitively resolve presynaptic structures at our imaging resolution. We therefore focused our analysis on Zip organization further into the muscle. Using WEKA-based image analysis of muscle Zip immunofluorescence, we distinguished two Zip populations in muscle that differ by an order of magnitude in size: smaller Zip spots and larger Zip aggregates. We then tested transsynaptic effects of neuronal NMII depletion. Upon *C155-Gal4* expression of *NMII^HC^*/*Zip^65947^* and *NMII^LC^/Sqh^32439^* (**Fig. 5A**), we observed an increased abundance, size, and brightness of Zip spots in the postsynaptic region surrounding boutons (**Fig. 5B**). Additionally, in muscles of *Zip^65947^*and *Sqh^32439^* larvae, we noted a greater number of Zip aggregates compared to controls (**Fig. 5C**). Similar Zip aggregates were previously reported in Sqh null- and phosphomimetic mutants and likely reflect assembly of NMII minifilaments when Sqh/Zip stoichiometry is altered, rather than protein denaturation (Jordan and Karess, 1997; Royou et al., 2004; Vasquez, Tworoger and Martin, 2014). Together, these results demonstrate that neuronal NMII depletion triggers transsynaptic changes in both light and heavy chain organization, reducing Sqh::GFP puncta while promoting Zip aggregate formation in muscles.

**Fig. 5.**
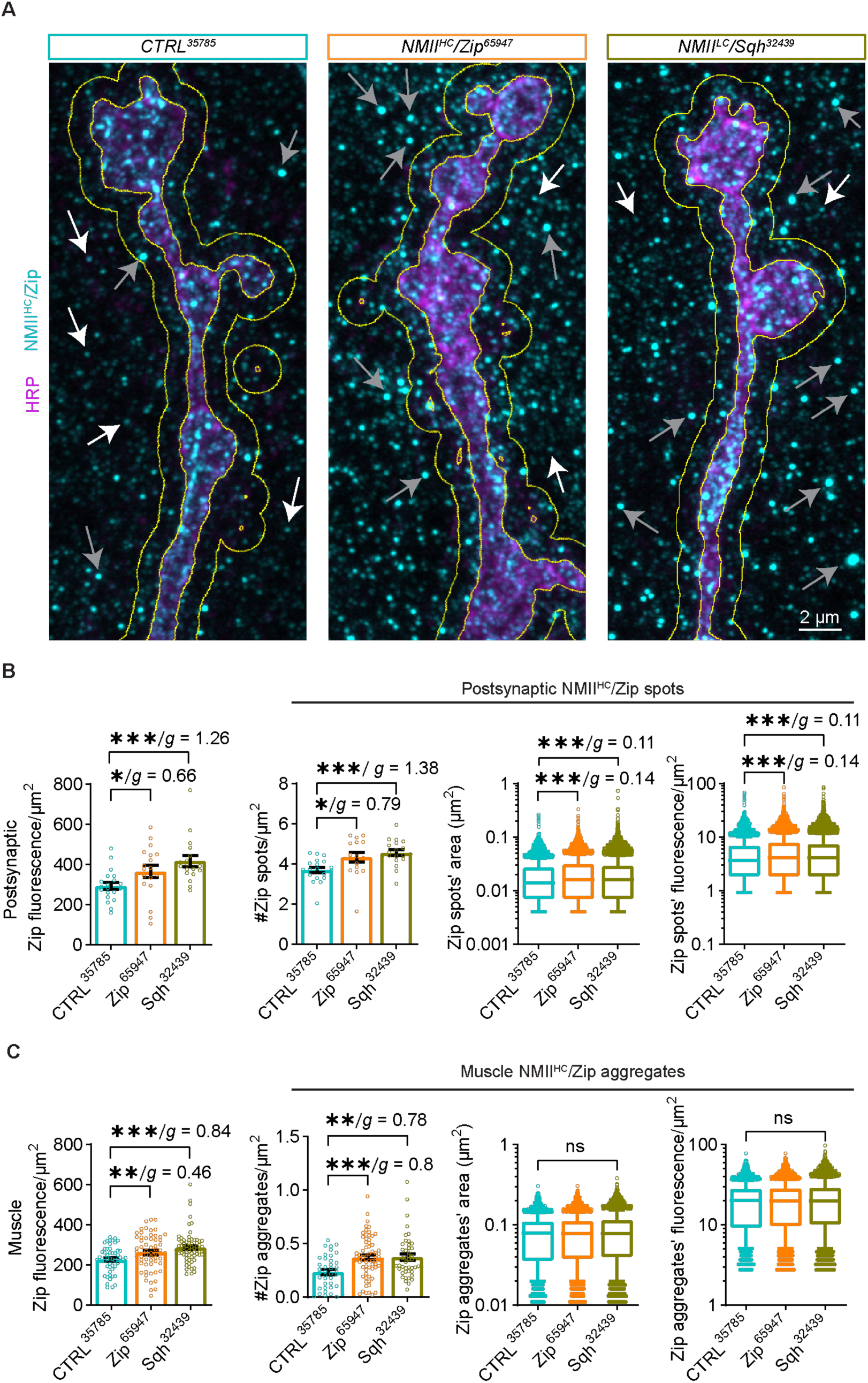
Depletion of neuronal NMII induces rearrangements of NMII^HC^/Zip pre- and postsynaptically. **A**) Endogenous NMII^HC^/Zip (cyan) at NMJs (HRP – magenta) in fixed larvae expressing *CTRL^35785^*, *Zip^65947^*, and *Sqh^32439^* RNAi in neurons (*C155-Gal4*); white arrows – Zip puncta, gray arrows – Zip aggregate-like puncta. **B**) Quantification of Zip fluorescence measured between the presynaptic area (masked by the neuronal HRP signal, inner yellow outline) and 1 µm outside the neuron in the postsynaptic area (outer yellow outline), along with analysis of the number, area and fluorescence of WEKA-segmented individual Zip particles in this postsynaptic compartment. N in bar graphs – NMJs; N in box and whiskers – individual Zip assemblies. **C**) Quantification of Zip fluorescence in different muscle ROIs, along with analysis of the number, area and fluorescence of WEKA-segmented individual Zip aggregate-like structures in muscles. N in bar graphs – NMJs or muscle ROIs; N in box and whiskers – individual Zip particles. Box and whiskers graphs were used to represent the results of the area and fluorescence intensity of the individual Zip particles. Whiskers represent 10^th^ to 90^th^ percentile, while the rest of the data point are shown as individual values. The Y-axis in these graphs represents log10, to capture the broad distribution of the individual values. In bar graphs with linear Y-axis, error bars represent SEM. ** p<0.01, *** p<0.001 after one-way ANOVA with Kruskal-Wallis multiple comparisons test. *g* – Hedges’ *g* represents effect size. Scale bar – 2 µm. **See Table 1 for detailed genotypes and N.**

### Presynaptic actomyosin depletion alters the organization of Integrin-β receptors

The linear presynaptic actin core shares key features with mechanosensitive structures like stress fibers, including its molecular composition, dependence on NMII-mediated contractility, and ability to propagate cytoskeletal changes between cells. We therefore hypothesized that disrupting these neuronal actomyosin assemblies drives the observed transsynaptic effects on NMII levels and organization. Stress fibers transmit mechanical signals through Integrin-based adhesions. Therefore we investigated whether the presynaptic actomyosin core similarly cooperates with transmembrane Integrin receptors by examining the distribution of the *Drosophila* Integrin-β PS subunit Myospheroid/Mys. We used a ubiquitously expressed Integrin-β PS subunit, tagged with YFP (ubiMys::YFP) (Yuan et al., 2010) to visualize Mys in live, dissected larvae (**Fig. 6A**), and a well-characterized antibody (Dani, Zhu and Broadie, 2014; Wang et al., 2018) to visualize Integrin- β in fixed larvae (**Fig. 6B**). In live larvae, we observed a subset of ubiMys::YFP Integrin-β formations aligned along the Lifeact::Halo-labeled linear actin core (**Fig. 6A**, **Fig. S8A**, **Supp. Movie S9**). Similar linear Integrin-β formations were detectable in fixed larvae (**Fig. 6B**), as observed in a previous report (Wang et al., 2018). We also detected spot-like Integrin-β entities of different sizes both pre- and postsynaptically (**Fig. 6B**), possibly reflecting different stages of maturation of Integrin structures (Kechagia, Ivaska and Roca-Cusachs, 2019; Livne and Geiger, 2016; Orr, Fetter and Davis, 2022), and suggesting a complex fine-tuning of cellular forces at the NMJ.

**Fig. 6.**
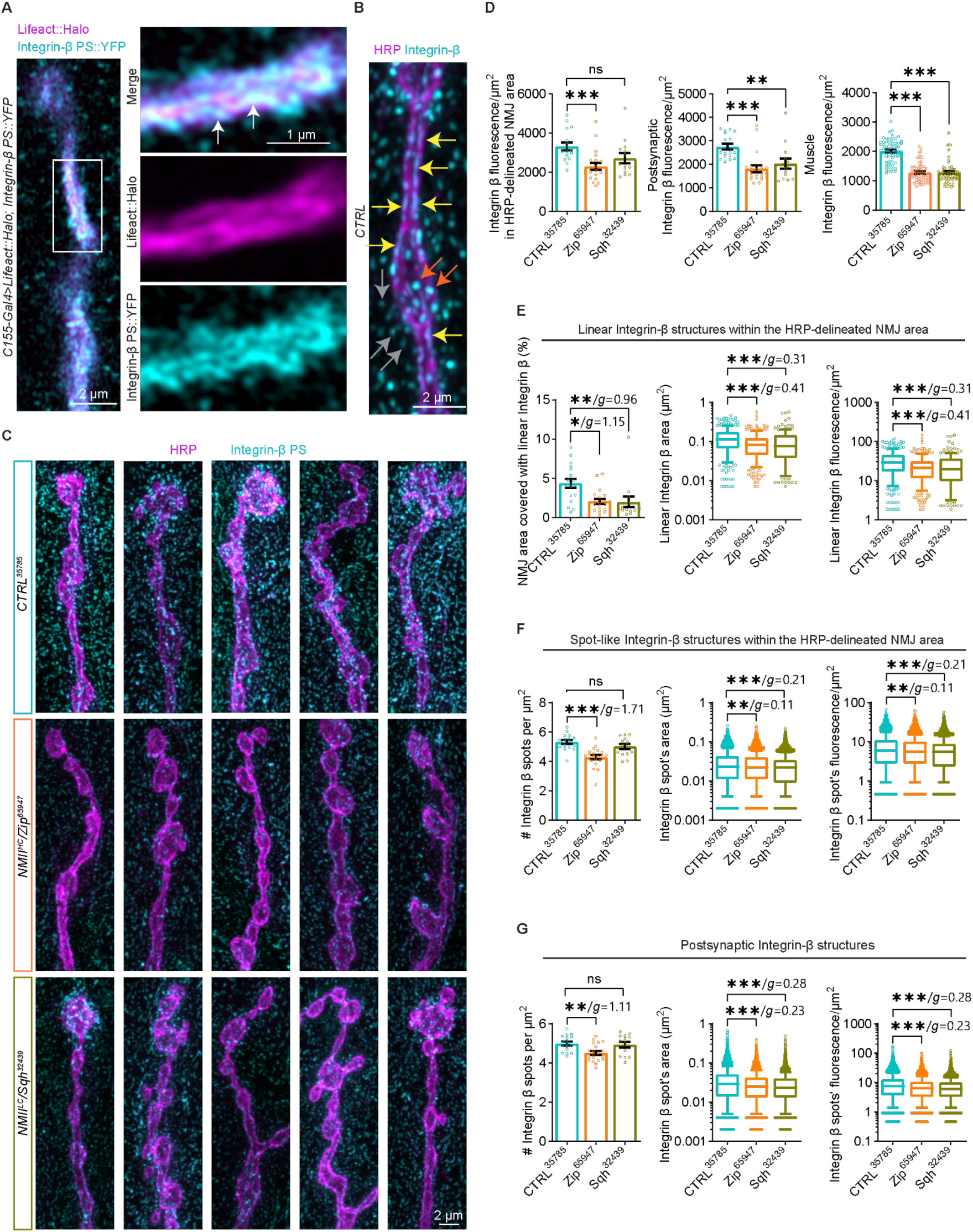
Neuronal depletion of non-muscle Myosin II rearranges Integrin receptors at the NMJ. **A**) Distribution of YFP-tagged, ubiquitously expressed Integrin-β (cyan) at NMJs in live larvae expressing Lifeact::Halo (magenta) in neurons (see also **Supp. Movie S9**). White arrows point to areas of overlap between Integrin-β and linear F-actin. **B**) Immunostaining of Integrin-β (cyan) at NMJs of control larvae. Yellow arrows depict Integrin-β in linear structures, red arrows point to presynaptic spot-like Integrin-β structures, while gray arrows point to postsynaptic spot-like Integrin-β. **C**) Reorganization of Integrin-β at NMJs of larvae expressing *NMII^HC^*/*Zip^65947^* and *NMII^LC^*/*Sqh^32439^*in neurons, compared to controls. **D**) Integrin-β fluorescence in the HRP-delineated-, postsynaptic-, and muscle ROIs. **E**) Quantifications of the abundance, size and fluorescence of WEKA-segmented linear Integrin-β structures within the HRP-delineated area. **F**) Analysis of the number, size and fluorescence of WEKA-segmented Integrin-β foci within the HRP-delineated area. **G**) Analysis of the number, size and fluorescence of WEKA-segmented postsynaptic Integrin-β foci. Box and whiskers graphs were used to represent the results of the area and fluorescence intensity of the individual Integrin-β particles. Whiskers represent 10^th^ to 90^th^ percentile, while the rest of the data points are shown as individual values. The Y-axis in these graphs represents log10, to capture the broad distribution of the individual values. In bar graphs with linear Y-axis, error bars represent SEM. N in bar graphs – NMJs or muscle ROIs; N in box and whiskers – individual Integrin-β assemblies. Ns – non-significant, * p<0.05, ** p<0.01, and *** p<0.001 after one-way ANOVA with Kruskal-Wallis multiple comparisons test. *g* – Hedges’ *g* represents effect size. Scale bars – 2 µm, inset in **A**) with scale bar – 1 µm. **See Table 1 for detailed genotypes and N.**

We next examined how neuronal NMII knockdown affects Integrin-β organization. *NMII^HC^/Zip^65947^* expression significantly reduced Integrin-β levels as well as the abundance, size and intensity of both linear and spot-like Integrin-β structures at the α-HRP-delineated NMJ area (**Fig. 6C-F**). *NMII^LC^/Sqh^32439^* expression similarly perturbed Integrin-β entities at the α-HRP- delineated NMJ area (**Fig. 6C-F**). Both *Zip^65947^*and *Sqh^32439^* significantly decreased postsynaptic Integrin-β levels near boutons and reduced the size and intensity of postsynaptic spots (**Fig. 6D and G**). These effects were reproduced with an independent *Zip^36727^* RNAi line (**Fig. S8B-C**). Two controls confirmed the specificity of our finding: Zip RNAi without the *Gal4* driver had no effect on Integrin-β levels (**Fig. S8D-F**), and Integrin-β levels at myotendinous junctions (in the muscle but far from the NMJ) were unchanged in *C155-Gal4>Zip^65947^* larvae (**Fig. S8G-H**). These results suggest that neuronal NMII depletion triggers local reorganization of Integrin-β in proximity to the NMJ.

Finally, we examined the effects of neuronal Integrin-β knockdown using an *Integrin-β/Mys^33642^* RNAi that effectively decreased Integrin-β levels in axons (**Fig. S9A-B**), though not at the NMJ, where it is likely masked by the high density of Integrin structures near the α-HRP-delineated NMJ region (**Fig. S9C-D**). Therefore, we analyzed Integrin-β outside this region. Strikingly, though overall postsynaptic and muscle Integrin-β levels remained unchanged (**Fig. S9D**), the postsynaptic Integrin-β structures were smaller and dimmer in *Mys^33642^* (**Fig. S9E-F**). This demonstrates that neuronal down- regulation of Integrin-β can trigger changes in postsynaptic Integrin-β organization without affecting total muscle levels, ruling out ectopic Gal4 expression in the muscle as a trivial explanation for the effects we observed in the muscle upon neuronal NMII manipulation. Instead, these results demonstrate that purely neuronal perturbations can trigger specific changes in postsynaptic protein organization through transsynaptic signaling. Together with our NMII knockdown results, these data suggest that presynaptic actomyosin is mechanically coupled to transmembrane Integrin receptors, allowing force-dependent signals to propagate across the neuromuscular junction.

### Mechanical severing of neurons causes rearrangement of presynaptic linear actin in control but not NMII^HC^-depleted neurons

To complement our chronic NMII depletion studies, we examined how acute disruption of neuronal tension affects the presynaptic actin core by performing axotomy near the ventral ganglion (**Fig. 7A)**, similar to studies of the effects of mechanical tension on synaptic vesicle distribution (Siechen et al., 2009), albeit at different developmental stage. This procedure is widely used in studies of the NMJ, as it prevents excessive muscle contraction in larval fillets, while the preparation maintains muscle potential and capacity for evoked neurotransmitter release (Feng, Ueda and Wu, 2004). We analyzed presynaptic actin assemblies labeled with neuronal QmNGMA, in larvae with intact versus cut axons; larvae were imaged live within ∼20 minutes after axotomy (**Fig. 7B**, **Supp. Movies S10-11**). While total F-actin levels and structure numbers remained unchanged after axotomy (**Fig. 7C**), WEKA-based analysis revealed that severed axons had larger and brighter round actin structures (**Fig. 7D**) but smaller and dimmer linear F-actin assemblies (**Fig. 7E**). This reorganization resembles the effects of NMII depletion. To test whether NMII mediates this acute response, we performed axotomy in *Zip^65947^* larvae (**Fig. 7F**). Despite reduced overall F-actin levels (**Fig. 7G**), axotomy in NMII-depleted neurons increased round F-actin structures but failed to further alter linear F-actin (**Fig. 7H-I**), suggesting that NMII is required for tension-dependent remodeling of the presynaptic actin core.

**Fig. 7.**
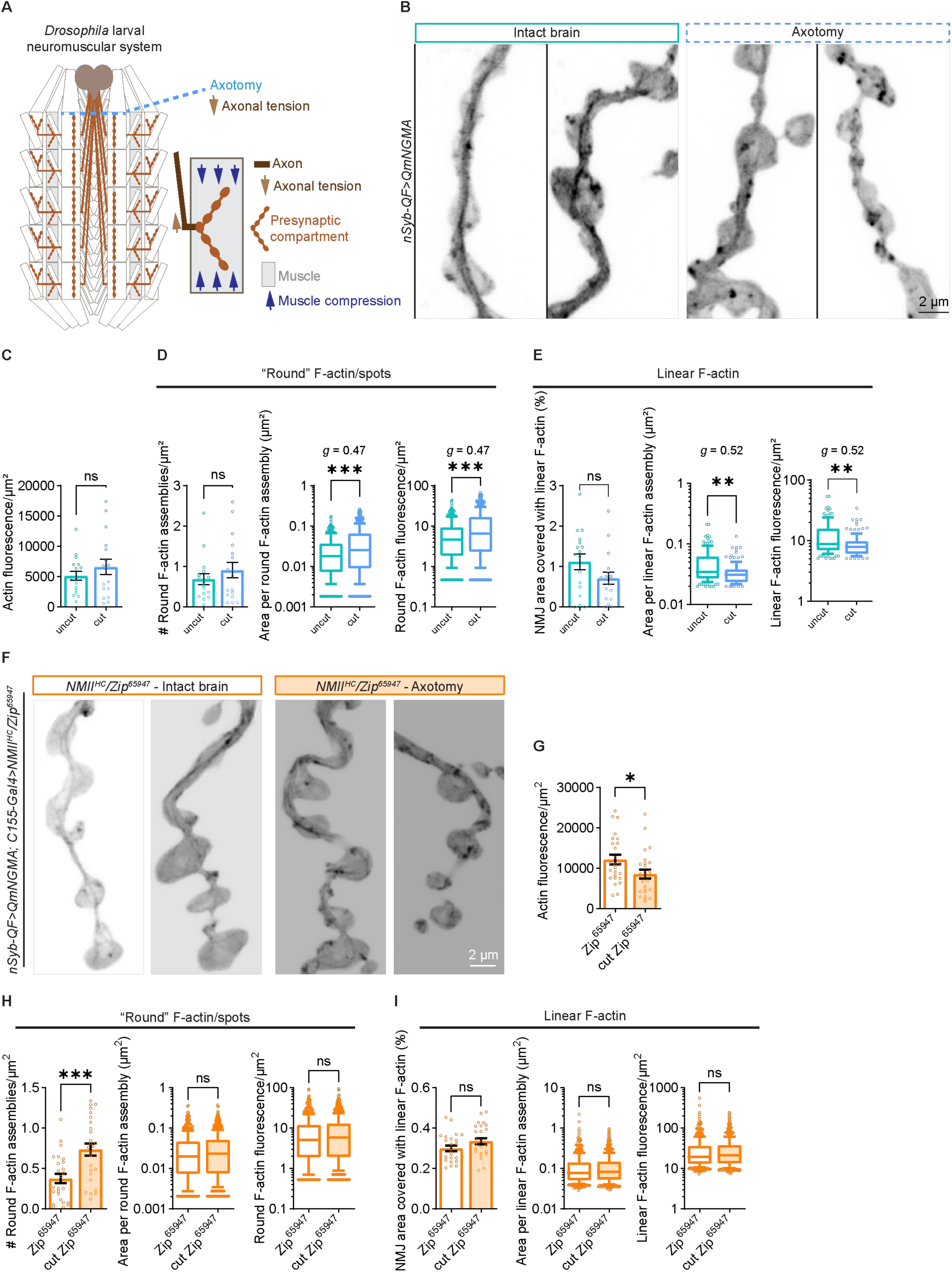
Mechanical severing of larval axons induces changes in presynaptic F-actin. **A**) Diagram of *Drosophila* fillet preparation upon axotomy (light blue dashed line) of all proximal neurons descending from the larval brain. This procedure removes the axonal tension exerted at presynaptic compartments. **B**) mNGMA-labeled presynaptic actin at NMJs from live larvae with intact brains (left) or upon severing of the brain/axotomy (right) (see also **Supp. Movies S10-11**). **C**) Quantification of the fluorescence per µm^2^ for the QmNGMA actin marker in whole NMJs. **D**) Quantifications of the number of round F-actin assemblies per µm^2^, as well as the area and fluorescence of the individual round presynaptic actin structures. **E**) Graphs representing the percentage of NMJ area covered with linear F-actin, as well as the area and fluorescence of the individual assemblies. **F**) mNGMA-labeled presynaptic actin at NMJs from live *NMII^HC^/Zip^65947^* larvae with intact brains (left) or upon severing of the brain/axotomy (right). **G-I**) Analyses of the mNGMA fluorescence, as well as round and linear F-actin assemblies. Box and whiskers graphs were used to represent the results of the area and fluorescence intensity of the individual presynaptic actin structures. Whiskers represent 10^th^ to 90^th^ percentile, while the rest of the data points are shown as individual values. The Y-axis in these graphs represents log10, to capture the broad distribution of the individual values. In bar graphs with linear Y-axis, error bars represent SEM. N in bar graphs – NMJs; N in box and whiskers – individual actin assemblies. ns – not significant, ** p<0.01, and *** p<0.001 upon unpaired, non-parametric Mann-Whitney test. Error bars represent SEM. *g* – Hedges’ *g* represents effect size. Scale bar – 2 µm. **See Table 1 for detailed genotypes and N.**

### Mechanical severing of axons initiates reorganization of Integrin-β receptors

We next examined whether acute loss of axonal tension affects Integrin-β organization by analyzing NMJs 15 minutes after axotomy (**Fig. 8A-B**). While total Integrin-β levels at the α-HRP-delineated- and postsynaptic area at the NMJ remained unchanged (**Fig. 8C**), we observed fragmentation of the linear Integrin-β structures (**Fig. 8D**), similarly to conditions of neuronal NMII depletion. Strikingly, overall muscle Integrin-β levels decreased within this 15-minute window (**Fig. 8C**), demonstrating that muscles rapidly sense and respond to changes in neuronal tension. Since these acute effects parallel our observations from chronic genetic manipulations, they suggest that reorganization of synaptic Integrins reflects a physiological response to changes in neuronal actomyosin rather than an artifact of prolonged genetic manipulation.

**Fig. 8.**
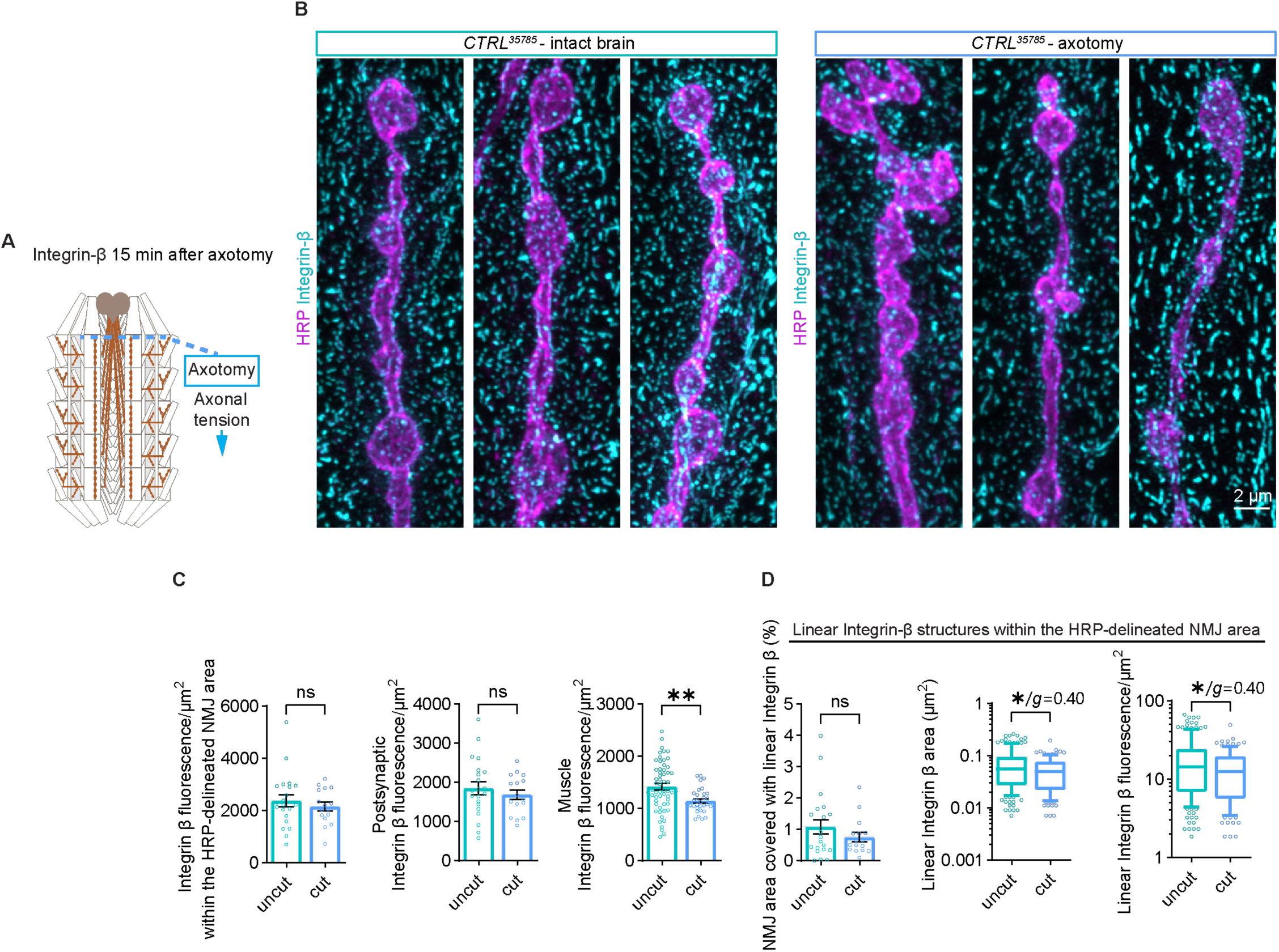
Integrin receptors rearrange within 15 minutes of mechanical severing of larval axons. **A**) Diagrams of *Drosophila* fillet preparation upon axotomy (left, light blue dashed line), a procedure that should decrease the axonal tension exerted at presynaptic compartments. **B**) Immunostaining of Integrin-β (cyan) at NMJs of control larvae (*C155-Gal4>CTRL^35785^*) after 15 minutes with uncut and cut axons. **C**) Integrin-β fluorescence in the HRP-delineated-, postsynaptic-, and muscle ROIs. **D**) Quantifications of the abundance, size and fluorescence of WEKA-segmented Integrin-β structures’ along linear areas. Box and whiskers graphs were used to represent the results of the area and fluorescence intensity of the individual Integrin-β particles. Whiskers represent 10^th^ to 90^th^ percentile, while the rest of the data points are shown as individual values. The Y-axis in these graphs represents log10, to capture the broad distribution of the individual values. In bar graphs with linear Y-axis, error bars represent SEM. N in bar graphs – NMJs or muscle ROIs; N in box and whiskers – individual Integrin-β assemblies. Ns – non-significant, * p<0.05, ** p<0.01, and *** p<0.001 after one-way ANOVA with Kruskal-Wallis multiple comparisons test. *g* – Hedges’ *g* represents effect size. Scale bars – 2 µm. **See Table 1 for detailed genotypes and N.**

## Discussion

Multicellularity requires intracellular molecular machineries that sense and integrate the dynamics of the extracellular environment and of neighboring cells and tissues. This balance becomes especially important when the structural, biochemical, and mechanical properties of interacting cells and tissues are different, posing a challenge to the long-term maintenance of their contact. The neuromuscular junction is an impressive metazoan solution to these mechanical challenges, yet our knowledge on the molecular mechanisms involved remains limited. Our understanding of neuronal mechanobiology comes largely from cell culture studies, which have established essential roles for the actomyosin cytoskeleton and integrin receptors in neuronal development and function (Chighizola et al., 2019). How mechanical forces are generated, sensed, and transmitted within neurons and their surrounding muscles has remained understudied for several reasons. First, neuronal actomyosin structures are submicron in size and visible only with super-resolution microscopy (Berger et al., 2018; Vassilopoulos et al., 2019). Second, the actin cytoskeleton is inherently sensitive to fixation, and using genetically-encoded actin markers to visualize the cytoskeleton live has its own challenges and caveats (Leterrier, 2021). Finally, it is difficult to distinguish cytoskeletal components between closely apposed neurons and muscle cells. In this study, we addressed these challenges using the *Drosophila* larval NMJ as a tissue mechanics model, combining tissue-specific genetic tools, complementary actin markers compatible with live imaging, high-resolution Airyscan microscopy, and machine learning-based image analysis to reveal distinct populations of presynaptic actin structures.

### A novel presynaptic actin core may be analogous to ventral stress fibers

Using these tools, we identified a presynaptic actin assembly that traverses through the center of the NMJ, forming a linear actin core that is a candidate mechanosensitive structure. Our live imaging and FRAP experiments demonstrated that the core is constituted of long-lived, low turnover actin, and is sensitive to the actin depolymerizing drug LatA. The linear nature of the actin core, as well as its lifetime and turnover dynamics, resemble the microns-long ventral stress fibers in adherent cells (Livne and Geiger, 2016). Like ventral stress fibers, which form by merging pre-existing structures rather than *de novo* polymerization (Tojkander et al., 2015), the presynaptic actin core shows no specific enrichment of Arp2/3 or formin nucleators. While both nucleation mechanisms are important at the NMJ - Arp2/3 for endocytosis (Del Signore et al., 2021) and mesh-like structures (Bingham et al., 2023), and formins for presynaptic actin assembly (Bingham et al., 2023) - our FRAP data suggests the linear F-actin structures likely arise from reorganization of existing filaments rather than rapid new nucleation. Like ventral stress fibers, the actin core contained Tropomyosin, an actin stabilizer that facilitates NMII recruitment and enables contractile properties (Tojkander et al., 2011). Interestingly, earlier studies in cultured neurons identified linear F-actin along axons and noted their structural similarity to stress fibers (Gallo, 2006). However, this parallel was dismissed because stress fibers were known to transmit forces to the substrate through focal adhesions - a function that seemed implausible for axonal F-actin at the time. Our whole-animal studies now suggest this initial intuition about stress fiber similarity was correct, as we find the presynaptic actomyosin core does indeed couple to substrate (muscle) through integrin-based adhesions.

The core’s molecular composition and dependence on NMII further supports its similarity to contractile structures. NMII^HC^/Zip partially overlaps with the core, in both smaller puncta and larger assemblies resembling bipolar filaments with doublets spaced ∼260 nm apart (close to the 280-400 nm spacing seen in stress fiber NMII filaments) (Svitkina et al., 1989). Importantly, either decreasing NMII^HC^ levels or altering NMII^LC^ phosphorylation state acutely disrupted presynaptic actin core organization, with dominant negative NMII^LC^ having particularly strong effects on core integrity. This dependence of the actin core on NMII further suggests that it has an antiparallel actin filament organization required for NMII-based contractility. However, we cannot rule out that some filaments may have uniform polarity like yeast actin cables (Moseley and Goode, 2006). Finally, our observation that axotomy disrupted the presynaptic core in a NMII-dependent fashion strongly suggests that it is a mechanoresponsive structure.

### Transsynaptic mechanical signaling via the presynaptic actomyosin core and Integrins

In muscles of larvae with neuronally down-regulated NMII^HC^/Zip or NMII^LC^/Sqh, we observed striking transsynaptic change in the levels of NMII^LC^/Sqh and altered organization of NMII^HC^/Zip. We speculate that the reduction of NMII^LC^/Sqh::GFP could be a result of down-regulation of its expression via transcription regulating pathways, responsive to mechanical triggers (Dupont and Wickström, 2022). Another possibility is changes in proteostasis of NMII in muscles with altered tension, as reported for different proteins undergoing unfolding upon force sensing (Höhfeld et al., 2021). The Zip aggregations we observed might be a physiologically-relevant manner of storing NMII^HC^ in conditions where its activation is temporarily or chronically abrupted (Jordan and Karess, 1997; Royou et al., 2004), especially given that we also observe aggregates in wild-type muscles. Interestingly, accumulations of muscle myosin beneath the sarcolemma and between myofibrils are observed in myosin storage myopathies/myosinopathies (Tajsharghi and Oldfors, 2013), with features indicating defective degradation of misfolded proteins. Thus, it would be interesting to understand whether there is a cellular trigger that changes reversible accumulations of myosins into misfolded pathological aggregates, under sustained mechanical stress or disrupted proteostasis as they unfold upon force sensing (Höhfeld et al., 2021).

In parallel, we detected reduced Integrin levels and altered Integrin organization at NMJs and muscles of larvae with neuronally down-regulated NMII^HC^/Zip. Acute disruption of neuronal tension via axotomy produced similar changes in muscle integrin organization. Since NMII^HC^/Zip downregulation and axotomy both caused disruption of the presynaptic actin core, this suggests that the core can mediate transsynaptic effects on mechanoproteins in the muscle. Our findings suggest that like ventral stress fibers, the presynaptic actomyosin core couples to integrin-based adhesions, perhaps through mechanosensitive linker proteins like talin and vinculin (Goult, Yan and Schwartz, 2018). This inside-out activation of integrins likely promotes both cytoplasmic effector binding that drives adhesion maturation (Schiller and Fässler, 2013) and extracellular conformational changes that facilitate interactions with ligands in the ECM (Shattil, Kim and Ginsberg, 2010) or on the neighboring tissue, i.e., the muscle. Without proper presynaptic actomyosin function, this activation pathway appears disrupted, reducing integrin foci both pre- and postsynaptically while preventing remaining foci from reaching normal size. These local changes at the neuron-muscle interface affect NMII levels near the synapse while sparing distant integrin accumulations at myotendinous junctions. An interesting corollary of these findings is that muscle contractions may reciprocally provide mechanical feedback through outside-in integrin mechanosensing (Chen et al., 2012), creating a feedback loop that contributes to activity-dependent synaptic plasticity at the *Drosophila* NMJ (Dani, Zhu and Broadie, 2014; Lee et al., 2017; Tsai et al., 2012).

Interestingly, we observed both linear and round integrin-β assemblies at the NMJ, likely reflecting distinct organizational states and ligand interactions. This complexity aligns with the diversity of integrin heterodimers and ECM interactions in *Drosophila* (Moreno-Layseca et al., 2019, Humphries, Byron and Humphries, 2006). The varying sizes of integrin foci even in wild-type conditions may reflect different maturation states (Kechagia, Ivaska and Roca-Cusachs, 2019), with recent work linking integrin/talin foci expansion to homeostatic synaptic plasticity at the *Drosophila* NMJ (Orr, Fetter and Davis, 2022). We found that the linear integrin-β populations partially align with the presynaptic actin core, consistent with actin-dependent integrin organization (Swaminathan et al., 2017). Further, both linear and round integrin assemblies are disrupted by neuronal NMII depletion, suggesting the actomyosin core regulates their organization. Together, these findings suggest that synapses employ multiple modes of integrin-based adhesion to maintain mechanical continuity between neurons and muscle.

Another interesting question that arises from our study is how changes in muscle integrin levels arise. The relative amount of active integrins is thought to be controlled via two major mechanisms, both affected by changes in cell membrane tension. One controls the activation of the proteins already at the cell surface, while the other mechanism couples acute mechanical stress to integrin recycling and activation (Lolo et al., 2022). Thus, both mechanisms might be at play in muscles experiencing chronic neuronal actomyosin depletion, or acute changes in tension via axotomy.

### The role of actomyosin in supporting a mechanical continuum in neurons

The axon and presynaptic terminal form a mechanical continuum that must withstand the contractile forces of the muscle and the pulling forces of axons. Previous work has revealed several types of actomyosin organization in axons that may relate to our findings at the NMJ. First, the submembranous cytoskeleton (SMC) consists of periodic actin rings interconnected by spectrin tetramers, which organizes numerous molecules important for neurotransmission, cell adhesion, and molecular transport (Zhou et al., 2022). Additionally, the SMC acts as a mechanical scaffold and tension buffer system for strained axons (Dubey et al., 2020). Pharmacological inhibition or knockdown of NMII suggested that SMC-associated NMII regulates axonal diameter and conduction (Costa et al., 2020; Zhou et al., 2022), as well as bulky cargo transport (Wang et al., 2020) without disrupting periodic SMC organization, suggesting that it primarily regulates radial contractility. By contrast, we observe major rearrangements of presynaptic actin when we manipulate NMII, suggesting that NMII plays a distinct role in maintaining linear actin structures at the NMJ compared to its functions in the SMC.

Several studies have identified longitudinal actin structures in axons that may be more relevant to the core we describe. ’Actin trails’ observed in hippocampal neurons (D’Este et al., 2015; Ganguly et al., 2015), DRG sensory cultured neurons (Unsain et al., 2018), and sensory neurons in *C. elegans* (Sood et al., 2018) are dynamic, bidirectional structures that typically terminate at boutons and may serve as actin delivery vehicles and reservoirs. If generated via mechanisms similar to ventral stress fibers - through reorganization of existing filaments rather than *de novo* assembly - the presynaptic core could serve a dual role as both a mechanosensitive structure and an actin reservoir. Interestingly, axonal actin trails were sensitive to moderate concentrations of SMIFH2, which is a formin inhibitor with off-target effects on NMII activity (Ganguly et al., 2015), raising the possibility that trails are NMII-dependent like the NMJ actin core. Earlier work also identified NMII-dependent longitudinal actin bundles involved in growth cone collapse and axon retraction (Gallo, 2006), while similar structures were recently associated to activity-dependent plasticity in the axon initial segment (Micinski and Hotulainen, 2024). Like these structures, the *Drosophila* NMJ presynaptic actin core depends on NMII activity and responds to mechanical disruption via axotomy. Notably, NMII activity was necessary for severing-induced axon retraction in cultured neurons (Gallo, 2004), which was later associated with rearrangements of axonal F-actin bundles (Phillips et al., 2019). Together with these previous findings, our work suggests neurons employ specialized actomyosin arrays in both axons and presynaptic terminals to maintain mechanical integrity.

Our experiments demonstrate that axotomy of wild-type neurons projecting to the NMJ induces similar fragmentation of the presynaptic actin core to NMII neuronal depletion. In line with this result, we observed that the presynaptic actin core does not fragment further at NMJs with down-regulated NMII. Our findings suggest that there is either a continuous component or a relay that facilitates sensitivity to axotomy. We envisage a spring-like tension sensor that traverses the whole neuron, from the cell body up to the terminals. Our data indicate that a component of this tension sensor at the axon terminal is actomyosin-based. Whether this presynaptic actin core is part of a larger physical mechanical continuum remains to be clarified by more detailed future studies of axonal actin. On the other hand, compartmentalized mechanical strain- mitigating strategies might be employed via local cytoskeletal remodeling (Mutalik et al., 2018). For example, axonal actomyosin can protect against mild mechanical stress through reversible axon beading and dynamic diameter changes. This protection is lost when actomyosin is inactivated, leading to calcium dysregulation and degeneration study (Pan et al., 2024). We propose that the linear actin structures we describe might serve a similar protective function, helping to balance tension along neurons and at the neuron/muscle interface while providing mechanical flexibility to the neuromuscular system.

The presynaptic linear actin core that we described in the current study is an excellent candidate structure to be studied in *Drosophila* models of neuromuscular disorders with underlying actin-related pathomechanisms (Ermanoska et al., 2023). In addition, our findings highlight the potential contribution of muscle dysfunction (myopathy) to neuromuscular disorders traditionally classified as peripheral neuropathies, suggesting a broader pathophysiological framework that includes both neural and muscular components. This is illustrated by patients with an autosomal dominant mutation in MYH14 (encoding the non-muscle myosin NMIIC isoform), who present with complex phenotypes including peripheral and cranial neuropathy combined with myopathy (Choi et al., 2011). Interestingly, NMII-C forms sarcomere-like structures on the periphery of epithelial cells that are precisely aligned with similar structures in the neighbor cell, forming a transcellular contractile network (Ebrahim et al., 2013). Thus, actomyosin rearrangements and mechanotransduction at the interface between different cells, including neuron and muscles, offer potentially valuable therapeutic possibilities.

### Limitations of the Study

This study was constrained by technical challenges inherent to studying actomyosin dynamics at intact synapses in an animal. We employed multiple complementary approaches and controls to address these limitations. A major challenge was visualizing presynaptic F-actin. Phalloidin staining is not compatible with live imaging and, more importantly, small presynaptic actin assemblies are obscured by the prominent muscle F-actin in our *in vivo* system. While we relied on genetically encoded F-actin markers that we previously validated for visualizing presynaptic actin (Del Signore et al., 2021), we cannot completely rule out their effects on actin dynamics in the core (Kumari et al., 2020; Montes-Rodriguez and Kost, 2017; Spracklen et al., 2014). However, our confidence in these tools stems from our previous work showing that three different markers with distinct actin-binding mechanisms produced identical F-actin dynamics, suggesting minimal functional interference (Del Signore et al., 2021). Furthermore, in the current study, we observed both endogenous and tagged Tropomyosin traversing the NMJ in a pattern matching the actin core, providing marker-independent validation of this structure.

Additional technical limitations arose from studying protein localization and mechanical forces. While antibodies against actin-associated proteins helped validate the presynaptic core structure, immunostaining is restricted to fixed samples, and dense postsynaptic signals often obscure presynaptic proteins. We addressed this challenge through detailed WEKA-based image analysis. Next, though GFP-tagged NMII subunits are established tools, their overexpression can perturb NMII biology (Heissler and Sellers, 2015), as evidenced by our finding that Zip::GFP expression alters presynaptic actin organization. We therefore complemented these experiments with endogenous Zip staining. Finally, since our genetic manipulations affect neurons throughout development, some phenotypes may reflect systemic rather than direct effects. While we addressed this through acute manipulation via axotomy, we still lack tools to directly measure or quantitatively manipulate tension within neurons or muscles. Development of tension biosensors and methods for combining precise mechanical manipulation with high-resolution imaging will be crucial for understanding how mechanical forces shape synaptic organization and function.

## Methods

### *Drosophila* maintenance and genetics

*Drosophila melanogaster* stocks were raised on a standard cornmeal medium or molasses formulation for experiments in Fig. 2, Fig. 3F-K, Fig. 5, Fig. 6C-G, Fig. 8, Fig. S3, Fig. S6E-G, Fig. S7C-E, Fig. S9 Crosses were maintained at 25° C, on 12 h light-dark cycles. Detailed genotype information (including sex) of larvae is listed in the source data table and **Table 1**.

### *Drosophila* clones and generation of transgenic flies

The following clones were generated by nucleotide synthesis at VectorBuilder, Inc (Chicago, IL): mNeonGreen::GMA (GMA sequence described in (Bloor and Kiehart, 2001)), mNeonGreen::Tm1 isoform A (NCBI Accession: NP_524360.2), mNeonGreen::Tm1 isoform L (Accession: NP_996216.1), and mScarlet-I::Act5C (NCBI Accession: NP_001284915.1). These constructs were then subcloned into pQUAST-attB (DGRC Stock 1438 ; https://dgrc.bio.indiana.edu//stock/1438; RRID:DGRC_1438). All plasmids were sequence verified (IDG and Plasmidsaurus, Inc). Transgenic flies carrying these constructs were generated at BestGene, Inc., using the PhiC31 system at attP40 or attP2 landing sites (see source data table).

### Dissection procedures for live imaging experiments

All larvae dissected are third instar wandering larvae, coming from at least three different crosses both for control and mutant conditions. The crosses are set with 4-6 virgin female flies to avoid larval overcrowding. Third instar larvae with desired genotypes were held by two pins (anterior and posterior), dissected in Ca^2+^-free HL3.1 (pH 7.2) (Feng, Ueda and Wu, 2004) in a Sylgard polymer-covered plastic Petri dish. Internal organs were carefully removed to preserve muscles, brain and axons from damage. After dissection, one or two larvae were immediately transferred to a microscopic cover slip (ventral side down) in a drop of ∼50 µl. HL3.1. The larvae were carefully sandwiched, using permanent double-sided tape spacers (Scotch 3M ID CBGNHW011141) and a No. 1.5 glass coverslip, and imaged immediately. When two different conditions were tested (e.g. LatA treatment and DMSO, or intact brain and axotomy), we mounted pairs of larvae, and alternated the order in which the two conditions were imaged between pairs. For the Latrunculin A treatment, 1 mM InSolution Latrunculin A (Sigma-Aldrich, 428026) in DMSO was brought to a concentration of 1 µM with HL3.1 dissection medium. As a vehicle control, we used 100% DMSO, similarly dissolved 1000x in HL3.1. Two larvae were dissected as described above, with one undergoing a 30-minute incubation in LatA while the other was incubated in DMSO (in separate Sylgard-coated small Petri dishes). After this pre-incubation, the larvae were immediately mounted in a drop of LatA or DMSO containing HL3.1 on a microscopic slide for imaging, as described above. To visualize the Lifeact::Halo actin marker, dissected larvae were incubated in HL3.1 with 2 µM Janelia Fluor® HaloTag® Ligand-549 for 5 minutes, and directly mounted in ∼50 µl HL3.1 without additional washing. For the axotomy experiment, two larvae were dissected in parallel, then just before mounting, we cut the axons (close to the ventral ganglion) from one larva. Larvae were imaged within 20 minutes of axotomy, on average.

### *Drosophila* immunohistochemistry

Third instar larvae were dissected in HL3.1 (pH 7.2) in a Sylgard-coated dissection dish, fixed for 10 minutes in 4% PFA in HL3.1, washed 3 x 15 minutes in 1xPBS, and permeabilized with 0.1 % PBX (1xPBS with Triton-X) for 3 x 5 minutes. Incubation with the primary antibody was performed either for 2 hours at room temperature, or overnight at 4° C. After a wash (3 x 5 minutes in 0.1 % PBX), the samples were incubated with secondary antibody for 1 hour, followed by a final wash in 0.1 % PBX. Larvae were mounted in ProLong™ Diamond Antifade Mountant (Invitrogen) and cured at room temperature for 48-72 hours before imaging. HRP-Red Rhodamine was used at 1:250. The GFP signal of Sqh::GFP was enhanced by staining with FluoTag®-X4 anti-GFP (NanoTag Biotechnologies, 1:250). Mouse monoclonal anti-Integrin beta PS (DSHB CF.6G11, 1:50) primary, and Alexa Fluor® 488 AffiniPure Goat Anti-Mouse IgG (H+L) (Jackson ImmunoResearch) or CF®488A Donkey Anti-Rabbit IgG (H+L), Highly Cross-Adsorbed Antibody secondary antibodies were used to detect Integrin at the NMJ. Rabbit anti-Tm1 (D. Montell, 1:500, precleared by incubation with dissected, fixed and permeabilized w^1118^ larvae) and Alexa Fluor® 488 AffiniPure Goat Anti-Rabbit IgG (H+L) (Jackson ImmunoResearch) secondary antibodies were used to detect Tm1-A/L at the NMJ. Rabbit anti-Zip (A.M. Sokac, 1:1000, precleared by spinning at 150, 000g for 1,5 h) and CF®488A Donkey Anti-Rabbit IgG (H+L), Highly Cross-Adsorbed secondary antibodies (Biotium, Inc.) were used to detect NMII^HC^/Zip. Rhodamine Red™-X (RRX) AffiniPure Goat Anti-Horseradish Peroxidase (Jackson ImmunoResearch) was used to detect the neuronal membrane of NMJs and axons.

### Microscopy

Imaging was performed on an inverted Zeiss LSM 880 or LSM 900 microscopes with an Airyscan 2 detector, using a 63X oil immersion objective (NA 1.4). Raw images were subjected to 3D Airyscan processing with default settings in Zen Black software. Z-stacks of NMJs in fixed larvae were captured at voxel size (0.044 x 0.044 x 0.185) µm. Z-stacks of NMJs in live larvae were captured with a pixel size 0.044 x 0.044 µm, and Z-step varying from 250-500 µm. For the FRAP experiment, we used NMJs with clearly visible actin core, photobleached them at 100% laser power, and followed the recovery of the signal for 90 seconds with 1 Hz sampling frequency.

### Image analysis

All image analyses were performed in FIJI (Schindelin et al., 2012). Unless otherwise stated, we used maximum intensity projections for all image analyses.

For measuring the fluorescence intensity per µm^2^ of the signals-of-interest in the presynaptic compartment of fixed samples, we segmented and masked NMJs using the HRP signal. The HRP signal was also used to clean parts of images containing axons and debris when analyzing the presynaptic compartment of the NMJ. Before analyzing images with Sqh::GFP and endogenous Zip, we cropped out sections HRP-labeled NMJ in proximity to glia, trachea or axons (all compartments where Sqh::GFP under its own promoter or Zip is enriched in a distinct pattern from the spot-like signal at the NMJ). This way we prevented measuring of irrelevant non-neuronal or non-muscle Sqh::GFP and Zip signal. To obtain a mask the following steps were run on maximum intensity projections: Gaussian Blur with sigma=2 > Auto Threshold method – Huang > Convert to Mask > Erode.

When measuring fluorescence per µm^2^ of dim signals present in both the pre- and post-synaptic compartments of the NMJ (like Sqh::GFP), the background fluorescence per µm^2^ (measured in areas where no signal enrichment was observed) was subtracted from the neuronal fluorescence per µm^2^.

For the remaining image analyses, NMJs were masked using the fluorescently tagged actin or actin marker, or Tm1 (using the HRP masking work flow). To specifically analyze objects in the 1 µm ring surrounding the boutons, we used the “enlarge” function to extend the neuronal mask by 1 µm, and used the XOR function in FIJI to intersect that specific region. In the data presented in **Fig. S2F-G** (LatA treatment of larvae expressing the actin marker QmNGMA in neurons), we generated a WEKA classifier (Arganda-Carreras et al., 2017) that masks the NMJ core, and determined changes in F-actin fluorescence in this area of interest.

#### Segmentation and analysis of individual F-actin assemblies

We built a two-class WEKA classifier by manually annotating round and linear F-actin (**Fig. S1A-C**). The classifier was applied to the first frame of the captured movies and separate probability maps of round and linear F-actin assemblies were obtained as a 32-bit images. In the next step of the image analysis pipeline, we used batch processing to perform particle analysis in Fiji. Round and linear F-actin particles were masked and analyzed separately.

The 32-bit probability map image of linear F-actin after applying the WEKA classifier gave good coverage of predominantly linear structures (**Fig. S1B**). Next, the 32-bit image was transformed to a 16-bit image, which was further used to generate a binary mask. The probability map to binary image transition included more F-actin structures than the linear population. By optimizing the subsequent steps, we designed a pipeline where the binary mask of linear F-actin was further despeckled and skeletonized. Further particle analysis of the skeletonized mask performed with a size cutoff=0.02 µm^2^-Infinity and a circularity cutoff=0.00-0.20, ensuring that the majority of analyzed particles represented linear F-actin structures. The 32- bit probability map image of round F-actin after applying the WEKA classifier similarly gave good coverage of spot-like structures (**Fig. S1C**), while the transition to binary mask did not introduce many linear actin structures. To better segment individual round assemblies, the binary mask underwent despeckling and erosion. All round F-actin particles larger than 0.0018 µm^2^ were included in the analysis. In general, our image analysis pipeline is powerful in providing unbiased and high throughput analysis of specific, individual F-actin assemblies at the synapse. However, we think that it underestimates the size and brightness of these structures as a result of our efforts to segment and analyze them as individual, largely non-overlapping entities.

#### Segmentation and analysis of individual Sqh- and Integrin particles

We built single-class WEKA classifiers that were trained by manually annotating the Sqh or Integrin structures. The classifiers were applied to the maximum intensity projections of the Sqh or Integrin channels, accordingly. The obtained masked Sqh::GFP particles were despeckled and eroded before finally analyzed with no size or circularity filters. To distinguish between linear and round integrin entities, the following cut-offs were applied at the particle analysis step: for spot-like integrin, a size cut-off of 0.0018 µm^2^ (1 pixel^2^) and a circularity cut-off 0.51-1; for linear integrin analysis the segmented entities were analyzed as particles with a size higher than 0.0018 µm^2^ and a circularity between 0-0.5.

#### Segmentation and analysis of individual Zip particles

three-class WEKA classifier (“genuine” spots, aggregates, and background Zip contained in residual axons or trachea) was manually trained to recognize these structures. The classifier was applied to the maximum intensity projections of the Zip channel. The obtained masked Zip particles were analyzed with the following cut-offs: “genuine” Zip spots with a size higher than 0.0018 µm^2^ (1 pixel^2^) and Zip aggregates with a size higher than 0.01 µm^2^. Zip doublets were manually distinguished as paired spot-like structures with each having comparable fluorescent intensity, and thus considered to likely represent formations of bipolar NMII filaments. We then measured the distance between the maxima of the two spots.

Co-localization analysis was performed in Fiji, using a customized macro that segments the whole NMJ as a region of interest (ROI), and then applies the Coloc 2 plugin to individual slices of the two channel z-stacks. Segmentation of the ROI was based on the Lifeact::Halo signal that fills the whole NMJ. The Coloc 2 analysis calculates numerous parameters, including the Pearson’s R value, which we used in our study to score for colocalization of two fluorophores.

### Statistics

We used GraphPad Prism V9 for statistical analyses and generation of graphs. Data were not tested for normality and nonparametric tests were used for analyses. Mann-Whitney tests were used when analyzing two conditions, and one-way ANOVA with, Kruskal-Wallis or Sidek tests with multiple comparison were used to determine the statistical significance of the data from more than two different conditions. To determine the effect size, we measured Hedges’ *g* value, which we also added to graphs. We calculated Hedges’ *g* using the following formula: *g* = M_1_-M_2_/SD_pooled_, where M_1_ – M_2_ is the difference in means, while SD*_pooled_ is the pooled and weighted standard deviation. We used bar graphs or box and whiskers graphs to represent our data. In the bar graphs, the error bars represent standard error of mean (SEM). In the box and whiskers graphs, the whiskers are drawn down to the 10^th^ percentile and up to the 90^th^ percentile. Data points below or above the whiskers are shown as individual points. The line passing through the box represents the median. We used logarithmic scale along the Y axis to display the wide range of values for the area and fluorescence of the different actin assemblies, sqh- and integrin particles in a compact fashion. The FRAP curves in **Fig. 1F** and **Fig. S2D** were constructed after measuring the fluorescence intensity per µm^2^ within the bleached region, as well as in a non-photobleached region to account for the background photobleaching during imaging. The measured values were normalized to the maximum fluorescence intensity value and fitted into a double-exponential curve in GraphPad.

## Acknowledgements

We thank the *Drosophila* Genomics Resource Center (NIH Grant 2P40OD010949); Bloomington *Drosophila* Stock Center (Indiana University, Bloomington, IN, NIH 595 P40OD018537); Guy Tanentzapf (UBC) for *PUbi-mys.YFP*; and Denise Montell (UCSB) for the Tm1 antibody; Adam Martin (MIT) for UAS-Zip::GFP flies; Anna Marie Sokac (UIUC) for Zip antibody; Bob Asselbergh and Simona Manzella (VIB/CMN/UAntwerp) for help with Zeiss LSM 900 microscopy, and to the Brandeis University Light Microscopy Core Facility, RRID:SCR_025892.

## Funding sources

B.E. is currently supported by Marie Sklodowska Curie Postdoctoral fellowship 101107344; J.B. is supported by the Research Fund - Flanders (FWO) via a Senior Clinical Researcher mandate 1805021N. This work was supported by kBOF FFB240038 to B.E.; Research Fund - Flanders (FWO) Research Grant (G071723N) and EU Horizon 2020 program/Solve-RD under grant agreement 779257 to J.B.; NINDS grant R01 NS116375 to A.A.R.

J.B. is member of the European Reference Network for Rare Neuromuscular Diseases (ERN EURO-NMD). J.B. and B.E. are members of the µNEURO Research Centre of Excellence of the University of Antwerp.

## Competing interests

The authors declare no competing interests. J.B. has received *ad hoc* consultancy compensation for activities with Novartis, Sanofi, CSL Behring, Alnylam, Roche, ARGENX and Amylyx.

## Author contributions

Conceptualization: B.E., A.A.R. Methodology: B.E., A.A.R. Investigation: B.E. Visualization: B.E. Funding acquisition: B.E., J.B. and A.A.R. Project administration: A.A.R. Supervision: J.B. and A.A.R. Writing – original draft: B.E. Writing—review & editing: B.E., J.B. and A.A.R.

## Notes

### Competing Interest Statement

The authors have declared no competing interest.

### Summary of Updates

We have provided new independent data supporting the conclusion that the presynaptic actin core is actually actomyosin, validated transsynaptic changes of NMII, included additional negative controls, and discussed in more detail potentially analogous axonal structures. These changes are represented in ten new data sets in the revised manuscript (five new figures: Fig. 2, 5, 8, S3, and S9; five additional data sets: Fig. 3F-K, Fig. 6C-G, Fig. 7F-I, Fig. S6E-G, and Fig. 7C-E)

## References

Abouelezz, A., H. Stefen, M. Segerstråle, D. Micinski, R. Minkeviciene, L. Lahti, E.C. Hardeman, P.W. Gunning, C.C. Hoogenraad, T. Taira, T. Fath, and P. Hotulainen. 2020. Tropomyosin Tpm3.1 Is Required to Maintain the Structure and Function of the Axon Initial Segment. iScience. 23:101053.

Akamatsu, M., R. Vasan, D. Serwas, M.A. Ferrin, P. Rangamani, and D.G. Drubin. 2020. Principles of self-organization and load adaptation by the actin cytoskeleton during clathrin-mediated endocytosis. eLife. 9:e49840.

Alioto, S.L., M.V. Garabedian, D.R. Bellavance, and B.L. Goode. 2016. Tropomyosin and Profilin Cooperate to Promote Formin-Mediated Actin Nucleation and Drive Yeast Actin Cable Assembly. Curr Biol. 26:3230–3237.

Arganda-Carreras, I., V. Kaynig, C. Rueden, K.W. Eliceiri, J. Schindelin, A. Cardona, and H. Sebastian Seung. 2017. Trainable Weka Segmentation: a machine learning tool for microscopy pixel classification. Bioinformatics. 33:2424–2426.

Ataman, B., J. Ashley, M. Gorczyca, P. Ramachandran, W. Fouquet, S.J. Sigrist, and V. Budnik. 2008. Rapid activity-dependent modifications in synaptic structure and function require bidirectional Wnt signaling. Neuron. 57:705–718.

Ataman, B., V. Budnik, and U. Thomas. 2006. Scaffolding proteins at the *Drosophila* neuromuscular junction. Int Rev Neurobiol. 75:181–216.

Berger, S.L., A. Leo-Macias, S. Yuen, L. Khatri, S. Pfennig, Y. Zhang, E. Agullo-Pascual, G. Caillol, M.S. Zhu, E. Rothenberg, C.V. Melendez-Vasquez, M. Delmar, C. Leterrier, and J.L. Salzer. 2018. Localized Myosin II Activity Regulates Assembly and Plasticity of the Axon Initial Segment. Neuron. 97:555–570.e556.

Beumer, K.J., J. Rohrbough, A. Prokop, and K. Broadie. 1999. A role for PS integrins in morphological growth and synaptic function at the postembryonic neuromuscular junction of *Drosophila*. Development. 126:5833–5846.

Bindels, D.S., L. Haarbosch, L. van Weeren, M. Postma, K.E. Wiese, M. Mastop, S. Aumonier, G. Gotthard, A. Royant, M.A. Hink, and T.W. Gadella, Jr. 2017. mScarlet: a bright monomeric red fluorescent protein for cellular imaging. Nat Methods. 14:53–56.

Bingham, D., C.E. Jakobs, F. Wernert, F. Boroni-Rueda, N. Jullien, E.M. Schentarra, K. Friedl, J. Da Costa Moura, D.M. van Bommel, G. Caillol, Y. Ogawa, M.J. Papandréou, and C. Leterrier. 2023. Presynapses contain distinct actin nanostructures. J Cell Biol. 222.

Bloor, J.W., and D.P. Kiehart. 2001. zipper Nonmuscle myosin-II functions downstream of PS2 integrin in *Drosophila* myogenesis and is necessary for myofibril formation. Dev Biol. 239:215–228.

Brand, A.H., and N. Perrimon. 1993. Targeted gene expression as a means of altering cell fates and generating dominant phenotypes. Development. 118:401–415.

Brettle, M., S. Patel, and T. Fath. 2016. Tropomyosins in the healthy and diseased nervous system. Brain Research Bulletin. 126:311–323.

Chastney, M.R., J.R.W. Conway, and J. Ivaska. 2021. Integrin adhesion complexes. Curr Biol. 31:R536–r542.

Chen, W., J. Lou, E.A. Evans, and C. Zhu. 2012. Observing force-regulated conformational changes and ligand dissociation from a single integrin on cells. J Cell Biol. 199:497–512.

Chighizola, M., T. Dini, C. Lenardi, P. Milani, A. Podestà, and C. Schulte. 2019. Mechanotransduction in neuronal cell development and functioning. Biophys Rev. 11:701–720.

Cho, A., M. Kato, T. Whitwam, J.H. Kim, and D.J. Montell. 2016. An Atypical Tropomyosin in *Drosophila* with Intermediate Filament-like Properties. Cell Rep. 16:928–938.

Choi, B.O., S.H. Kang, Y.S. Hyun, S. Kanwal, S.W. Park, H. Koo, S.B. Kim, Y.C. Choi, J.H. Yoo, J.W. Kim, K.D. Park, K.G. Choi, S.J. Kim, S. Züchner, and K.W. Chung. 2011. A complex phenotype of peripheral neuropathy, myopathy, hoarseness, and hearing loss is linked to an autosomal dominant mutation in MYH14. Hum Mutat. 32:669–677.

Costa, A.R., S.C. Sousa, R. Pinto-Costa, J.C. Mateus, C.D. Lopes, A.C. Costa, D. Rosa, D. Machado, L. Pajuelo, X. Wang, F.Q. Zhou, A.J. Pereira, P. Sampaio, B.Y. Rubinstein, I. Mendes Pinto, M. Lampe, P. Aguiar, and M.M. Sousa. 2020. The membrane periodic skeleton is an actomyosin network that regulates axonal diameter and conduction. Elife. 9.

D’Este, E., D. Kamin, F. Göttfert, A. El-Hady, and S.W. Hell. 2015. STED nanoscopy reveals the ubiquity of subcortical cytoskeleton periodicity in living neurons. Cell Rep. 10:1246–1251.

Dani, N., H. Zhu, and K. Broadie. 2014. Two protein N-acetylgalactosaminyl transferases regulate synaptic plasticity by activity-dependent regulation of integrin signaling. J Neurosci. 34:13047–13065.

Del Signore, S.J., C.F. Kelley, E.M. Messelaar, T. Lemos, M.F. Marchan, B. Ermanoska, M. Mund, T.G. Fai, M. Kaksonen, and A.A. Rodal. 2021. An autoinhibitory clamp of actin assembly constrains and directs synaptic endocytosis. eLife. 10:e69597.

Dubey, S., N. Bhembre, S. Bodas, S. Veer, A. Ghose, A. Callan-Jones, and P. Pullarkat. 2020. The axonal actin-spectrin lattice acts as a tension buffering shock absorber. eLife. 9:e51772.

Dupont, S., and S.A. Wickström. 2022. Mechanical regulation of chromatin and transcription. Nat Rev Genet. 23:624–643.

Ebrahim, S., T. Fujita, B.A. Millis, E. Kozin, X. Ma, S. Kawamoto, M.A. Baird, M. Davidson, S. Yonemura, Y. Hisa, M.A. Conti, R.S. Adelstein, H. Sakaguchi, and B. Kachar. 2013. NMII forms a contractile transcellular sarcomeric network to regulate apical cell junctions and tissue geometry. Curr Biol. 23:731–736.

Ermanoska, B., B. Asselbergh, L. Morant, M.L. Petrovic-Erfurth, S. Hosseinibarkooie, R. Leitão-Gonçalves, L. Almeida-Souza, S. Bervoets, L. Sun, L. Lee, D. Atkinson, A. Khanghahi, I. Tournev, P. Callaerts, P. Verstreken, X.L. Yang, B. Wirth, A.A. Rodal, V. Timmerman, B.L. Goode, T.A. Godenschwege, and A. Jordanova. 2023. Tyrosyl-tRNA synthetase has a noncanonical function in actin bundling. Nat Commun. 14:999.

Feng, Y., A. Ueda, and C.F. Wu. 2004. A modified minimal hemolymph-like solution, HL3.1, for physiological recordings at the neuromuscular junctions of normal and mutant *Drosophila* larvae. J Neurogenet. 18:377-402.

Fernandes, A.R., J.P. Martins, E.R. Gomes, C.S. Mendes, and R.O. Teodoro. 2023. *Drosophila* motor neuron boutons remodel through membrane blebbing coupled with muscle contraction. Nat Commun. 14:3352.

Gallo, G. 2004. Myosin II activity is required for severing-induced axon retraction in vitro. Exp Neurol. 189:112–121.

Gallo, G. 2006. RhoA-kinase coordinates F-actin organization and myosin II activity during semaphorin-3A-induced axon retraction. J Cell Sci. 119:3413–3423.

Ganguly, A., Y. Tang, L. Wang, K. Ladt, J. Loi, B. Dargent, C. Leterrier, and S. Roy. 2015. A dynamic formin-dependent deep F-actin network in axons. J Cell Biol. 210:401–417.

Garrido-Casado, M., G. Asensio-Juárez, V.C. Talayero, and M. Vicente-Manzanares. 2024. Engines of change: Nonmuscle myosin II in mechanobiology. Curr Opin Cell Biol. 87:102344.

Goult, B.T., J. Yan, and M.A. Schwartz. 2018. Talin as a mechanosensitive signaling hub. J Cell Biol. 217:3776–3784.

He, J., R. Zhou, Z. Wu, M.A. Carrasco, P.T. Kurshan, J.E. Farley, D.J. Simon, G. Wang, B. Han, J. Hao, E. Heller, M.R. Freeman, K. Shen, T. Maniatis, M. Tessier-Lavigne, and X. Zhuang. 2016. Prevalent presence of periodic actin-spectrin-based membrane skeleton in a broad range of neuronal cell types and animal species. Proc Natl Acad Sci U S A. 113:6029–6034.

Heissler, S.M., and J.R. Sellers. 2015. Four things to know about myosin light chains as reporters for non-muscle myosin-2 dynamics in live cells. Cytoskeleton (Hoboken). 72:65–70.

Höhfeld, J., T. Benzing, W. Bloch, D.O. Fürst, S. Gehlert, M. Hesse, B. Hoffmann, T. Hoppe, P.F. Huesgen, M. Köhn, W. Kolanus, R. Merkel, C.M. Niessen, W. Pokrzywa, M.M. Rinschen, D. Wachten, and B. Warscheid. 2021. Maintaining proteostasis under mechanical stress. EMBO Rep. 22:e52507.

Humphries, J.D., A. Byron, and M.J. Humphries. 2006. Integrin ligands at a glance. J Cell Sci. 119:3901–3903.

Iyer, S.R., S.B. Shah, and R.M. Lovering. 2021. The Neuromuscular Junction: Roles in Aging and Neuromuscular Disease. Int J Mol Sci. 22.

Jordan, P., and R. Karess. 1997. Myosin light chain-activating phosphorylation sites are required for oogenesis in *Drosophila*. J Cell Biol. 139:1805–1819.

Kechagia, J.Z., J. Ivaska, and P. Roca-Cusachs. 2019. Integrins as biomechanical sensors of the microenvironment. Nat Rev Mol Cell Biol. 20:457–473.

Kumari, A., S. Kesarwani, M.G. Javoor, K.R. Vinothkumar, and M. Sirajuddin. 2020. Structural insights into actin filament recognition by commonly used cellular actin markers. Embo j. 39:e104006.

Lee, J.Y., J. Geng, J. Lee, A.R. Wang, and K.T. Chang. 2017. Activity-Induced Synaptic Structural Modifications by an Activator of Integrin Signaling at the *Drosophila* Neuromuscular Junction. J Neurosci. 37:3246–3263.

Leterrier, C. 2021. A Pictorial History of the Neuronal Cytoskeleton. J Neurosci. 41:11–27.

Livne, A., and B. Geiger. 2016. The inner workings of stress fibers - from contractile machinery to focal adhesions and back. J Cell Sci. 129:1293–1304.

Lolo, F.N., D.M. Pavón, A. Grande-García, A. Elosegui-Artola, V.I. Segatori, S. Sánchez, X. Trepat, P. Roca-Cusachs, and M.A. Del Pozo. 2022. Caveolae couple mechanical stress to integrin recycling and activation. Elife. 11.

Michelot, A., and D.G. Drubin. 2011. Building distinct actin filament networks in a common cytoplasm. Curr Biol. 21:R560–569.

Micinski, D., and P. Hotulainen. 2024. Actin polymerization and longitudinal actin fibers in axon initial segment plasticity. Front Mol Neurosci. 17:1376997.

Montes-Rodriguez, A., and B. Kost. 2017. Direct Comparison of the Performance of Commonly Employed In Vivo F-actin Markers (Lifeact-YFP, YFP-mTn and YFP-FABD2) in Tobacco Pollen Tubes. Front Plant Sci. 8:1349.

Moreno-Layseca, P., J. Icha, H. Hamidi, and J. Ivaska. 2019. Integrin trafficking in cells and tissues. Nat Cell Biol. 21:122–132.

Moseley, J.B., and B.L. Goode. 2006. The yeast actin cytoskeleton: from cellular function to biochemical mechanism. Microbiol Mol Biol Rev. 70:605–645.

Murrell, M., P.W. Oakes, M. Lenz, and M.L. Gardel. 2015. Forcing cells into shape: the mechanics of actomyosin contractility. Nat Rev Mol Cell Biol. 16:486–498.

Mutalik, S.P., J. Joseph, P.A. Pullarkat, and A. Ghose. 2018. Cytoskeletal Mechanisms of Axonal Contractility. Biophys J. 115:713–724.

Orr, B.O., R.D. Fetter, and G.W. Davis. 2022. Activation and expansion of presynaptic signaling foci drives presynaptic homeostatic plasticity. Neuron. 110:3743–3759.e3746.

Pan, X., Y. Hu, G. Lei, Y. Wei, J. Li, T. Luan, Y. Zhang, Y. Chu, Y. Feng, W. Zhan, C. Zhao, F.A. Meunier, Y. Liu, Y. Li, and T. Wang. 2024. Actomyosin-II protects axons from degeneration induced by mild mechanical stress. J Cell Biol. 223.

Park, Y.K., and Y. Goda. 2016. Integrins in synapse regulation. Nat Rev Neurosci. 17:745–756.

Phillips, J.K., S.A. Sherman, K.Y. Cotton, J.M. Heddleston, A.B. Taylor, and J.D. Finan. 2019. Characterization of neurite dystrophy after trauma by high speed structured illumination microscopy and lattice light sheet microscopy. J Neurosci Methods. 312:154–161.

Piccioli, Z.D., and J.T. Littleton. 2014. Retrograde BMP signaling modulates rapid activity-dependent synaptic growth via presynaptic LIM kinase regulation of cofilin. J Neurosci. 34:4371–4381.

Pollard, T.D. 2016. Actin and Actin-Binding Proteins. Cold Spring Harb Perspect Biol. 8.

Potter, C.J., B. Tasic, E.V. Russler, L. Liang, and L. Luo. 2010. The Q system: a repressible binary system for transgene expression, lineage tracing, and mosaic analysis. Cell. 141:536–548.

Qu, Y., I. Hahn, S.E. Webb, S.P. Pearce, and A. Prokop. 2017. Periodic actin structures in neuronal axons are required to maintain microtubules. Mol Biol Cell. 28:296–308.

Quintanilla, M.A., J.A. Hammer, and J.R. Beach. 2023. Non-muscle myosin 2 at a glance. J Cell Sci. 136.

Riedl, J., A.H. Crevenna, K. Kessenbrock, J.H. Yu, D. Neukirchen, M. Bista, F. Bradke, D. Jenne, T.A. Holak, Z. Werb, M. Sixt, and R. Wedlich-Soldner. 2008. Lifeact: a versatile marker to visualize F-actin. Nature Methods. 5:605–607.

Royou, A., C. Field, J.C. Sisson, W. Sullivan, and R. Karess. 2004. Reassessing the role and dynamics of nonmuscle myosin II during furrow formation in early *Drosophila* embryos. Mol Biol Cell. 15:838–850.

Russell, R.J., A.Y. Grubbs, S.P. Mangroo, S.E. Nakasone, R.B. Dickinson, and T.P. Lele. 2011. Sarcomere length fluctuations and flow in capillary endothelial cells. Cytoskeleton (Hoboken). 68:150–156.

Schiller, H.B., and R. Fässler. 2013. Mechanosensitivity and compositional dynamics of cell-matrix adhesions. EMBO Rep. 14:509–519.

Schindelin, J., I. Arganda-Carreras, E. Frise, V. Kaynig, M. Longair, T. Pietzsch, S. Preibisch, C. Rueden, S. Saalfeld, B. Schmid, J.-Y. Tinevez, D.J. White, V. Hartenstein, K. Eliceiri, P. Tomancak, and A. Cardona. 2012. Fiji: an open-source platform for biological-image analysis. Nature Methods. 9:676-682.

Seabrooke, S., X. Qiu, and B.A. Stewart. 2010. Nonmuscle Myosin II helps regulate synaptic vesicle mobility at the *Drosophila* neuromuscular junction. BMC Neurosci. 11:37.

Seabrooke, S., and B.A. Stewart. 2011. Synaptic transmission and plasticity are modulated by nonmuscle myosin II at the neuromuscular junction of *Drosophila*. J Neurophysiol. 105:1966–1976.

Sellers, J.R. 1991. Regulation of cytoplasmic and smooth muscle myosin. Curr Opin Cell Biol. 3:98–104.

Sellers, J.R., and S.M. Heissler. 2019. Nonmuscle myosin-2 isoforms. Curr Biol. 29:R275–r278.

Shaner, N.C., G.G. Lambert, A. Chammas, Y. Ni, P.J. Cranfill, M.A. Baird, B.R. Sell, J.R. Allen, R.N. Day, M. Israelsson, M.W. Davidson, and J. Wang. 2013. A bright monomeric green fluorescent protein derived from *Branchiostoma lanceolatum*. Nature Methods. 10:407–409.

Shattil, S.J., C. Kim, and M.H. Ginsberg. 2010. The final steps of integrin activation: the end game. Nat Rev Mol Cell Biol. 11:288–300.

Siechen, S., S. Yang, A. Chiba, and T. Saif. 2009. Mechanical tension contributes to clustering of neurotransmitter vesicles at presynaptic terminals. Proc Natl Acad Sci U S A. 106:12611–12616.

Sokac, A.M., and E. Wieschaus. 2008. Local Actin-Dependent Endocytosis Is Zygotically Controlled to Initiate *Drosophila* Cellularization. Developmental Cell. 14:775–786.

Sood, P., K. Murthy, V. Kumar, M.L. Nonet, G.I. Menon, and S.P. Koushika. 2018. Cargo crowding at actin-rich regions along axons causes local traffic jams. Traffic. 19:166–181.

Spracklen, A.J., T.N. Fagan, K.E. Lovander, and T.L. Tootle. 2014. The pros and cons of common actin labeling tools for visualizing actin dynamics during *Drosophila* oogenesis. Dev Biol. 393:209–226.

Svitkina, T.M., I.G. Surguchova, A.B. Verkhovsky, V.I. Gelfand, M. Moeremans, and J. De Mey. 1989. Direct visualization of bipolar myosin filaments in stress fibers of cultured fibroblasts. Cell Motil Cytoskeleton. 12:150–156.

Swaminathan, V., J.M. Kalappurakkal, S.B. Mehta, P. Nordenfelt, T.I. Moore, N. Koga, D.A. Baker, R. Oldenbourg, T. Tani, S. Mayor, T.A. Springer, and C.M. Waterman. 2017. Actin retrograde flow actively aligns and orients ligand-engaged integrins in focal adhesions. Proc Natl Acad Sci U S A. 114:10648–10653.

Tajsharghi, H., and A. Oldfors. 2013. Myosinopathies: pathology and mechanisms. Acta Neuropathol. 125:3–18.

Tojkander, S., G. Gateva, A. Husain, R. Krishnan, and P. Lappalainen. 2015. Generation of contractile actomyosin bundles depends on mechanosensitive actin filament assembly and disassembly. eLife. 4:e06126.

Tojkander, S., G. Gateva, G. Schevzov, P. Hotulainen, P. Naumanen, C. Martin, Peter W. Gunning, and P. Lappalainen. 2011. A Molecular Pathway for Myosin II Recruitment to Stress Fibers. Current Biology. 21:539–550.

Tsai, P.I., M. Wang, H.H. Kao, Y.J. Cheng, Y.J. Lin, R.H. Chen, and C.T. Chien. 2012. Activity-dependent retrograde laminin A signaling regulates synapse growth at *Drosophila* neuromuscular junctions. Proc Natl Acad Sci U S A. 109:17699–17704.

Tyler, W.J. 2012. The mechanobiology of brain function. Nat Rev Neurosci. 13:867–878.

Unsain, N., M.D. Bordenave, G.F. Martinez, S. Jalil, C. von Bilderling, F.M. Barabas, L.A. Masullo, A.D. Johnstone, P.A. Barker, M. Bisbal, F.D. Stefani, and A.O. Cáceres. 2018. Remodeling of the Actin/Spectrin Membrane-associated Periodic Skeleton, Growth Cone Collapse and F-Actin Decrease during Axonal Degeneration. Sci Rep. 8:3007.

Vasin, A., N. Sabeva, C. Torres, S. Phan, E.A. Bushong, M.H. Ellisman, and M. Bykhovskaia. 2019. Two Pathways for the Activity-Dependent Growth and Differentiation of Synaptic Boutons in *Drosophila*. eNeuro. 6.

Vasquez, C.G., M. Tworoger, and A.C. Martin. 2014. Dynamic myosin phosphorylation regulates contractile pulses and tissue integrity during epithelial morphogenesis. J Cell Biol. 206:435–450.

Vassilopoulos, S., S. Gibaud, A. Jimenez, G. Caillol, and C. Leterrier. 2019. Ultrastructure of the axonal periodic scaffold reveals a braid-like organization of actin rings. Nature Communications. 10:5803.

Vicente-Manzanares, M., X. Ma, R.S. Adelstein, and A.R. Horwitz. 2009. Non-muscle myosin II takes centre stage in cell adhesion and migration. Nat Rev Mol Cell Biol. 10:778–790.

Wang, Q., T.H. Han, P. Nguyen, M. Jarnik, and M. Serpe. 2018. Tenectin recruits integrin to stabilize bouton architecture and regulate vesicle release at the *Drosophila* neuromuscular junction. Elife. 7.

Wang, T., W. Li, S. Martin, A. Papadopulos, M. Joensuu, C. Liu, A. Jiang, G. Shamsollahi, R. Amor, V. Lanoue, P. Padmanabhan, and F.A. Meunier. 2020. Radial contractility of actomyosin rings facilitates axonal trafficking and structural stability. J Cell Biol. 219.

Wen, P.J., S. Grenklo, G. Arpino, X. Tan, H.S. Liao, J. Heureaux, S.Y. Peng, H.C. Chiang, E. Hamid, W.D. Zhao, W. Shin, T. Näreoja, E. Evergren, Y. Jin, R. Karlsson, S.N. Ebert, A. Jin, A.P. Liu, O. Shupliakov, and L.G. Wu. 2016. Actin dynamics provides membrane tension to merge fusing vesicles into the plasma membrane. Nat Commun. 7:12604.

Xu, K., G. Zhong, and X. Zhuang. 2013. Actin, spectrin, and associated proteins form a periodic cytoskeletal structure in axons. Science. 339:452–456.

Yuan, L., M.J. Fairchild, A.D. Perkins, and G. Tanentzapf. 2010. Analysis of integrin turnover in fly myotendinous junctions. J Cell Sci. 123:939–946.

Zhou, R., B. Han, R. Nowak, Y. Lu, E. Heller, C. Xia, A.H. Chishti, V.M. Fowler, and X. Zhuang. 2022. Proteomic and functional analyses of the periodic membrane skeleton in neurons. Nature Communications. 13:3196.

