## Supplementary Figures, Tables and Legends for "Non-muscle myosin II regulates presynaptic actin assemblies and neuronal mechanobiology in *Drosophila*"

Supplementary information

**This supplementary information file includes:**

Supplementary Figures 1 to 9

References for the supplementary information

Resource data table

Table 1. Experimental genotype and statistical reporting

Legends for Supplementary Movies 1-11

**Other supplementary materials for this manuscript include the following:**
Supplementary Movies 1-11

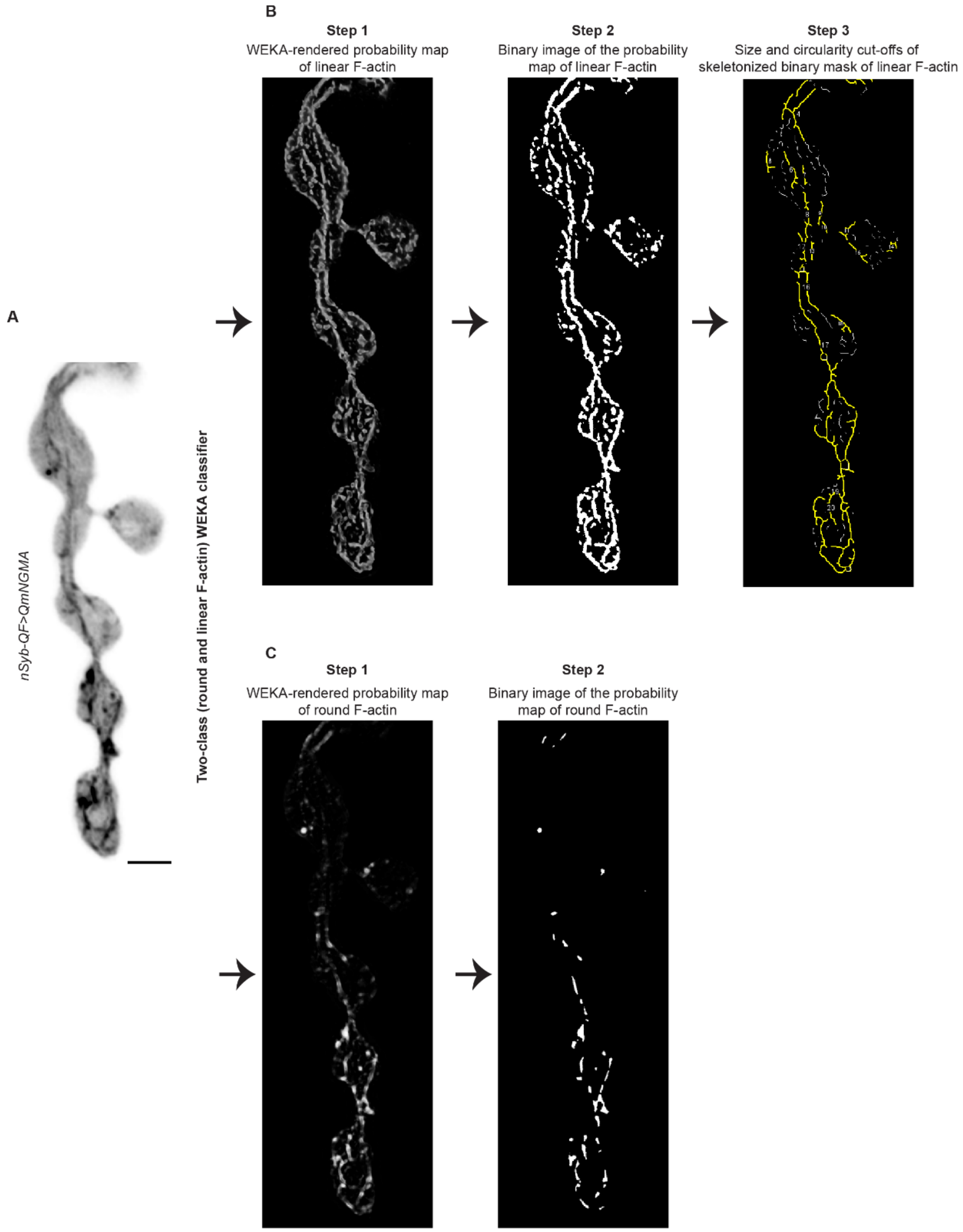

**Supplemental figure 1. Segmentation and analysis of round and linear F-actin assemblies. A**) Example of image analysis pipeline: Inverted contrast image of mNGMA-labeled presynaptic actin in live dissected larvae. A two-class (round and linear F-actin) WEKA classifier was applied. **B**) The product of the WEKA-based segmentation of linear actin is a 32-bit probability map image, which is converted to 16-bit image to make a binary mask (8-bit). The binary mask was further skeletonized, and a circularity (0-0.2) and size (0.02 $\mu m^2$-Infinity) cutoffs were implemented at the particle analysis step to distinguish *bona fide* linear F-actin. **C**) The 32-bit probability map image of round F-actin was similarly converted to 16-bit image, which was used to make a binary mask of the structures. The binary mask was used in particle analysis of structures with size higher than 0.0018 $\mu m^{2,}$ i.e. 1 pixel$^2$. See additional description in Methods.

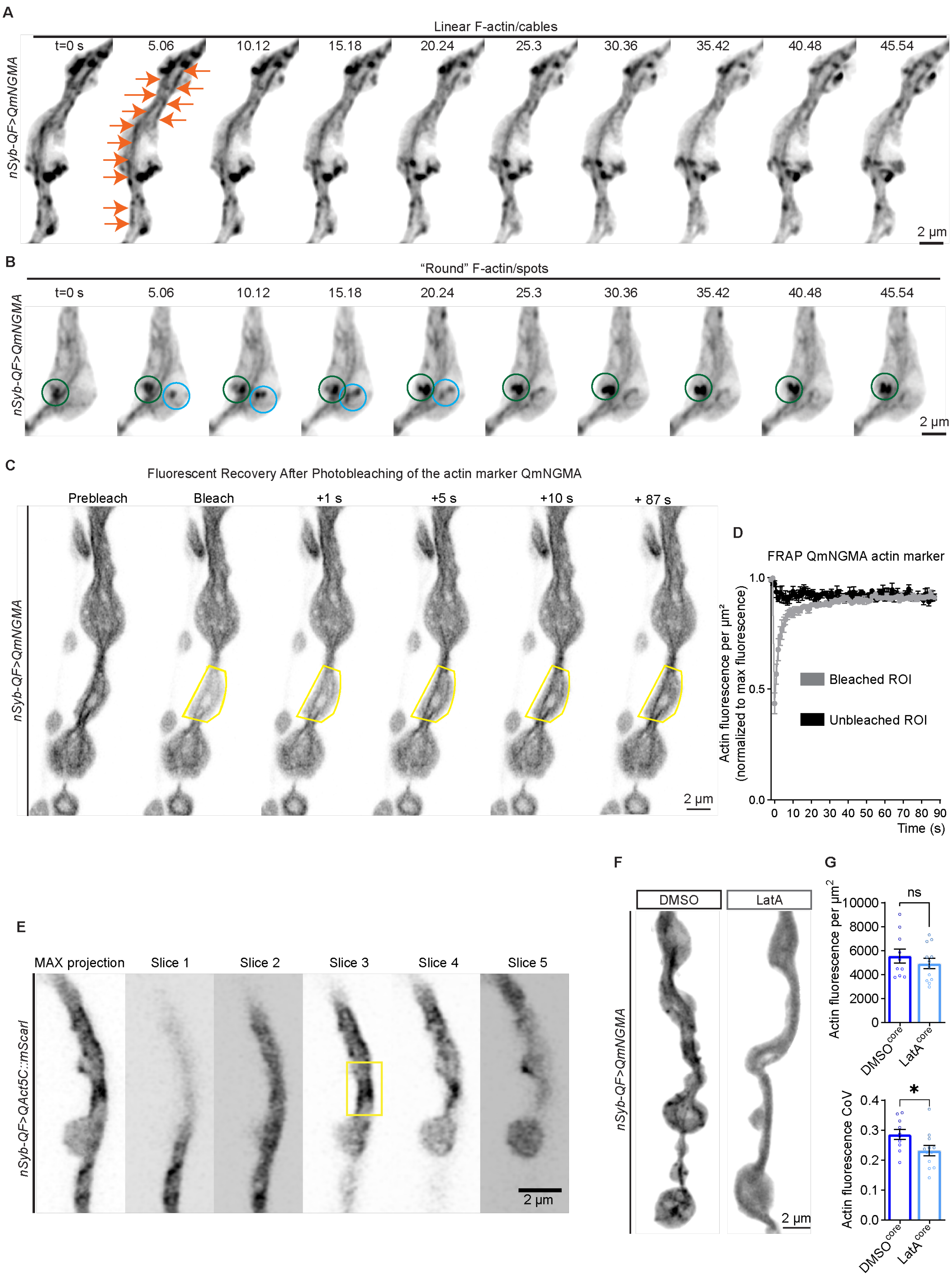
A
Linear F-actin/cables
t=0 s
5.06
10.12
15.18
20.24
25.3
30.36
35.42
40.48
45.54
nSyb-QF>QmNGMA
2 µm
B
"Round" F-actin/spots
C
Fluorescent Recovery After Photobleaching of the actin marker QmNGMA
Prebleach
Bleach
+1 s
+5 s
+10 s
+ 87 s
D
FRAP QmNGMA actin marker
Actin fluorescence per µm²
(normalized to max fluorescence)
Bleached ROI
Unbleached ROI
Time (s)
E
MAX projection
Slice 1
Slice 2
Slice 3
Slice 4
Slice 5
nSyb-QF>QAct5C::mScarl
F
DMSO
LatA
G
ns
Actin fluorescence per µm²
DMSOcore
LatAcore
*
Actin fluorescence CoV

**Supplemental Figure 2. Presynaptic actin cytoskeleton at the larval *Drosophila* neuromuscular junction. A**) Time-lapse insets (from Supp. movie 1) showing the structure and lifetime of linear F-actin/cable-like structures at the NMJ. **B**) Time-lapse insets of round presynaptic F-actin assemblies (encircled) with different lifetimes (green circle around a longer-lived assembly, light-blue circle around a short-lived, possibly endocytic event) visualized via the actin marker QmNGMA at the NMJ. **C**) Time-lapse of fluorescence recovery after photobleaching (FRAP) of the neuronally expressed actin marker QmNGMA at a bouton containing linear F-actin assemblies. The bleached bouton is outlined in yellow. **D**) FRAP quantification of linear F-actin-containing region showing recovery of QmNGMA (1 frame/sec, n=3). **E**) Inverted contrast montage of the maximum intensity projection, along with individual Z-stacks of an NMJ from a larvae expressing mScarlet-I::Act5C neuronally (*nSybQF>QAct5C::mScarI*). The yellow rectangle marks a typical region of the NMJ selected for FRAP analysis of the presynaptic actin core. **F**) Presynaptic actin assemblies are sensitive to a treatment with the actin depolymerization-inducing drug Latrunculin A (LatA). **G**) Quantifications of the fluorescence per $\mu m^2$ and coefficient of variation (CoV) for the QmNGMA actin marker in DMSO and LatA-treated larval fillets, measured through the middle of NMJs, where the actin core predominantly traverses. Error bars represent SEM. N in bar graphs – NMJs. ns – non-significant, * $p<0.05$, ** $p<0.01$, and *** $p<0.001$ upon unpaired, non-parametric Mann-Whitney test. Scale bars – 2 µm. **See Table 1 for detailed genotypes and N.**

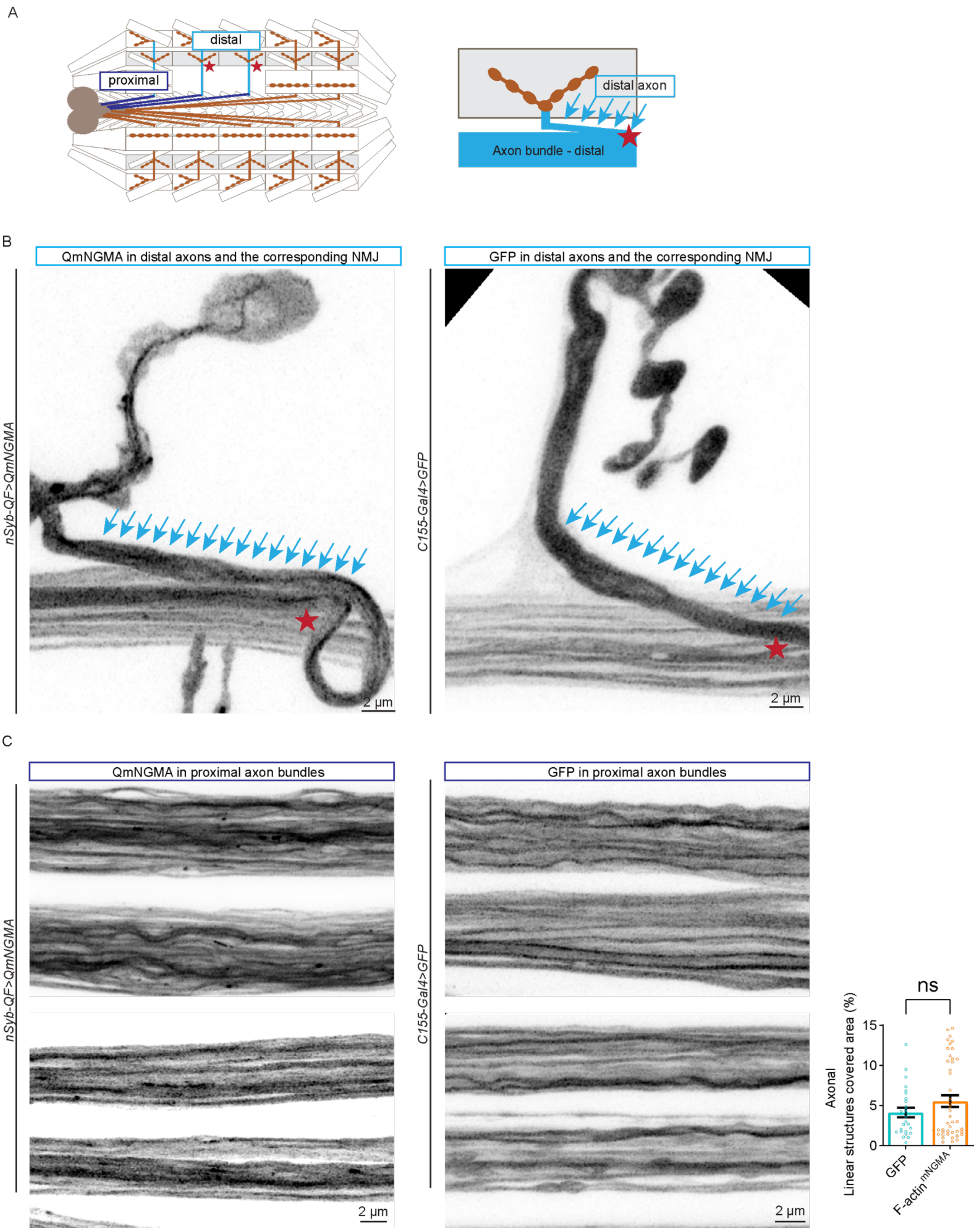
A
distal
proximal
distal axon
Axon bundle - distal
B
QmNGMA in distal axons and the corresponding NMJ
GFP in distal axons and the corresponding NMJ
nSyb-QF>QmNGMA
C155-Gal4>GFP
2 µm
2 µm
C
QmNGMA in proximal axon bundles
GFP in proximal axon bundles
nSyb-QF>QmNGMA
C155-Gal4>GFP
2 µm
2 µm
ns
Axonal
Linear structures covered area (%)
15
10
5
0
GFP
F-actin mNGMA

**Supplemental Figure 3. Assessment for linear F-actin in axons. A**) Diagrams of *Drosophila* fillet preparations denoting proximal and distal axons along the larva. For simplicity, in the diagram on the left, we denoted only three axonal bundles and a few muscles were left out. On right, distal axons branch out ("exit" marked with red star) from distal axonal bundles (light blue) to form NMJs. **B**) Distal axons (blue arrows) of larvae expressing neuronally (*nSybQF*) the F-actin marker QmNGMA feature possible linear F-actin structures along with visible linear F-actin at the NMJ. By contrast, larvae expressing neuronal (*C155-Gal4*) GFP exhibit diffuse GFP signal both in the distal axon and NMJ. **C**) In the narrow axonal shafts of proximal axonal bundles, putative linear F-actin QmNGMA structures are not distinguishable from the distribution of free GFP, either visually or quantified via WEKA segmentation. Error bars represent SEM. N – axons. ns – non-significant upon unpaired, non-parametric Mann-Whitney test.

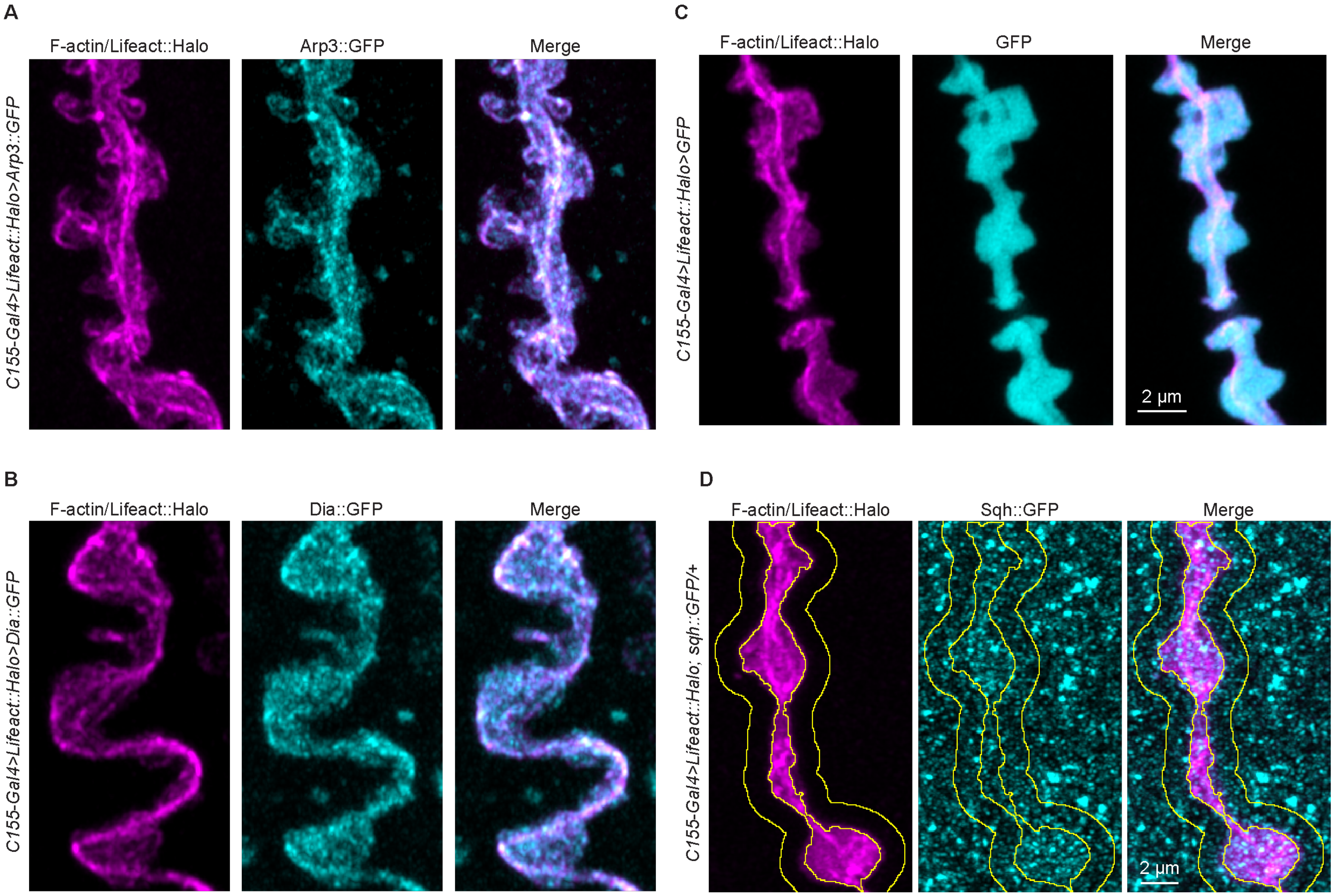

**Supplemental Figure 4. Actin-associated proteins at the presynaptic actin core. A**) Live imaging of the branched actin nucleator Arp2/3, visualized via Arp3::GFP (cyan), shows decoration of spot-like and linear F-actin assemblies labeled with Lifeact::Halo (magenta). See also **Supp. Movie S2**. **B**) The linear actin nucleator formin/Dia::GFP (cyan) decorates actin cables (magenta), in addition to its punctate distribution throughout the NMJ (see also **Supp. Movie S3**). **C**) Diffuse cytosolic GFP (cyan) shows no specific enrichment or co-localization with presynaptic actin assemblies (magenta); see also **Supp. Movie S4**. **D**) Distribution of the non-muscle myosin II light chain subunit sqh::GFP at the NMJ in live larvae expressing Lifeact::Halo in neurons. Inner yellow line denotes the Lifeact::Halo-positive neuronal region; outer yellow line denotes a 2 µm dilation defining the postsynaptic region. See also **Supp. Movie S5**. Scale bars – 2 µm. **See Table 1 for detailed genotypes.**

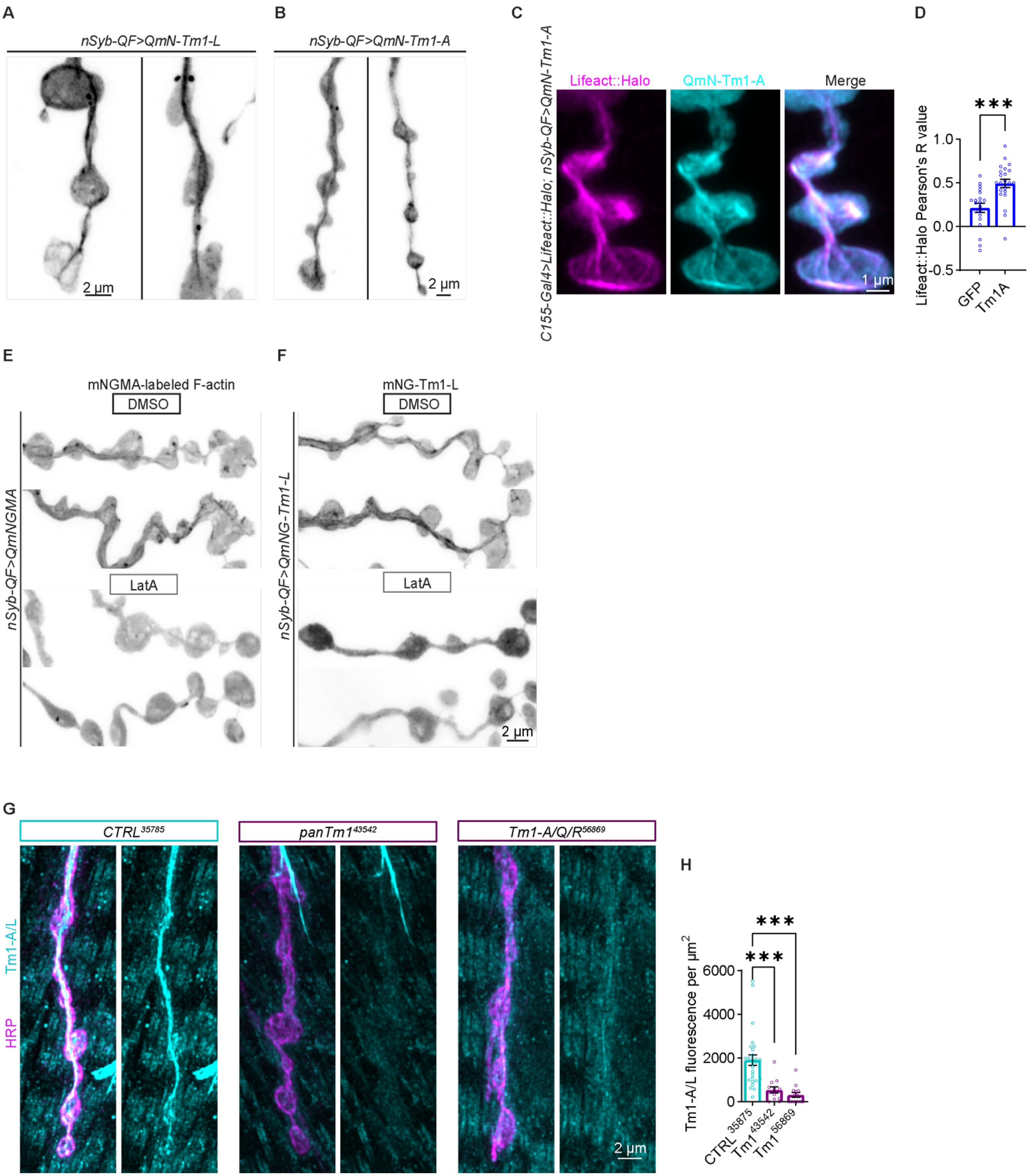
A
nSyb-QF>QmN-Tm1-L
2 µm
B
nSyb-QF>QmN-Tm1-A
2 µm
C
C155-Gal4>Lifeact::Halo; nSyb-QF>QmN-Tm1-A
Lifeact::Halo
QmN-Tm1-A
Merge
1 µm
D
Lifeact::Halo Pearson's R value
***
GFP
Tm1A
E
mNGMA-labeled F-actin
DMSO
LatA
nSyb-QF>QmNGMA
F
mNG-Tm1-L
DMSO
LatA
nSyb-QF>QmNG-Tm1-L
2 µm
G
CTRL35785
panTm143542
Tm1-A/Q/R56869
Tm1-A/L
HRP
2 µm
H
Tm1-A/L fluorescence per µm2
***
***
CTRL 35875
Tm1 43542
Tm1 56869

**Supplemental Figure 5. Association of the F-actin-stabilizing protein Tropomyosin with linear F-actin structures in the presynaptic core. A**) Inverted contrast single frame from live imaging depicting the localization pattern upon neuronal expression of mNeonGreen-tagged Tm1-L (mNG-Tm1-L or "long" Tm1) and Tm1-A (mN-Tm1-A or "short" Tm1) in **B**). **C**) The F-actin stabilizer Tm1-A (mNG-Tm1-A) is enriched in the linear Lifeact::Halo-marked actin assemblies along the presynaptic core. **D**) Pearson's R correlation between Lifeact::Halo and GFP or mNG-Tm1-A. **E**) The presynaptic localization of mNGMA-labeled actin assemblies and **F**) mNG-Tm1-L is sensitive to treatment with LatA, and not the vehicle DMSO, supporting F-actin-Tm1 association with the presynaptic actin core. **G**) Endogenous Tm1 (cyan), detected via immunofluorescence, traverses the NMJ (labeled with HRP – magenta) in larvae expressing control RNAi in all neurons. The Tm1 signal is significantly decreased at NMJs of animals neuronally expressing a Tm1 RNAi line targeting all known Tm1 isoforms ($panTm1^{43542}$), and the complementary line $Tm1^{56869}$, targeting the Tm1 isoforms A, Q and R. **H**) Quantification of the Tm1-A/L fluorescence per μm$^2$ at NMJs from larvae neuronally expressing *CTRL*, $Tm1^{43542}$ or $Tm1^{56869}$ RNAi lines. Ns – non-significant, *** p<0.001, and *** p<0.001 upon unpaired, non-parametric Mann-Whitney test, or after one-way ANOVA with Kruskal-Wallis multiple comparisons test. N in bar graphs – NMJs. Error bars are SEM. Scale bars – 2 μm. **See Table 1 for detailed genotypes and N.**

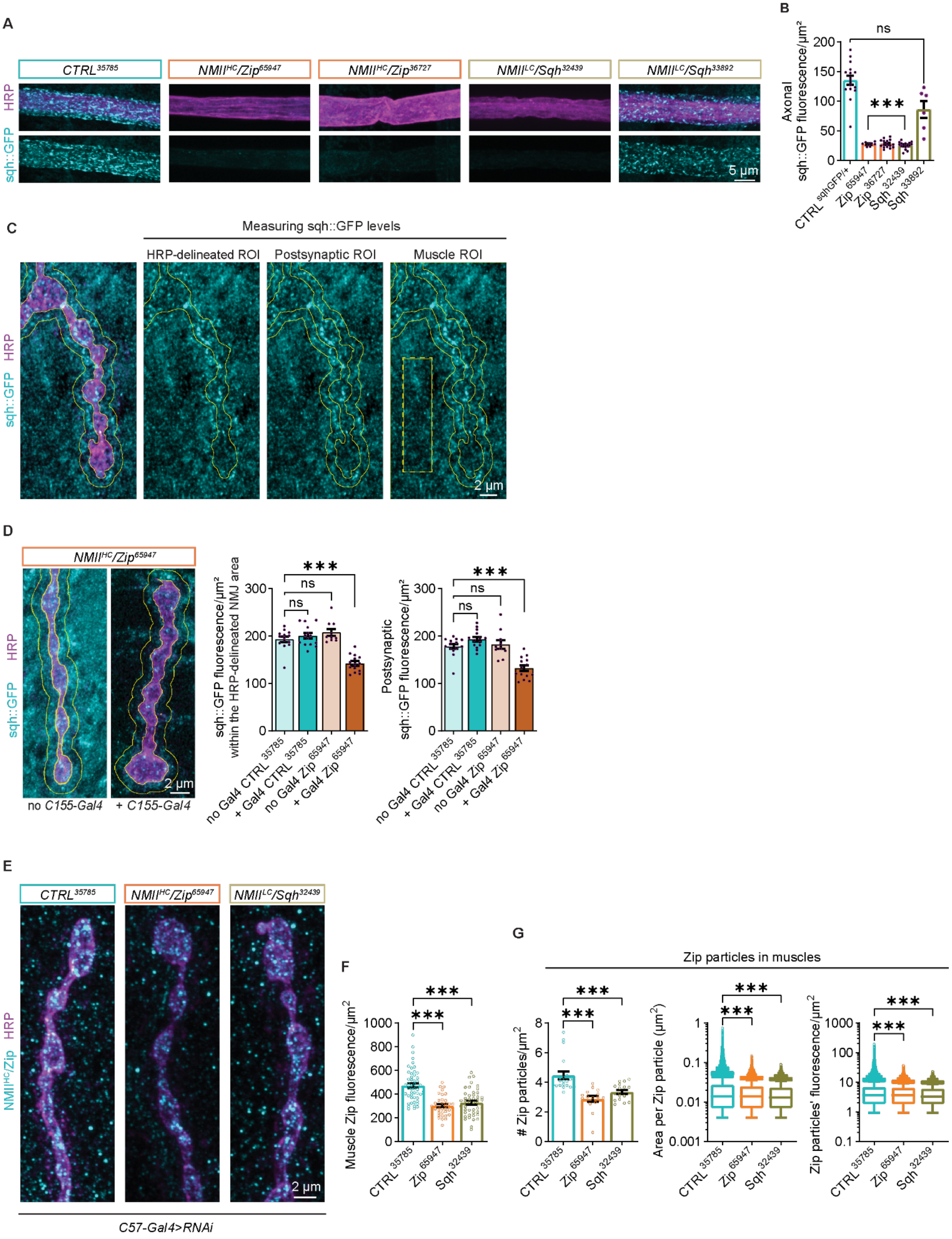
A
CTRL35785
NMIIHC/Zip65947
NMIIHC/Zip36727
NMIILC/Sqh32439
NMIILC/Sqh33892
HRP
sqh::GFP
5 µm
B
ns
***
Axonal sqh::GFP fluorescence/µm²
CTRL sqhGFP/+
Zip 65947
Zip 36727
Sqh 32439
Sqh 33892
C
Measuring sqh::GFP levels
HRP-delineated ROI
Postsynaptic ROI
Muscle ROI
2 µm
D
NMIIHC/Zip65947
no C155-Gal4
+ C155-Gal4
sqh::GFP fluorescence/µm² within the HRP-delineated NMJ area
Postsynaptic sqh::GFP fluorescence/µm²
no Gal4 CTRL 35785
+ Gal4 CTRL 35785
no Gal4 Zip 65947
+ Gal4 Zip 65947
E
CTRL35785
NMIIHC/Zip65947
NMIILC/Sqh32439
NMIIHC/Zip
C57-Gal4>RNAi
F
Muscle Zip fluorescence/µm²
G
Zip particles in muscles
### Zip particles/µm²
Area per Zip particle (µm²)
Zip particles' fluorescence/µm²
CTRL 35785
Zip 65947
Sqh 32439

**Supplemental figure 6. Validation of *UAS-RNAi* lines targeting NMII$^{HC}$/Zip and NMII$^{LC}$/Sqh.** **A**) Distribution of Sqh::GFP (cyan) in axons (labeled with HRP in magenta) proximal to muscle 4 in Sqh::GFP/+ heterozygous larvae, and larvae expressing in neurons two independent *UAS-RNAi* lines (*Zip*$^{65947}$ and *Zip*$^{36727}$) targeting NMII$^{HC}$/Zip, as well as the NMII$^{LC}$/Sqh targeting lines *Sqh*$^{32439}$ and *Sqh*$^{33892}$. **B**) Quantification of the Sqh::GFP fluorescence in axons of larvae with the different genotypes shown in **A**. **C**) Example of image analysis in Fiji, where the Sqh::GFP fluorescence was measured within the HRP delineated NMJ ROI, in a 1 µm rim surrounding the bouton (postsynaptic ROI), and in the muscle ROI (rectangle with yellow dashed lines) (excluding the HRP delineated area and the surrounding 1 µm rim). **D**) Quantifications of the Sqh::GFP fluorescence at NMJs from larvae carrying *Zip*$^{65947}$ in absence and presence of the neuronal *C155-Gal* driver, in an experiment validating the lack of nonspecific expression of *Zip*$^{65947}$ in neurons and muscles in absence of Gal4 driver. **E**) Immunostaining of endogenous Zip (cyan) at NMJs (HRP magenta) and muscles of larvae expressing *C57-Gal4*-driven control and RNAi lines targeting NMII$^{HC}$/Zip and NMII$^{LC}$/Sqh, respectively. **F**) Quantifications of muscle Zip fluorescence and, **G**) Zip particle abundance, size and brightness in larvae expressing *C57-Gal4*-driven *CTRL*$^{35785}$, *Zip*$^{65947}$ and *Sqh*$^{32439}$. Box and whiskers graphs were used to represent the results of the area and fluorescence intensity of the individual Zip particles. Whiskers represent 10$^{th}$ to 90$^{th}$ percentile, while the rest of the data points are shown as individual values. The Y-axis in these graphs represents log10, to capture the broad distribution of the individual values. In bar graphs with linear Y-axis, error bars represent SEM. N – axons in B, NMJs in D and G, muscle ROIs in F. N in box and whiskers – individual Zip particles. ** $p<0.01$, *** $p<0.001$ after one-way ANOVA with Kruskal-Wallis multiple comparisons test or upon unpaired, non-parametric Mann-Whitney test. Scale bar in A – 5 µm; all the rest – 2 µm. **See Table 1 for detailed genotypes and N.**

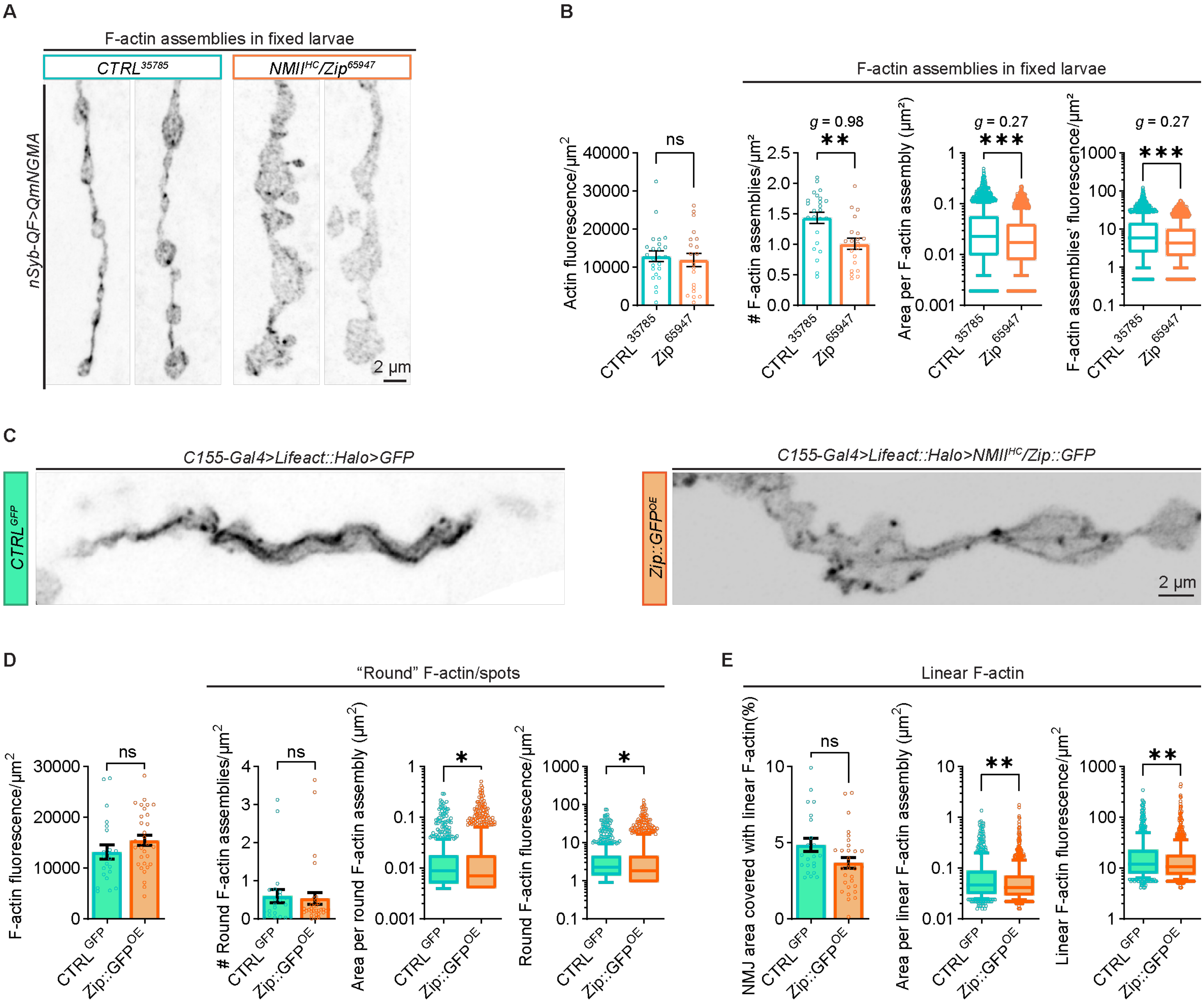

**Supplemental figure 7. Presynaptic actin assemblies upon down-regulation and overexpression of NMII$^{HC}$/Zip in neurons.** **A**) Inverted contrast mNGMA-labeled presynaptic actin in fixed larvae co-expressing *CTRL*$^{35785}$- or *NMII*$^{HC}$*/Zip*$^{65947}$ RNAi lines in neurons. **B**) Quantification of the fluorescence per µm$^2$ for the mNGMA actin marker in whole NMJs, along with the number of actin assemblies per µm$^2$, and the area and fluorescence intensity of the individual presynaptic actin structure. The actin particles were analyzed without specifying their morphological features as round or linear to account for fragmentation of the presynaptic cytoskeleton as a result of fixation[1]. **C**) Live-imaged inverted contrast Lifeact::Halo-labeled presynaptic actin in control larvae expressing GFP and larvae expressing NMII$^{HC}$/Zip::GFP. **D-E**) Analyses of the F-actin fluorescence, as well as individual WEKA-segmented round and linear F-actin entities in GFP- and Zip::GFP-expressing animals. Box and whiskers graphs were used to represent the results of the area and fluorescence intensity of the individual presynaptic actin structures. Whiskers represent 10$^{th}$ to 90$^{th}$ percentile, while the rest of the data points are shown as individual values. The Y-axis in these graphs represents log10, to capture the broad distribution of the individual values. In bar graphs with linear Y-axis, error bars represent SEM. N in bar graphs – NMJs; N in box and whiskers – individual actin assemblies. Ns – non-significant, ** p<0.01, *** p<0.001 upon unpaired, non-parametric Mann-Whitney test. Scale bars – 2 µm. **See Table 1 for detailed genotypes and N.**

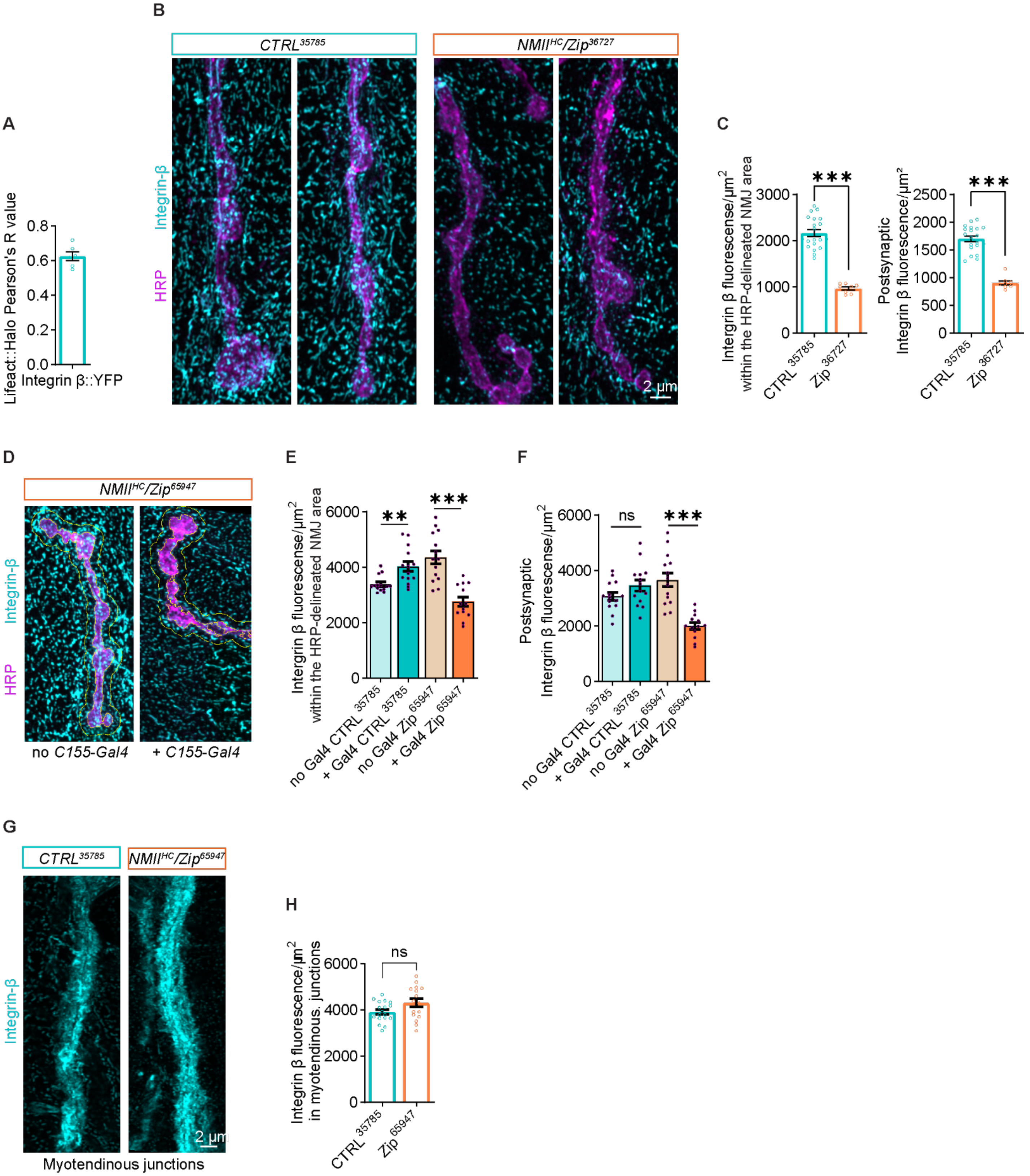
A
B
C
D
E
F
G
H
Lifeact::Halo Pearson's R value
Integrin β::YFP
CTRL35785
NMIIHC/Zip36727
Integrin-β
HRP
2 µm
Integrin β fluorescense/µm2 within the HRP-delineated NMJ area
Postsynaptic Integrin β fluorescence/µm2
***
NMIIHC/Zip65947
no C155-Gal4
+ C155-Gal4
**
ns
no Gal4 CTRL 35785
+ Gal4 CTRL 35785
no Gal4 Zip 65947
+ Gal4 Zip 65947
Myotendinous junctions
Integrin β fluorescence/µm2 in myotendinous. junctions
CTRL 35785
Zip 65947

**Supplemental Figure 8. Neuronal depletion of non-muscle Myosin II rearranges Integrin receptors at the NMJ. A)** Pearson's R correlation between Lifeact::Halo and Integrin-β::YFP at NMJs of live larvae. **B**) Reorganization of Integrin-β (cyan) at NMJs (HRP, magenta) of larvae expressing an independent RNAi line $NMII^{HC}/Zip^{36727}$ in neurons, compared to controls. **C**) Fluorescence intensity per µm$^2$ of Integrin-β within the HRP-delineated and postsynaptic areas in control and larvae expressing the independent $Zip^{36727}$. **D**) Integrin-β (cyan) immunofluorescence at NMJs (HRP, magenta) in larvae carrying $Zip^{65947}$ in absence and presence of *C155-Gal4* driver. **E-F**) Quantification of the fluorescence intensity per µm$^2$ of Integrin-β within the HRP-delineated and postsynaptic areas in larvae carrying the $CTRL^{35785}$- and $Zip^{65947}$ RNAi lines, in absence and presence of *C155-Gal4*. **G**) Integrin-β immunofluorescence (cyan) at myotendinous junctions of larvae expressing *CTRL*- and $Zip^{65947}$ in neurons. **H**) Quantification of the fluorescence per µm$^2$ of Integrin-β in myotendinous junctions in *CTRL*- and $Zip^{65947}$-expressing larvae (*C155-Gal4>RNAi*). Error bars represent SEM. N – NMJs or myotendinous junctions, respectively. Ns – non-significant, ** p<0.01, *** p<0.001 upon unpaired, non-parametric Mann-Whitney test. Scale bar – 2 µm. **See Table 1 for detailed genotypes and N.**

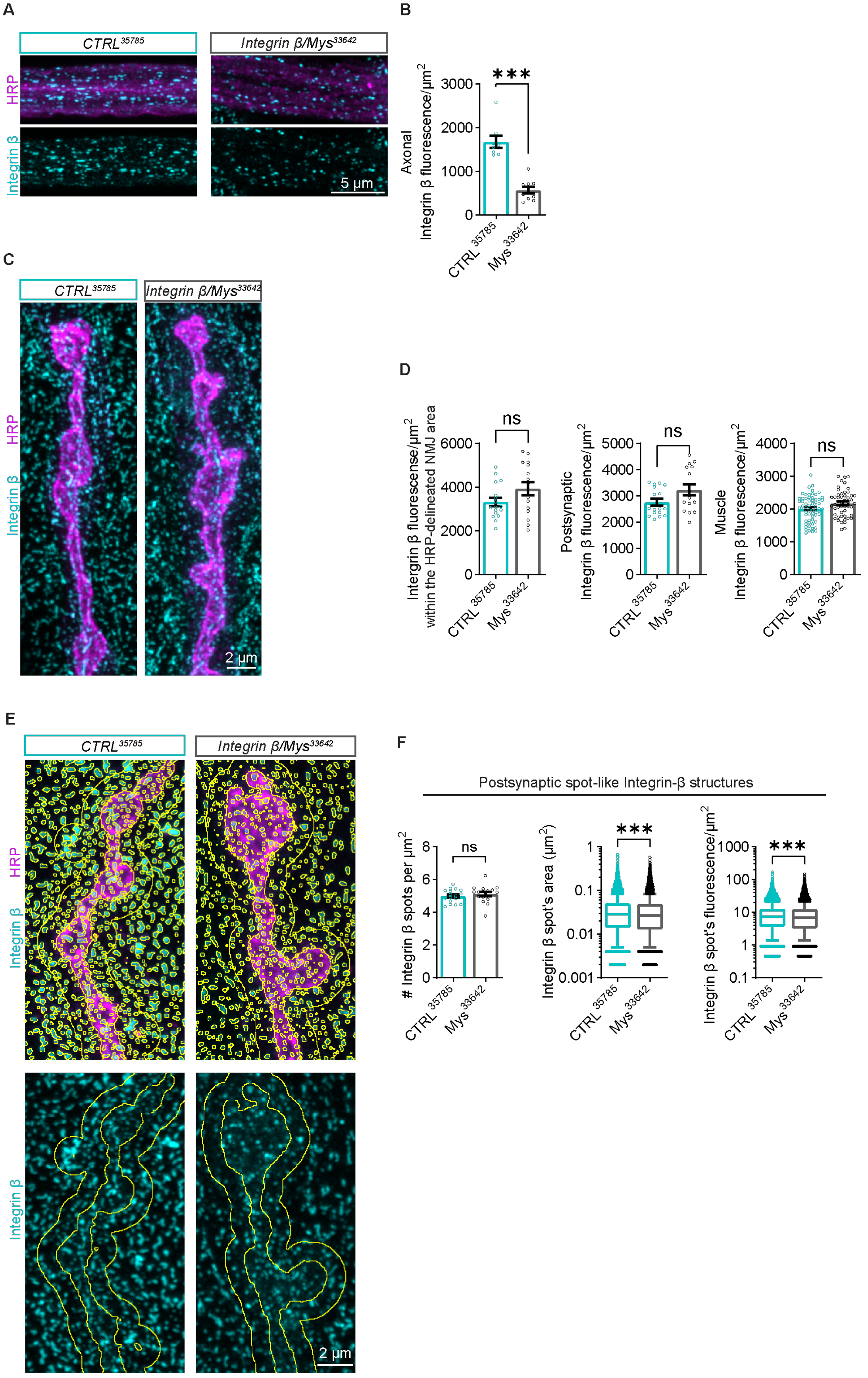
A
CTRL35785
Integrin β/Mys33642
HRP
Integrin β
5 µm
B
Axonal Integrin β fluorescence/µm2
3000
2000
1000
0
***
CTRL 35785
Mys 33642
C
CTRL35785
Integrin β/Mys33642
HRP
Integrin β
2 µm
D
Integrin β fluorescense/µm2 within the HRP-delineated NMJ area
6000
4000
2000
0
ns
Postsynaptic Integrin β fluorescence/µm2
5000
4000
3000
2000
1000
0
Muscle Integrin β fluorescence/µm2
E
CTRL35785
Integrin β/Mys33642
HRP
Integrin β
2 µm
F
Postsynaptic spot-like Integrin-β structures
### Integrin β spots per µm2
8
6
4
2
0
Integrin β spot's area (µm2)
1
0.1
0.01
0.001
Integrin β spot's fluorescence/µm2
1000
100
10
1
0.1

**Supplemental Figure 9. Organization of integrin receptors at the NMJ upon Integrin-β neuronal down-regulation. A)** Integrin-β (cyan) in axonal bundles (HRP in magenta) in control and larvae expressing the *Integrin-β/Mys*$^{33642}$ RNAi in neurons (*C155-Gal4*). **B**) Quantification of axonal Integrin-β fluorescence in *CTRL*$^{35785}$ vs *Mys*$^{33642}$ neuronal RNAi. **C**) Integrin-β (cyan) at NMJs (HRP in magenta) in control and *Integrin-β/Mys*$^{33642}$ animals. **D**) Analyses of Integrin-β fluorescence in HRP-delineated and postsynaptic areas, as well as in muscles of control and *Integrin-β/Mys*$^{33642}$ larvae. **E**) Integrin-β WEKA-segmented particles in controls and *Integrin-β/Mys*$^{33642}$ larvae. **F**) Analyses of spot-like Integrin-β structures within the postsynaptic ROI. Control datasets are reused from Fig. 6. Box and whiskers graphs were used to represent the results of the area and fluorescence intensity of the individual Integrin-β particles. Whiskers represent 10$^{th}$ to 90$^{th}$ percentile, while the rest of the data point are shown as individual values. The Y-axis in these graphs represents log10, to capture the broad distribution of the individual values. In bar graphs with linear Y-axis, error bars represent SEM. N in bar graphs – NMJs; N in box and whiskers – individual Integrin-β assemblies. Ns – non-significant, and *** $p<0.001$ upon unpaired, non-parametric Mann-Whitney test. Error bars are SEMs. Scale bar – 2 µm. **See Table 1 for detailed genotypes and N.**

**Legends for Supplementary Movies 1-11:**

**Supplementary Movie S1.** Time-lapse movie showing QmNGMA-labeled presynaptic F-actin structures at the NMJ. Each frame is a maximum intensity projection of Z-stacks of presynaptic terminals imaged at a sampling frequency of 1 frame/5 sec. The movie is played at a speed of 2 frames per second. Scale bar – 2 µm.

**Supplementary Movie S2.** Time-lapse movie of Arp3::GFP (cyan) and the F-actin marker Lifeact::Halo (magenta) at the NMJ. Frames represent maximum intensity projections of Z-stacks of presynaptic terminals imaged at a sampling frequency of 1 frame/2.16 s. The movie is played at a speed of 5 frames per second. Scale bar – 2 µm.

**Supplementary Movie S3.** Time-lapse movie of Dia::GFP (cyan) and the F-actin marker Lifeact::Halo (magenta) at the NMJ. Frames represent maximum intensity projections of Z-stacks of presynaptic terminals imaged at a sampling frequency of 1 frame/1.3 s. The movie is played at a speed of 5 frames per second. Scale bar – 2 µm.

**Supplementary Movie S4.** Time-lapse movie of GFP (cyan) and the F-actin marker Lifeact::Halo (magenta) at the NMJ. Frames represent maximum intensity projections of Z-stacks of presynaptic terminals imaged with 1/13.32 s sampling frequency. The movie is played at a speed of 1 frame per second. Scale bar – 2 µm.

**Supplementary Movie S5.** Time-lapse movie of sqh::GFP (cyan) and the F-actin marker Lifeact::Halo (magenta) at the NMJ. Frames represent maximum intensity projections of Z-stacks of presynaptic terminals imaged at a sampling frequency of 1 frame/13.38 s. The movie is played at a speed of 1 frame per second. Scale bar – 2 µm.

**Supplementary Movie S6.** Time-lapse movie of Zip::GFP (cyan) and the F-actin marker Lifeact::Halo (magenta) at the NMJ. Frames represent maximum intensity projections of Z-stacks of presynaptic terminals imaged at a sampling frequency of 1 frame/4 s. The movie is played at a speed of 1 frame per second. Scale bar – 2 µm.

**Supplementary Movie S7.** Time-lapse movie showing QmNGMA-labeled presynaptic F-actin structures at the NMJ in larvae expressing *CTRL* RNAi in neurons. Frames represent maximum intensity projections of Z-stacks of presynaptic terminals imaged at a sampling frequency of with 1 frame/5 s. The movie is played at a speed of 1 frame per second. Scale bar – 2 µm.

**Supplementary Movie S8.** Time-lapse movie showing QmNGMA-labeled presynaptic F-actin structures at the NMJ in larvae expressing *Zip*$^{65947}$ RNAi in neurons. Frames represent maximum intensity projections of Z-stacks of presynaptic terminals imaged at a sampling frequency of 1 frame/5 s. The movie is played at a speed of 1 frame per second. Scale bar – 2 µm.

**Supplementary Movie S9.** Ventral (muscle proximal) to dorsal (muscle distal) Z-stack played as a movie showing ubiquitously expressed Integrin-β (cyan) and the F-actin marker Lifeact::Halo (magenta) through an NMJ. Scale bar – 2 µm.

**Supplementary Movie S10.** Time-lapse movie showing QmNGMA-labeled presynaptic F-actin structures at the NMJ in larvae with intact brain and axons. Frames represent maximum intensity projections of Z-stacks of presynaptic terminals imaged at a sampling frequency of 1 frame/1.93 s. The movie is played at a speed of 5 frames per second speed. Scale bar – 2 μm.

**Supplementary Movie S11.** Time-lapse movie showing QmNGMA-labeled presynaptic F-actin structures at the NMJ in larvae with severed brain and axons. Frames represent maximum intensity projections of Z-stacks of presynaptic terminals imaged at a sampling frequency of 1 frame/1.93 s. The movie is played at a speed of 5 frames per second. Scale bar – 2 μm.

**Supplementary reference:**

1. Pereira, P.M. *et al.* Fix Your Membrane Receptor Imaging: Actin Cytoskeleton and CD4 Membrane Organization Disruption by Chemical Fixation. *Front Immunol* **10**, 675 (2019).

**Resource Data Table**

| REAGENT or RESOURCE | SOURCE | IDENTIFIER |
|---|---|---|
| **Antibodies** | | |
| FluoTag®-X4 anti-GFP (1:250) | NanoTag Biotechnologies | N0304 |
| Rabbit anti-Tm1A/L (1:500) | Denise Montel, University of California, Santa Barbara | NA |
| Mouse anti-Integrin betaPS (1:50) | Developmental studies hybridoma bank | Cat# cf.6g11 |
| Rabit anti-Zip (1:1000) | Anna Marie Sokac (UIIC) | NA |
| Rhodamine Red™-X (RRX) AffiniPure Goat Anti-Horseradish Peroxidase (1:500) | Jackson ImmunoResearch | 123-295-021 |
| CF®488A Donkey Anti-Rabbit IgG (H+L), Highly Cross-Adsorbed Antibody | Biotium, Inc. | 20015 |
| CF®488A Donkey Anti-Mouse IgG (H+L), Highly Cross-Adsorbed Antibody | Biotium, Inc. | 20014 |
| Alexa Fluor® 488 AffiniPure Goat Anti-Mouse IgG (H+L) | Jackson ImmunoResearch | 111-545-003 |
| **Chemicals, peptides and recombinant proteins** | | |
| DMSO | | |
| InSolution Latrunculin A | Sigma | 428026 |
| Janelia Fluor® HaloTag® Ligand-549 | Promega | GA111A |
| **Experimental models: Organisms/strains** | | |
| *D. melanogaster* QUAS-mNeonGreen::GMA | This paper | NA |
| *D. melanogaster QUAS-mScarlet-I::Act5C* | This paper | NA |
| *D. melanogaster QUAS-Tm1-A* | This paper | NA |
| *D. melanogaster QUAS-Tm1-L* | This paper | NA |
| *D. melanogaster UAS-Lifeact::Halo2* | BDSC | Stock#: *67625* |
| *D. melanogasterUAS-Arp3::GFP* | BDSC | Stock#: *39723* |
| *D. melanogaster UAS-Dia::GFP* | BDSC | Stock#: *56751* |
| *D. melanogaster UAS-GFP* | Leslie Griffith, Brandeis University | |
| *D. melanogaster* UASp-sqh.A20A21 | BDSC | Stock#: *64114* |
| *D. melanogaster UASp-sqh.E20E21* | BDSC | Stock#: *64411* |
| *D. melanogaster UAS-CTRL-RNAi35785* | BDSC | Stock#: *35785* |
| *D. melanogaster UAS-Zip-RNAi65947* | BDSC | Stock#: *65947* |
| *D. melanogaster UAS-Zip-RNAi32727* | BDSC | Stock#: *32727* |
| *D. melanogaster UAS-Sqh-RNAi32439* | BDSC | Stock#: *32439* |
| *D. melanogaster UAS-Sqh-RNAi33892* | BDSC | Stock#: *33892* |
| *D. melanogaster UAS-Tm1-RNAi43542* | BDSC | Stock#: *43542* |
| *D. melanogaster UAS-Tm1-RNAi56869* | BDSC | Stock#: *56869* |
| *D. melanogaster UAS-Mys-RNAi33642* | BDSC | Stock#: *33642* |
| *D. melanogaster UAS-Zip::GFP* | Adam Martin, MIT | NA |
| *D. melanogaster sqh::GFP* | BDSC | Stock#: 57145 |
| *D. melanogaster P{Ubi-mys.YFP}* | Guy Tanentzapf, University of British Columbia | Yuan et al., 2010 |
| *D. melanogaster C155-Gal4* | BDSC | Stock#: 458 |
| *D. melanogaster nSyb-QF2* | BDSC | derived from Stock#: 51954 |

| | | |
|---|---|---|
| *D. melanogaster C57-Gal4* | Budnik et al, 1996 | https://doi.org/10.1016/S0896-6273(00)80196-8 |
| **Software and algorithms** | | |
| Fiji | | |
| GraphPad Prism | | |

**Table 1.** Experimental genotypes and statistical reporting.

| Figure | Genotypes/Conditions | N | Measurement | Statistical Test(s) |
|---|---|---|---|---|
| **Figure 1** | **Fig. 1. Presynaptic actin cytoskeleton at the *Drosophila* larval NMJ.** | | | |
| **Fig. 1A:** Diagram of the *Drosophila* larval neuromuscular system as a tissue mechanics model. | | | | |
| **Fig. 1B:** A single frame from live imaging of presynaptic actin assemblies visualized via neuronal expression of the genetically encoded actin marker mNeonGreen-GMA (mNGMA) in larval preparations with intact brains<br><br>Repeated in multiple experiments. | *C155-Gal4/Y; QUAS-mNeonGreenMA(GMA)/+; nSyb-QF2/+* | | | |
| **Fig. 1C:** A single frame from live imaging of NMJs expressing an independent genetically encoded actin marker Lifeact::Halo delineates similar spot-like and cable-like (core) actin assemblies.<br>Repeated in multiple experiments. | *C155-Gal4/Y;UAS-Lifeact//Halo:+* | | | |
| **Fig. 1D:** A single frame from live imaging of the *Drosophila* actin isoform 5C, tagged with the red fluorescent protein mScarlet-I (QAct5C::mScarI), similarly delineates the actin core in addition to round F-actin assemblies.<br>Repeated in single experiment, multiple crosses. | *C155-Gal4/Y; QUAS-Act5C::mScarI/+; nSyb-QF2/+* | 9 larva/ 16 NMJs | | |
| **Fig. 1E:** Image sequences of FRAP of QAct5C::mScarI-labeled actin core. | *C155-Gal4/Y; QUAS-Act5C::mScarI/+; nSyb-QF2/+* | 9 larvae/ 16 NMJs | | |
| **Fig. 1F:** FRAP curve of QAct5C::mScarI | *C155-Gal4/Y; QUAS-Act5C::mScarI/+; nSyb-QF2/+* | 9 larvae/ 16 NMJs | Actin fluorescence per μm$^2$ | None. |

| Figure 2. | **Fig. 2. Distribution of endogenous and neuronally expressed NMII$^{HC}$/Zip at *Drosophila* larval NMJs.** | | | |
|---|---|---|---|---|
| **Fig.2A:** Immunostaining of endogenous NMII$^{HC}$/Zip (cyan) at NMJs (HRP, magenta) of wild-type larvae labels large and small puncta at the NMJ and in muscles (yellow arrows – large Zip, white arrows – small Zip structures). Smooth, aggregate-like Zip puncta are visible predominantly in muscles (gray arrows).<br>Repeated in two independent experiments, multiple crosses | *C155-Gal4/Y;; CTRL$^{35785}$/ +* | | | None |
| **Fig. 2B:** Zip puncta (cyan) in distal axonal bundles (HRP in magenta) in animals neuronally (*C155-Gal4*) expressing *CTRL$^{35785}$*-, *NMII$^{HC}$/Zip$^{65947}$* and *NMII$^{LC}$/Sqh$^{32439}$* RNAi lines. | *C155-Gal4/Y;;CTRL$^{35785}$/+*<br><br>*C155-Gal4/Y;; NMII$^{HC}$/Zip$^{65947}$ /+*<br><br>*C155-Gal4/Y ;;NMII$^{LC}$/Sqh$^{32439}$/+* | | | None |
| **Fig. 2C:** Quantification of Zip fluorescence in axonal bundles.<br>Repeated in two independent experiments, multiple crosses. | *C155-Gal4/Y;;CTRL$^{35785}$/+*<br><br>*C155-Gal4/Y;; NMII$^{HC}$/Zip$^{65947}$ /+*<br><br>*C155-Gal4/Y ;;NMII$^{LC}$/Sqh$^{32439}$/+* | 20 axons (10 larvae) | Zip fluorescence per µm$^2$ | ns – not significant, *** p<0.001 after One-way ANOVA with Kruskal-Wallis multiple comparison test.<br>N – axons |
| **Fig. 2D:** Live imaging of neuronally co-expressed NMII$^{HC}$/Zip::GFP (cyan) and Lifeact::Halo (magenta) similarly reveals Zip puncta with different size at the NMJ, some of which are enriched at spot-like and linear F-actin assemblies (see also **Supp. Movie S6**).<br>Repeated in two independent experiments, multiple crosses | *C155-Gal4/Y; UAS-Lifeact::Halo/+; UAS-Zip::GFP/+* | | | |
| **Fig. 2E:** Independent examples of live-imaged NMJs of larvae expressing Zip::GFP and Lifeact::Halo. White arrows point to putative bipolar Zip filaments. | *C155-Gal4/Y; UAS-Lifeact::Halo/+; UAS-Zip::GFP/+* | | | |

| | | | | |
|---|---|---|---|---|
| **Fig. 2F:** Pearson's R correlation between Lifeact::Halo and Zip::GFP. | *C155-Gal4/Y; UAS-Lifeact::Halo/+: UAS-GFP/+*<br><br>*C155-Gal4/Y; UAS-Lifeact::Halo/+: UAS-Zip::GFP/+* | 23 NMJs (8 larvae)<br><br>38 NMJs (9 larvae) | | ns – not significant, *** p<0.0001, upon unpaired, non-parametric Mann-Whitney test.<br>N – number of NMJs |
| **Fig. 2G:** Quantification of the abundance and size of smaller and larger Zip puncta at the NMJ, as well as the average distance between adjacent Zip puncta in putative bipolar Zip filaments. | *C155-Gal4/Y; UAS-Lifeact::Halo/+: UAS-Zip::GFP/+* | 31 NMJs (9 larvae) | Tiny Zip spots per $\mu m^2$<br><br>Larger Zip spots per $\mu m^2$ | N – number of NMJs |
| | | Small Zip spots = 27855 (9 larvae/31 NMJs)<br><br>Larger Zip spots = 1047 (9 larvae/31 NMJs) | Area per Zip spots ($\mu m^2$) | *** p<0.0001, upon unpaired, non-parametric Mann-Whitney test.<br><br>N in box and whiskers – individual Zip assemblies |
| | | 41 Zip doublets (8 larvae) | Distance between Zip doublet's maxima | N – number of Zip doublets |
| **Figure 3.** | **Fig. 3. Non-muscle Myosin II regulates the integrity of presynaptic actin assemblies.** | | | |
| **Fig. 3A:** mNGMA-labeled presynaptic actin in larvae co-expressing *CTRL RNAi* line. See also **Supp. Movie S6**.<br>Repeated in two independent experiments, multiple crosses. | *C155-Gal4/Y; QUAS-mNeonGreenMA(GMA)/UAS-CTRL$^{35785}$; nSyb-QF2/+* | 8 larvae/ 23 NMJs | | |

| | | | | |
|---|---|---|---|---|
| **Fig. 3B:** mNGMA-labeled presynaptic actin in larvae co-expressing *NMII/Zip$^{65947}$* RNAi lines. See also **Supp. Movie S7**. | *C155-Gal4/Y; QUAS-mNeonGreenMA(GMA)/UAS-Zip$^{65947}$; nSyb-QF2/+* | 9 larvae/ 26 NMJs | | |
| **Fig. 3C:** Quantification of fluorescence per µm$^2$ for the QmNGMA actin marker in whole NMJs. | *C155-Gal4/Y;QUAS-mNeonGreenMA(GMA)/UAS-CTRL$^{35785}$; nSyb-QF2/+*<br><br>*C155-Gal4/Y;QUAS-mNeonGreenMA(GMA)/UAS-Zip$^{65947}$; nSyb-QF2/+* | 8 larvae/ 23 NMJs<br><br>9 larvae/ 26 NMJs | Actin fluorescence per µm$^2$ | ns – not significant, upon unpaired, non-parametric Mann-Whitney test.<br>N – NMJs |
| **Fig. 3D:** Quantifications of the number of round F-actin assemblies per µm$^2$, as well as the area and fluorescence of the individual round presynaptic actin structures. | *C155-Gal4/Y; QUAS-mNeonGreenMA(GMA)/UAS-CTRL$^{35785}$; nSyb-QF2/+*<br><br>*C155-Gal4/Y; QUAS-mNeonGreenMA(GMA)/UAS-Zip$^{65947}$; nSyb-QF2/+* | 8 larvae/ 23 NMJs<br><br>9 larvae/ 26 NMJs | Number of round F-actin assemblies per µm$^2$ | ns – not significant, * $p<0.05$,<br>upon unpaired, non-parametric Mann-Whitney test.<br>N – NMJs |
| | | 8 larvae/ 23 NMJs/976 actin spots (N)<br><br>9 larvae/ 26 NMJs/1010 actin spots (N) | Area per round F-actin assembly (µm$^2$) | ns – not significant, upon unpaired, non-parametric Mann-Whitney test.<br>N – number of individual round F-actin assemblies |
| | | 8 larvae/ 23 NMJs/976 actin spots (N)<br><br>9 larvae/ 26 NMJs/1010 actin spots (N) | Round F-actin fluorescence per µm$^2$ | |
| **Fig. 3E:** Graphs representing the percentage of NMJ area covered with linear F-actin, as well as the area and fluorescence of the individual assemblies. | *C155-Gal4/Y; QUAS-mNeonGreenMA(GMA)/UAS-CTRL$^{35785}$; nSyb-QF2/+*<br><br>*C155-Gal4/Y; QUAS-mNeonGreenMA(GMA)/UAS-Zip$^{65947}$; nSyb-QF2/+* | 8 larvae/ 23 NMJs<br><br>9 larvae/ 26 NMJs | NMJ area covered with linear F-actin assemblies (%) | ns – not significant, ** $p<0.01$, *** $p<0.001$, upon unpaired, non-parametric Mann-Whitney test.<br>N – NMJs |
| | | 8 larvae/ 23 NMJs/519 structures<br><br>9 larvae/ 26 NMJs/821 structures | Area per linear F-actin assembly (µm$^2$) | ns – not significant, ** $p<0.01$,<br>upon unpaired, non-parametric Mann-Whitney test. |

| | | | | |
|---|---|---|---|---|
| | | 8 larvae/ 23 NMJs/519 linear assemblies (N)<br><br>9 larvae/ 26 NMJs/821 linear assemblies (N) | Linear F-actin fluorescence per $\mu m^2$ | N – number of linear F-actin assemblies |
| **Fig. 3F-H:** Lifeact::Halo-labeled presynaptic F-actin in *GFP*-expressing control, or larvae expressing unphosphorylatable $Sqh^{DN}$ and constitutively phosphorylated $Sqh^{CA}$, respectively<br>Repeated in two independent experiments, multiple crosses. | *C155-Gal4/Y; UAS-Lifeact::Halo/+: UAS-GFP/+*<br><br>*C155-Gal4/Y; UAS-Lifeact::Halo/+: UAS-$Sqh^{DN}$/+*<br><br>*C155-Gal4/Y; UAS-Lifeact::Halo/+: UAS-$Sqh^{CA}$/+* | | | |
| **Fig. 3I:** Quantification of fluorescence per $\mu m^2$ for the Lifeact::Halo F-actin marker in whole NMJs. | *C155-Gal4/Y; UAS-Lifeact::Halo/+: UAS-GFP/+*<br><br>*C155-Gal4/Y; UAS-Lifeact::Halo/+: UAS-$Sqh^{DN}$/+*<br><br>*C155-Gal4/Y; UAS-Lifeact::Halo/+: UAS-$Sqh^{CA}$/+* | 31 NMJs (12 larvae)<br><br>29 NMJs (10 larvae)<br><br>29 NMJs (9 larvae) | Actin fluorescence per $\mu m^2$ | ns – not significant, after One-way ANOVA with Kruskal-Wallis multiple comparison test.<br><br>N – NMJs |
| **Fig. 3J:** Quantifications of the number of round F-actin assemblies per $\mu m^2$, as well as their area and fluorescence. | *C155-Gal4/Y; UAS-Lifeact::Halo/+: UAS-GFP/+*<br><br>*C155-Gal4/Y; UAS-Lifeact::Halo/+: UAS-$Sqh^{DN}$/+*<br><br>*C155-Gal4/Y; UAS-Lifeact::Halo/+: UAS-$Sqh^{CA}$/+* | 33 NMJs (12 larvae)<br><br>29 NMJs (10 larvae)<br><br>28 NMJs (9 larvae) | Number of round F-actin assemblies per $\mu m^2$ | ns – not significant, after One-way ANOVA with Kruskal-Wallis multiple comparison test.<br><br>N – NMJs |
| | | 1239 spots 33 NMJs (12 larvae)<br><br>1186 spots 29 NMJs (10 larvae)<br><br>1253 spots | Area per round F-actin assembly ($\mu m^2$) | ns – not significant, and *** $p<0.001$ after One-way ANOVA with Kruskal-Wallis multiple comparison test. |

| | | | | |
|---|---|---|---|---|
| | | (28 NMJs/9 larvae) | | N – round F-actin assemblies |
| | | 1239 spots<br>33 NMJs<br>(12 larvae)<br><br>1186 spots<br>29 NMJs (10 larvae)<br><br>1253 spots<br>(28 NMJs/9 larvae) | Round F-actin fluorescence per $\mu m^2$ | |
| **Fig. 3K:** Graphs representing the percentage of NMJ area covered with linear F-actin, as well as the area and fluorescence of the individual assemblies.<br>Repeated in two independent experiments, multiple crosses. | *C155-Gal4/Y; UAS-Lifeact::Halo/+: UAS-GFP/+*<br><br>*C155-Gal4/Y; UAS-Lifeact::Halo/+: UAS-Sqh*$^{DN}$*/+*<br><br>*C155-Gal4/Y; UAS-Lifeact::Halo/+: UAS-Sqh*$^{CA}$*/+* | 33 NMJs<br>(12 larvae)<br><br>29 NMJs (10 larvae)<br><br>28 NMJs (9 larvae) | NMJ area covered with linear F-actin assemblies (%) | ** p=0.005 and * p=0.04 after One-way ANOVA with Kruskal-Wallis multiple comparison test.<br><br>N – NMJs |
| | | 792 structures<br>(33 NMJs/12 larvae)<br><br>652 structures<br>(29 NMJs/10 larvae)<br><br>518 structures<br>(28 NMJs/9 larvae) | Area per linear F-actin assembly ($\mu m^2$) | *** p<0.001 and ns after One-way ANOVA with Kruskal-Wallis multiple comparison test.<br><br>N – linear F-actin assemblies |
| | | 792 structures<br>(33 NMJs/12 larvae)<br><br>652 structures<br>(29 NMJs/10 larvae)<br><br>518 structures<br>(28 NMJs/9 larvae) | Linear F-actin fluorescence per $\mu m^2$ | |
| **Figure 4.** | **Fig. 4 Depletion of neuronal NMII$^{HC}$ reduces the levels of the NMII$^{LC}$ subunit in the muscle.** | | | |

| | | | | |
|---|---|---|---|---|
| **Fig. 4A:** Sqh::GFP-labeled NMII at NMJs in fixed larvae expressing *CTRL*$^{35785}$, *Zip*$^{65947}$, and *Zip*$^{36727}$ RNAi in neurons (*C155-Gal4*). Repeated in three independent experiments, multiple crosses | *C155-Gal4/Y;; sqh::GFP/UAS-CTRL*$^{35785}$<br><br>*C155-Gal4/Y;; sqh::GFP/UAS-Zip*$^{65947}$<br><br>*C155-Gal4/Y;; sqh::GFP/UAS-Zip*$^{36727}$ | | | |
| **Fig. 4B:** Analysis of the Sqh::GFP fluorescence per µm$^2$ within the HRP delineated NMJ area, postsynaptic Sqh::GFP fluorescence per µm$^2$ and muscle Sqh::GFP fluorescence per µm$^2$ | *C155-Gal4/Y;; sqh::GFP/UAS-CTRL*$^{35785}$<br><br>*C155-Gal4/Y;; sqh::GFP/UAS-Zip*$^{65947}$<br><br>*C155-Gal4/Y;; sqh::GFP/UAS-Zip*$^{36727}$ | 10 larvae/ 20 NMJs<br><br>7 larvae/ 15 NMJs<br><br>7 larvae/ 14 NMJs | sqh::GFP fluorescence per µm$^2$ in HRP-delineated NMJ area<br>Postsynaptic sqh::GFP fluorescence per µm$^2$<br><br>Muscle sqh::GFP fluorescence per µm$^2$ | ** $p<0.01$, *** $p<0.001$ after one-way ANOVA with Šídák's multiple comparisons test.<br>N – NMJs and muscle ROIs |
| **Fig. 4C:** Example of WEKA-segmented individual sqh::GFP particles in the HRP-delineated- and postsynaptic compartments. | | | | |
| **Fig. 4D:** Analysis of the number, area and fluorescence WEKA-segmented sqh::GFP particles within the HRP-delineated NMJ area. | *C155-Gal4/Y;; sqh::GFP/UAS-CTRL*$^{35785}$<br><br>*C155-Gal4/Y;; sqh::GFP/UAS-Zip*$^{65947}$<br><br>*C155-Gal4/Y;; sqh::GFP/UAS-Zip*$^{36727}$ | 10 larvae/ 20 NMJs<br><br>7 larvae/ 15 NMJs<br><br>7 larvae/ 14 NMJs | Number of sqh::GFP particles per µm$^2$ HRP-delineated area | N – NMJs |
| | | 10 larvae/ 20 NMJs/ 11360 particles (N)<br><br>7 larvae/ 15 NMJs/5544 (N)<br><br>7 larvae/ 14 NMJs/5007 (N) | sqh::GFP particles' area (µm$^2$) | N – sqh::GFP particles |
| | | 10 larvae/ 20 NMJs/ 11360 particles (N) | sqh::GFP particles' fluorescence per µm$^2$ | |

| | | | | |
|---|---|---|---|---|
| | | 7 larvae/ 15 NMJs/5544 (N)<br><br>7 larvae/ 14 NMJs/5007 (N) | | |
| **Fig. 4E:** Analysis of the postsynaptic Sqh::GFP fluorescence per µm$^2$, as well as the number, area and fluorescence of WEKA-segmented individual sqh::GFP particles. | *C155-Gal4/Y;; sqh::GFP/UAS-CTRL$^{35785}$*<br><br>*C155-Gal4/Y;; sqh::GFP/UAS-Zip$^{65947}$*<br><br>*C155-Gal4/Y;; sqh::GFP/UAS-Zip$^{36727}$* | 10 larvae/ 20 NMJs<br><br>7 larvae/ 15 NMJs<br><br>7 larvae/ 14 NMJs | Postsynaptic sqh::GFP fluorescence per µm$^2$ | ** p<0.01, *** p<0.001 after one-way ANOVA with Šídák's multiple comparisons test.<br>N – NMJs |
| | | 10 larvae/ 20 NMJs<br><br>7 larvae/ 15 NMJs<br><br>7 larvae/ 14 NMJs | Number of sqh::GFP particles per µm$^2$ | |
| | | 10 larvae/ 20 NMJs/20096 particles<br><br>7 larvae/ 15 NMJs/9044 particles<br><br>7 larvae/ 14 NMJs/ 8916 particles | sqh::GFP particles' area (µm$^2$) | N – sqh::GFP particles |
| | | 10 larvae/ 20 NMJs/20096 particles<br><br>7 larvae/ 15 NMJs/9044 particles<br><br>7 larvae/ 14 NMJs/ 8916 particles | sqh::GFP particles' fluorescence per µm$^2$ | |
| **Figure 5.** | **Fig. 5 Depletion of neuronal NMII induces rearrangements of NMII$^{HC}$/Zip pre- and postsynaptically** | | | |
| **Fig. 5A:** Endogenous NMII$^{HC}$/Zip (cyan) at NMJs (HRP – magenta) in fixed larvae expressing *CTRL$^{35785}$*, *Zip$^{65947}$*, and *Sqh$^{32439}$* RNAi in neurons (*C155-Gal4*) | *C155-Gal4/Y;;CTRL$^{35785}$/+*<br><br>*C155-Gal4/Y;; NMII$^{HC}$/Zip$^{65947}$ /+*<br><br>*C155-Gal4/Y* | | | |

| | | | | |
|---|---|---|---|---|
| Repeated in two independent experiments, multiple crosses | *;;NMII*$^{LC}$*/Sqh*$^{32439}$*/+* | | | |
| **Fig. 5B:** Quantification of Zip fluorescence measured between the area masked by the neuronal HRP signal, (inner yellow outline) and 1 µm outside the neuron in the postsynaptic area (outer yellow outline), along with analysis of the number, area and fluorescence of WEKA-segmented individual Zip particles in this postsynaptic compartment. N in bar graphs – NMJs; N in box and whiskers – individual Zip assemblies. | *C155-Gal4/Y;;CTRL*$^{35785}$*/+*<br><br>*C155-Gal4/Y;; NMII*$^{HC}$*/Zip*$^{65947}$ */+*<br><br>*C155-Gal4/Y ;;NMII*$^{LC}$*/Sqh*$^{32439}$*/+* | 20 NMJs (10 larvae)<br><br>18 NMJs (10 larvae)<br><br>18 NMJs (9 larvae) | Postsynaptic Zip fluorescence per µm$^2$ | * p=0.02 and *** p<0.001 after One-way ANOVA with Kruskal-Wallis multiple comparison test.<br>N – NMJs |
| | | 19 NMJs (10 larvae)<br><br>16 NMJs (8 larvae)<br><br>19 NMJs (9 larvae) | Number of Zip spots per µm$^2$ | * p=0.02 and *** p<0.001 after One-way ANOVA with Kruskal-Wallis multiple comparison test.<br>N – NMJs |
| | | 7966 Zip spots (19 NMJs/10 larvae)<br>8591 Zip spots (16 NMJs/8 larvae)<br>8384 Zip spots (19 NMJs/9 larvae) | Area of Zip spots (µm$^2$) | *** p<0.001 after One-way ANOVA with Kruskal-Wallis multiple comparison test.<br>N – Zip spots |
| | | 7966 Zip spots (19 NMJs/10 larvae)<br>8591 Zip spots (16 NMJs/8 larvae)<br>8384 Zip spots (19 NMJs/9 larvae) | Fluorescence of Zip spots per µm$^2$ | *** p<0.001 after One-way ANOVA with Kruskal-Wallis multiple comparison test.<br>N – Zip spots |
| **Fig. 5C:** Quantification of Zip fluorescence in different muscle ROIs, along with analysis of the number, area and fluorescence of WEKA- | *C155-Gal4/Y;;CTRL*$^{35785}$*/+*<br><br>*C155-Gal4/Y;; NMII*$^{HC}$*/Zip*$^{65947}$ */+* | 55 Muscle ROIs (20 NMJs/10 larvae) | Muscle Zip fluorescence per µm$^2$ | ** p=0.009 and *** p<0.001 after One-way ANOVA with Kruskal-Wallis |

| | | | | |
|---|---|---|---|---|
| segmented individual Zip aggregate-like structures in muscles. | *C155-Gal4/Y ;;NMII$^{LC}$/Sqh$^{32439}$/+* | 61 Muscle ROIs (18 NMJs/9 larvae)<br><br>70 Muscle ROIs (18 NMJs/10 larvae) | | multiple comparison test.<br>N – Muscle ROIs |
| | | 40 Muscle ROIs(20 NMJs/10 larvae)<br>68 Muscle ROIs (18 NMJs/9 larvae)<br>46 Muscle ROIs (18 NMJs/10 larvae) | Number of Zip aggregates per µm$^2$ | *** $p<0.001$ and ** $p=0.005$ after One-way ANOVA with Kruskal-Wallis multiple comparison test.<br>N – Muscle ROIs |
| | | 2027 Zip aggregates (40 Muscle ROIs/20 NMJs/10 larvae)<br>3018 Zip aggregates (68 Muscle ROIs/18 NMJs/9 larvae)<br><br>3773<br>(46 Muscle ROIs /18 NMJs/10 larvae) | Area of Zip aggregates (µm$^2$) | ns – not significant, after One-way ANOVA with Kruskal-Wallis multiple comparison test.<br>N – Zip aggregates |
| | | 2027 Zip aggregates (40 Muscle ROIs/20 NMJs/10 larvae)<br>3018 Zip aggregates (68 Muscle ROIs/18 NMJs/9 larvae)<br><br>3773 Zip aggregates | Fluorescence of Zip aggregates per µm$^2$ | ns – not significant, after One-way ANOVA with Kruskal-Wallis multiple comparison test.<br>N – Zip aggregates |

| | | | | |
|---|---|---|---|---|
| | | (46 Muscle ROIs /18 NMJs/10 larvae) | | |
| **Figure 6.** | **Fig. 6. Neuronal depletion of non-muscle Myosin II rearranges integrin receptors at the NMJ.** | | | |
| **Fig. 6A:** Distribution of YFP-tagged, ubiquitously expressed Integrin-β (cyan) at NMJs in live larvae expressing Lifeact::Halo (magenta) in neurons. See also **Supp. Movie S8**. **BE107** | *C155-Gal4/Y;UAS-Lifeact::Halo/+; ubiIntegrin-β PS::YFP/+* | | | None |
| **Fig. 6B:** Immunostaining of Integrin-β (cyan) at NMJs of control larvae. Repeated in two independent experiments, multiple crosses | *C155-Gal4/Y;;UAS-CTRL$^{35785}$/+* | | | None |
| **Fig. 6C:** Reorganization of Integrin-β at NMJs of larvae expressing *NMII$^{HC}$/Zip$^{65947}$* and *NMII$^{LC}$/Sqh$^{32439}$* in neurons, compared to controls. | *C155-Gal4/Y;;UAS-CTRL$^{35785}$/+*<br><br>*C155-Gal4/Y;;UAS- Zip$^{65947}$/+*<br><br>*C155-Gal4/Y ;;NMII$^{LC}$/Sqh$^{32439}$/+* | | | None |
| **Fig. 6D:** Analysis of the Integrin-β fluorescence per μm$^2$ within the HRP delineated NMJ area , as well as in postsynaptic and muscle ROIs. | *C155-Gal4/Y;;UAS-CTRL$^{35785}$/+*<br><br>*C155-Gal4/Y;;UAS- Zip$^{65947}$/+*<br><br>*C155-Gal4/Y ;;NMII$^{LC}$/Sqh$^{32439}$/+* | 9 larvae/ 17 NMJs<br>10 larvae/ 20 NMJs<br>7 larvae/ 14 NMJs | Integrin-β fluorescence per μm$^2$ within the HRP-delineated NMJ area | ns – not significant, and *** p<0.001 after One-way ANOVA with Kruskal-Wallis multiple comparison test.<br><br>N – NMJs |
| | | 9 larvae/ 17 NMJs<br>10 larvae/ 20 NMJs<br>7 larvae/ 14 NMJs | Postsynaptic Integrin-β fluorescence per μm$^2$ | ** p=0.006 and *** p<0.001 after One-way ANOVA with Kruskal-Wallis multiple comparison test.<br><br>N – NMJs |
| | | 9 larvae/ 17 NMJs/63 muscle ROI<br><br>10 larvae/ 20 NMJs/61 muscle ROI | Muscle Integrin-β fluorescence per μm$^2$ | *** p<0.001 after One-way ANOVA with Kruskal-Wallis multiple comparison test.<br><br>N – muscle ROIs |

| | | | | |
|---|---|---|---|---|
| | | 7 larvae/ 14 NMJs/62 muscle ROI | | |
| **Fig. 6E:** Quantification of the percentage of NMJ area occupied by linear Integrin-β structures, as well as the area and fluorescence of WEKA-segmented individual particles. | *C155-Gal4/Y;;UAS-CTRL$^{35785}$/+*<br><br>*C155-Gal4/Y;;UAS- Zip$^{65947}$/+*<br><br>*C155-Gal4/Y ;;NMII$^{LC}$/Sqh$^{32439}$/+* | 9 larvae/ 17 NMJs<br>10 larvae/ 20 NMJs<br>7 larvae/ 14 NMJs | NMJ area covered with linear Integrin-β (%) | * $p<0.05$ and ** $p<0.01$ after One-way ANOVA with Kruskal-Wallis multiple comparison test.<br><br>N – NMJs |
| | | 9 larvae/ 17 NMJs/ 260 entities<br><br>10 larvae/ 20 NMJs/ 219 entities<br><br>7 larvae/ 14 NMJs/ 160 entities | Linear Integrin-β area (μm$^2$) | *** $p<0.001$ after One-way ANOVA with Kruskal-Wallis multiple comparison test.<br><br>N – Linear Integrin-β entities |
| | | 9 larvae/ 17 NMJs/ 260 entities<br><br>10 larvae/ 20 NMJs/ 219 entities<br><br>7 larvae/ 14 NMJs/ 160 entities | Linear Integrin-β fluorescence per μm$^2$ | *** $p<0.001$ after One-way ANOVA with Kruskal-Wallis multiple comparison test.<br><br>N – Linear Integrin-β entities |
| **Fig. 6F:** Analysis of the number of Integrin-β foci and the area and fluorescence of WEKA-segmented individual particles within the HRP-delineated NMJ area. | *C155-Gal4/Y;;UAS-CTRL$^{35785}$/+*<br><br>*C155-Gal4/Y;;UAS- Zip$^{65947}$/+*<br><br>*C155-Gal4/Y ;;NMII$^{LC}$/Sqh$^{32439}$/+* | 9 larvae/ 17 NMJs<br>10 larvae/ 20 NMJs<br>7 larvae/ 14 NMJs | Number of Integrin-β spots per μm$^2$ | *** $p<0.001$ and ns – non-significant after One-way ANOVA with Kruskal-Wallis multiple comparison test. |

| | | | | |
|---|---|---|---|---|
| | | | | N – NMJs |
| | | 9 larvae/ 17 NMJs/4013 entities<br><br>10 larvae/ 20 NMJs/4162 entities<br><br>7 larvae/ 14 NMJs/3683 entities | Integrin-β spots area ($\mu m^2$)<br><br>Integrin-β spots fluorescence per $\mu m^2$ | *** p<0.001 after One-way ANOVA with Kruskal-Wallis multiple comparison test.<br><br>N – Integrin-β spots |
| **Fig. 6G:** Analysis of the fluorescence per $\mu m^2$ of postsynaptic Integrin-β, as well as the number of postsynaptic Integrin-β foci and the area and fluorescence of WEKA-segmented individual particles. | *C155-Gal4/Y;;UAS-CTRL$^{35785}$/+*<br><br>*C155-Gal4/Y;;UAS- Zip$^{65947}$/+*<br><br>*C155-Gal4/Y ;;NMII$^{LC}$/Sqh$^{32439439}$/+* | 9 larvae/ 17 NMJs<br><br>10 larvae/ 20 NMJs<br><br>7 larvae/ 14 NMJs | Number of Integrin-β spots per $\mu m^2$ | ** p<0.01, and ns – non-significant upon after One-way ANOVA with Kruskal-Wallis multiple comparison test.<br><br>N – NMJs |
| | | 9 larvae/ 17 NMJs/7857 spots<br><br>10 larvae/ 20 NMJs/7938 spots<br><br>7 larvae/ 14 NMJs/6667 spots | Integrin-β spots' area ($\mu m^2$)<br><br>Integrin-β spots' fluorescence per $\mu m^2$ | *** p<0.001 after One-way ANOVA with Kruskal-Wallis multiple comparison test.<br><br>N – Integrin-β spots |
| **Figure 7.** | **Fig. 7. Mechanical severing of larval axons induces changes in presynaptic F-actin.** | | | |

| | | | | |
|---|---|---|---|---|
| **Fig. 7A:** Diagram of *Drosophila* fillet preparation upon axotomy (light blue dashed line) of all proximal neurons descending from the larval brain. This procedure removes the axonal tension exerted at presynaptic compartments. | | | | |
| **Fig. 7B:** mNGMA-labeled presynaptic actin at NMJs from live larvae with intact brains (left) or upon severing of the brain/axotomy (right). See also **Supp. Movies S9-10**.<br>Repeated in two independent experiments, multiple crosses | *C155-Gal4/Y;QUAS-mNeonGreenMA(GMA)/+; nSyb-QF2/+* | Intact brains | | None |
| | *C155-Gal4/Y;QUAS-mNeonGreenMA(GMA)/+; nSyb-QF2/+* | Axotomy | | |
| **Fig. 7C:** Quantification of the fluorescence per µm$^2$ for the QmNGMA actin marker in whole NMJs.<br>Repeated in two independent experiments, multiple crosses | *C155-Gal4/Y;QUAS-mNeonGreenMA(GMA)/+; nSyb-QF2/+* | <u>Intact brain:</u><br>8 larvae/17 NMJs<br><br><u>Axotomy:</u><br>8 larvae/17 NMJs | Actin fluorescence per µm$^2$ | ns – not significant, ** $p<0.01$, and *** $p<0.001$ upon unpaired, non-parametric Mann-Whitney test. |
| **Fig. 7D:** Quantifications of the number of round F-actin assemblies per µm$^2$, as well as the area and fluorescence of the individual round presynaptic actin structures. | *C155-Gal4/Y;QUAS-mNeonGreenMA(GMA)/+; nSyb-QF2/+* | <u>Intact brain:</u><br>8 larvae/17 NMJs<br><br><u>Axotomy:</u><br>8 larvae/17 NMJs | Number of round F-actin assemblies per µm$^2$ | |
| | | <u>Intact brain:</u><br>8 larvae/17 NMJs/400<br><br><u>Axotomy:</u><br>8 larvae/17 NMJs/600 | Area per round F-actin assembly (µm$^2$) | |
| | | | Round F-actin fluorescence per µm$^2$ | |
| **Fig. 7E:** Graphs representing the percentage of NMJ area covered with linear F-actin, as well as the area and fluorescence of the individual assemblies. | *C155-Gal4/Y;QUAS-mNeonGreenMA(GMA)/+; nSyb-QF2/+* | <u>Intact brain:</u><br>8 larvae/17 NMJs<br><br><u>Axotomy:</u><br>8 larvae/17 NMJs | NMJ area covered with linear F-actin (%) | |

| | | | | |
|---|---|---|---|---|
| | | Intact brain:<br>8 larvae/17 NMJs/123<br><br>Axotomy:<br>8 larvae/17 NMJs/132 | Area per linear F-actin assembly ($\mu m^2$)<br><br>Linear F-actin fluorescence per $\mu m^2$ | |
| **Fig. 7F:** mNGMA-labeled presynaptic actin at NMJs from live *NMII$^{HC}$/Zip$^{65947}$* larvae with intact brains (left) or upon severing of the brain/axotomy (right). | *C155-Gal4/Y; QUAS-mNeonGreenMA(GMA)/UAS-Zip$^{65947}$; nSyb-QF2/+* | | | |
| **Fig. 7G:** Analyses of the mNGMA fluorescence. Repeated in two independent experiments, multiple crosses | *C155-Gal4/Y; QUAS-mNeonGreenMA(GMA)/UAS-Zip$^{65947}$; nSyb-QF2/+* | Intact brain:<br>10 larvae/25 NMJs<br><br>Axotomy:<br>10 larvae/26 NMJs | Actin fluorescence per $\mu m^2$ | * p=0.0273 upon unpaired, non-parametric Mann-Whitney test.<br>N – NMJs |
| **Fig. 7H:** Analyses of the round F-actin assemblies. | | Intact brain:<br>10 larvae/25 NMJs<br><br>Axotomy:<br>10 larvae/26 NMJs | Number of round F-actin assemblies per $\mu m^2$ | *** p=0.0010 upon unpaired, non-parametric Mann-Whitney test.<br>N – NMJs |
| | *C155-Gal4/Y; QUAS-mNeonGreenMA(GMA)/UAS-Zip$^{65947}$; nSyb-QF2/+* | Intact brain:<br>658 spots (10 larvae/25 NMJs)<br>Axotomy:<br>1124 spots (10 larvae/26 NMJs) | Area per round F-actin assembly ($\mu m^2$)<br><br>Round F-actin fluorescence per $\mu m^2$ | Ns – non-significant upon unpaired, non-parametric Mann-Whitney test. |
| **Fig. 7I:** Analyses of the linear F-actin assemblies. | *C155-Gal4/Y; QUAS-mNeonGreenMA(GMA)/UAS-Zip$^{65947}$; nSyb-QF2/+* | Intact brain:<br>10 larvae/25 NMJs<br><br>Axotomy:<br>10 larvae/26 NMJs | NMJ area covered with linear F-actin (%) | ns – not significant upon unpaired, non-parametric Mann-Whitney test.<br>N – NMJs |
| | | Intact brain:<br>561 structures (10 larvae/25 NMJs)<br>Axotomy:<br>534 structures (10 larvae/26 NMJs) | Area per linear F-actin assembly ($\mu m^2$)<br><br>Linear F-actin fluorescence per $\mu m^2$ | ns – not significant upon unpaired, non-parametric Mann-Whitney test.<br>N – linear F-actin assemblies |

| | | | | |
|---|---|---|---|---|
| **Figure 8** | **Fig. 8. Integrin receptors rearrange within 15 minutes of mechanical severing of larval axons.** | | | |
| **Fig. 8A:** Diagrams of *Drosophila* fillet preparation upon axotomy (left, light blue dashed line), a procedure that should decrease the axonal tension exerted at presynaptic compartments. | | | | |
| **Fig. 8B:** Immunostaining of Integrin-β (cyan) at NMJs of control larvae (*C155-Gal4>CTRL*$^{35785}$) after 15 minutes with uncut and cut axons. | *C155-Gal4/Y;;UAS-CTRL*$^{35785}$*/+* | | | |
| **Fig. 8C:** Integrin-β fluorescence in the HRP-delineated-, postsynaptic-, and muscle area. | *C155-Gal4/Y;;UAS-CTRL*$^{35785}$*/+*<br>Uncut<br><br>Cut | Uncut: 10 larvae/ 21 NMJs<br><br>Cut: 8 larvae/ 16 NMJs | Integrin-β fluorescence in the HRP-delineated- and postsynaptic area | ns – not significant, upon unpaired, non-parametric Mann-Whitney test.<br><br>N – NMJs |
| | | Uncut: 10 larvae/ 21 NMJs/ 56 muscle ROIs<br><br>Cut: 8 larvae/ 16 NMJs/ 31 muscle ROIs | Integrin-β fluorescence in muscles | ** p<0.01 upon unpaired, non-parametric Mann-Whitney test.<br><br>N – muscle ROIs |
| **Fig. 8D:** Quantifications of the abundance, size and fluorescence of WEKA-segmented Integrin-β structures' along linear areas.<br><br>Repeated in two independent experiments, multiple crosses. | *C155-Gal4/Y;;UAS-CTRL*$^{35785}$*/+*<br>Uncut<br><br><br>Cut | Uncut: 10 larvae/ 21 NMJs<br><br>Cut: 8 larvae/ 16 NMJs | NMJ area covered with linear Integrin-β (%) | ns – not significant, upon unpaired, non-parametric Mann-Whitney test.<br><br>N – NMJs |

| | | | | |
|---|---|---|---|---|
| | | Uncut: 10 larvae/ 21 NMJs/ 154 entities<br><br>Cut: 8 larvae/ 16 NMJs/ 104 entities | Linear Integrin-β area ($\mu m^2$)<br><br>Linear Integrin-β fluorescence per $\mu m^2$ | *p=0.0186 upon unpaired, non-parametric Mann-Whitney test.<br><br>N – Integrin-β entities |
| **Supplemental figure 1.** | **Segmentation and analysis of round and linear F-actin assemblies.** | | | |
| **Supplemental Figure 2.** | **Supplemental Figure 2. Presynaptic actin cytoskeleton at the larval *Drosophila* neuromuscular junction.** | | | |
| **Fig. S2A:** Time-lapse insets (from **Supp. Movie S1**) showing the structure and lifetime of linear F-actin/cable-like structures at the NMJ. | *C155-Gal4/Y;QUAS-mNeonGreenMA(GMA)/+; nSyb-QF2/+* | | | None |
| **Fig.S2B:** Time-lapse insets (from **Supp. Movie S1**) of spot-like presynaptic F-actin assemblies (pointed with green circles) with different lifetimes (green circles point a longer-lived assembly, light-blue circle shows a short-lived, possibly endocytic event) visualized via the actin marker QmNGMA at the NMJ. | *C155-Gal4/Y;QUAS-mNeonGreenMA(GMA)/+; nSyb-QF2/+* | | | None |
| **Fig.S2C:** Time-lapse of a fluorescence recovery after photobleaching (FRAP) of the neuronally expressed actin marker QmNGMA at a bouton containing linear F-actin assemblies. The bleached bouton is outlined in yellow. | *C155-Gal4/Y;QUAS-mNeonGreenMA(GMA)/+; nSyb-QF2/+* | | | |
| **Fig.S2D:** FRAP quantification of linear F-actin-containing region showing recovery of QmNGMA.<br>One independent experiment, multiple crosses | *C155-Gal4/Y;QUAS-mNeonGreenMA(GMA)/+; nSyb-QF2/+* | | 2 larvae/ 5 NMJs | None |
| **Fig.S2E** Inverted contrast montage of the maximum intensity projection, along with individual Z-stacks of | *C155-Gal4/Y; QUAS-Act5C::mScarI/+; nSyb-QF2/+* | | | |

| | | | | |
|---|---|---|---|---|
| an NMJ from a larvae expressing mScarlet-I::Act5C neuronally (*nSybQF>QAct5C::mScarI*) | | | | |
| **Fig.S2F:** Presynaptic actin assemblies are sensitive to a treatment with the actin depolymerization-inducing drug Latrunculin A (LatA). One independent experiment, multiple crosses. | *C155-Gal4/Y;QUAS-mNeonGreenMA(GMA)/+; nSyb-QF2/+* | **DMSO condition:** 5 larvae/ 12 NMJs<br><br>**LatA treatment:** 6 larvae/ 18 NMJs | | |
| **Fig.S2G:** Quantifications of the fluorescence per $\mu m^2$ and coefficient of variation (CoV) for the QmNGMA actin marker in DMSO and LatA-treated larval fillets, measured through the middle of NMJs, where the actin core predominantly traverses. One independent experiment, multiple crosses. | *C155-Gal4/Y;QUAS-mNeonGreenMA(GMA)/+; nSyb-QF2/+* | **DMSO condition:** 5 larvae/ 12 NMJs<br><br>**LatA treatment:** 6 larvae/ 18 NMJs | Fluorescence per $\mu m^2$ and coefficient of variation (CoV) for the QmNGMA actin marker | ns – not significant, * $p<0.05$, ** $p<0.01$, and *** $p<0.001$ upon unpaired, non-parametric Mann-Whitney test.<br><br>N – NMJs |
| **Supplemental Figure 3** | **Supplemental Figure 3. Assessment for linear F-actin in axons.** | | | |
| **Fig.S3A:** Diagrams of *Drosophila* fillet preparations denoting proximal and distal axons along the larva. | | | | |
| **Fig.S3B:** Distal axons (blue arrows) of larvae expressing neuronally (*nSybQF*) the F-actin marker QmNGMA feature possible linear F-actin structures along with visible linear F-actin at the NMJ. By contrast, larvae expressing neuronal (*C155-Gal4*) GFP exhibit diffuse GFP signal both in the distal axon and NMJ. One independent experiment, multiple crosses. | *C155-Gal4/Y;QUAS-mNeonGreenMA(GMA)/+; nSyb-QF2/+*<br><br>*C155-Gal4/Y;GFP/+* | | | |
| **Fig. S3C:** In the narrow axonal shafts of proximal axonal bundles, putative linear F-actin QmNGMA structures are not distinguishable from the | *C155-Gal4/Y;QUAS-mNeonGreenMA(GMA)/+; nSyb-QF2/+*<br><br>*C155-Gal4/Y;GFP/+* | 45 axons/5 larvae<br><br>26 axons/6 larvae | Linear structures covered area (%) | ns – not significant, upon unpaired, non-parametric Mann-Whitney test. |

| | | | | |
|---|---|---|---|---|
| distribution of free GFP, either visually or quantified via WEKA segmentation. | | | | N – axons |
| **Supplemental Figure 4** | **Supplemental Figure 4. Actin-associated proteins at the presynaptic actin core.** | | | |
| **Fig. S4A:** Live imaging of the branched actin nucleator Arp2/3, visualized via Arp3::GFP (cyan), shows decoration of spot-like and linear F-actin assemblies labeled with Lifeact::Halo (magenta). See also **Supp. Movie S2**.<br>One independent experiment, multiple crosses. | *C155-Gal4/Y;UAS-Arp3::GFP/UAS-Lifeact::Halo* | | | |
| **Fig. S4B:** The linear actin nucleator formin/Dia::GFP (cyan) decorates actin cables (magenta), in addition to its punctate distribution throughout the NMJ (see also **Supp. Movie S3**).<br>One independent experiment, multiple crosses. | *C155-Gal4/Y;UAS-Dia::EGFP/UAS-Lifeact::Halo* | | | |
| **Fig. S4C:** Diffuse cytosolic GFP (cyan) shows no specific enrichment or co-localization with presynaptic actin assemblies (magenta). See also **Supp. Movie S4**.<br>One independent experiment, multiple crosses | *C155-Gal4/Y;UAS-GFP/UAS-Lifeact::Halo* | | | |
| **Fig. S4D:** Distribution of the non-muscle myosin II light chain subunit sqh::GFP at the NMJ in live larvae expressing Lifeact::Halo in neurons.<br>One independent experiment, multiple crosses. | *C155-Gal4/Y;UAS-Lifeact::Halo/+; sqh::GFP/+* | | | |
| **Supplemental Figure 5.** | **Supplemental Figure 5. Association of the F-actin-stabilizing protein Tropomyosin with linear F-actin structures in the presynaptic core** | | | |
| **Fig.S5A-B:** Inverted contrast single frame from live imaging depicting the localization pattern upon neuronal expression of mNeonGreen-tagged Tm1- | *C155-Gal4/Y;QUAS-mNeonGreen::Tm1-L/+; nSyb-QF2/+* | | | |

| | | | | |
|---|---|---|---|---|
| L (mNG-Tm1-L or "long" Tm1) and Tm1-A (mN-Tm1-A or "short" Tm1). One independent experiment, multiple crosses. | *C155-Gal4/Y;QUAS-mNeonGreen::Tm1-A/+; nSyb-QF2/+* | | | |
| **Fig.S5C:** The F-actin stabilizer Tm1-A (mNG-Tm1-A) is enriched in the linear Lifeact::Halo-marked actin assemblies along the presynaptic core. | *C155-Gal4/Y;UAS-Lifeact::Halo/QUAS-mNeonGreenTm1-L; nSyb-QF2/+*<br><br>*C155-Gal4/Y;UAS-Lifeact::Halo/UAS-GFP* | | | |
| **Fig.S5D:** Pearson's R correlation between Lifeact::Halo and GFP or mNG-Tm1-A. | *C155-Gal4/Y;QUAS-mNeonGreen::Tm1-A/UAS-GFP; nSyb-QF2/+*<br><br>*C155-Gal4/Y;UAS-Lifeact::Halo/UAS-GFP* | 7 larvae/20 NMJs<br><br>8 larvae/22 NMJs | Pearson's R correlation | *** p<0.001 upon unpaired, non-parametric Mann-Whitney test.<br><br>N – NMJs |
| **Fig.S5E-F:** Presynaptic localization of QmNGMA-labeled actin assemblies and mNG-Tm1-L is sensitive to treatment with LatA, but not to the vehicle DMSO, supporting F-actin-Tm1 association along the presynaptic actin core. One independent experiment, multiple crosses. | *C155-Gal4/Y;QUAS-mNeonGreenMA(GMA)/+; nSyb-QF2/+*<br><br>*C155-Gal4/Y;UAS-Lifeact::Halo/QUAS-mNeonGreenTm1-L; nSyb-QF2/+* | DMSO: 3 larvae/ 9 NMJs<br>LatA: 4 larvae/12 NMJs<br><br>DMSO: 3 larvae/ 9 NMJs<br>LatA: 4 larvae/ 9 NMJs | | None |
| **Fig.S5G:** Endogenous Tm1 (cyan), detected via immunofluorescence, traverses the NMJ (labeled with HRP – magenta) in larvae expressing control RNAi in all neurons. The Tm1 signal is significantly decreased at NMJs of animals neuronally expressing a Tm1 RNAi line targeting all known Tm1 isoforms (*panTm1*$^{43542}$), and the complementary line *Tm1*$^{56869}$, targeting the Tm1 isoforms A, Q and R. | *C155-Gal4/Y;; UAS-CTRL*$^{35785}$*/+*<br><br>*C155-Gal4/Y;; UAS- Tm1*$^{43542}$*/+*<br><br>*C155-Gal4/Y;; UAS- Tm1*$^{56869}$*/+* | | | |
| **Fig.S5H:** Quantification of the Tm1-A/L fluorescence per µm$^2$ at NMJs from larvae neuronally expressing *CTRL*, *Tm1*$^{43542}$ or *Tm1*$^{56869}$ RNAi lines. | *C155-Gal4/Y;; UAS-CTRL*$^{35785}$*/+*<br><br>*C155-Gal4/Y;; UAS- Tm1*$^{43542}$*/+*<br><br>*C155-Gal4/Y;; UAS- Tm1*$^{56869}$*/+* | 8 larvae/28 NMJs<br><br>5 larvae/13 | Tm1-A/L fluorescence per µm$^2$ | *** p<0.001 upon one way ANOVA with Kruskal-Wallis multicomparison test. |

| | | | | |
|---|---|---|---|---|
| | | 7 larvae/14 NMJs | | N – NMJs |
| **Supplemental figure 6.** | **Supplemental figure 6. Validation of *UAS-RNAi* lines targeting NMII$^{HC}$/Zip and NMII$^{LC}$/Sqh.** | | | |
| **Fig.S6A:** Distribution of Sqh::GFP (cyan) in axons (labeled with HRP in magenta) proximal to muscle 4 in Sqh::GFP/+ heterozygous larvae, and larvae expressing in neurons two independent *UAS-RNAi* lines ($Zip^{65947}$ and $Zip^{36727}$) targeting NMII$^{HC}$/Zip, as well as the NMII$^{LC}$/Sqh targeting lines $Sqh^{32439}$ and $Sqh^{33892}$. | *C155-Gal4/Y;; sqh::GFP/+*<br><br>*C155-Gal4/Y;; sqh::GFP/UAS-*$Zip^{65947}$<br>*C155-Gal4/Y;; sqh::GFP/UAS-*$Zip^{36727}$<br>*C155-Gal4/Y;; sqh::GFP/UAS-*$Sqh^{32439}$<br>*C155-Gal4/Y;; sqh::GFP/UAS-*$Sqh^{33892}$ | | | |
| **Fig.S6B:** Quantification of the Sqh::GFP fluorescence in axons of larvae with the different genotypes shown in **A**. | *C155-Gal4/Y;; sqh::GFP/+*<br><br>*C155-Gal4/Y;; sqh::GFP/UAS-*$Zip^{65947}$<br>*C155-Gal4/Y;; sqh::GFP/UAS-*$Zip^{36727}$<br>*C155-Gal4/Y;; sqh::GFP/UAS-*$Sqh^{32439}$<br>*C155-Gal4/Y;; sqh::GFP/UAS-*$Sqh^{33892}$ | 4 larvae/ 16 NMJs<br>4 larvae/ 8 NMJs<br>4 larvae/ 8 NMJs<br>4 larvae/ 16 NMJs<br>3 larvae/ 6 NMJs | Axonal sqh::GFP fluorescence per $\mu m^2$ | ns – not significant, *** $p<0.001$ after One-way ANOVA with Kruskal-Wallis multiple comparison test. |
| **Fig.S6C:** Example of image analysis in Fiji, where the Sqh::GFP fluorescence was measured within the HRP delineated NMJ area, in a 1 µm rim surrounding the bouton (postsynaptic ROI), and in the muscle ROI (rectangle with yellow dashed lines) (excluding the presynaptic compartment and the surrounding 1 µm rim). | | | | |
| **Fig.S6D:** Quantifications of Sqh::GFP fluorescence at NMJs from larvae carrying $Zip^{65947}$ in absence and presence of the neuronal *C155-Gal* driver, in an experiment validating the lack of nonspecific expression of $Zip^{65947}$ in neurons and muscles in absence of Gal4 driver. | *X/Y;; sqh::GFP/UAS-CTRL*$^{35785}$<br><br>*C155-Gal4/Y;; sqh::GFP/UAS-CTRL*$^{35785}$<br><br>*X/Y;; sqh::GFP/UAS-* $Zip^{65947}$<br><br>*C155-Gal4/Y;; sqh::GFP/UAS-*$Zip^{65947}$ | 7 larvae/14 NMJs<br><br>7 larvae/14 NMJs<br><br>5 larvae/10 NMJs<br><br>6 larvae/14 NMJs | sqh::GFP fluorescence per $\mu m^2$ | ns – not significant, *** $p<0.001$ after One-way ANOVA with Kruskal-Wallis multiple comparison test. |

| | | | | |
|---|---|---|---|---|
| One independent experiment, multiple crosses. | | | | |
| **Fig.S6E:** Immunostaining of endogenous Zip (cyan) at NMJs (HRP magenta) and muscles of larvae expressing *C57-Gal4*-driven control and RNAi lines targeting NMII$^{HC}$/Zip and NMII$^{LC}$/Sqh, respectively. Two independent experiments, multiple crosses. | *C155-Gal4/Y;; UAS-CTRL$^{35785}$/+*<br><br>*C155-Gal4/Y;; UAS- Zip$^{65947}$/+*<br><br>*C155-Gal4/Y;; sqh::GFP/UAS-Sqh$^{32439}$/+* | | | |
| **Fig.S6F:** Quantifications of muscle Zip fluorescence in larvae expressing *C57-Gal4*-driven *CTRL$^{35785}$*, *Zip$^{65947}$* and *Sqh$^{32439}$*. | *C57-Gal4/Y;; UAS-CTRL$^{35785}$/+*<br>*C57-Gal4/Y;; UAS- Zip$^{65947}$/+*<br>*C57-Gal4/Y;; sqh::GFP/UAS-Sqh$^{32439}$/+* | 52 muscle ROIs (10 larvae/20 NMJs)<br>37 muscle ROIs (8 larvae/16 NMJs)<br>45 muscle ROIs (9 larvae/18 NMJs) | Zip fluorescence per µm$^2$ | ns – not significant, *** p<0.001 after One-way ANOVA with Kruskal-Wallis multiple comparison test.<br>N – muscle ROIs |
| **Fig.S6G:** Zip particle abundance, size and brightness in larvae expressing *C57-Gal4*-driven *CTRL$^{35785}$*, *Zip$^{65947}$* and *Sqh$^{32439}$*. | *C57-Gal4/Y;; UAS-CTRL$^{35785}$/+*<br><br>*C57-Gal4/Y;; UAS- Zip$^{65947}$/+*<br><br>*C57-Gal4/Y;; sqh::GFP/UAS-Sqh$^{32439}$/+* | 20 Muscle ROIs (10 larvae/20 NMJs)<br>15 muscle ROIs (8 larvae/16 NMJs)<br>17 muscle ROIs (9 larvae/18 NMJs) | Number of Zip particles per µm$^2$ | *** p<0.001 after One-way ANOVA with Kruskal-Wallis multiple comparison test.<br>N – muscle ROIs |
| | | 54133 spots (10 larvae/20 NMJs)<br>32366 spots (8 larvae/16 NMJs)<br>40796 spots (9 larvae/18 NMJs) | Area per Zip particle (µm$^2$)<br><br>Zip particles' fluorescence per µm$^2$ | *** p<0.001 after One-way ANOVA with Kruskal-Wallis multiple comparison test.<br>N – Zip spots |
| **Supplemental figure 7.** | **Supplemental figure 7. Presynaptic actin assemblies upon down-regulation and overexpression of NMII$^{HC}$/Zip in neurons.** | | | |
| **Fig.S7A:** Inverted contrast mNGMA-labeled presynaptic actin in fixed larvae co-expressing | *C155-Gal4/Y;QUAS-mNeonGreenMA(GMA)/UAS-CTRL$^{35785}$; nSyb-QF2/+* | 6 larvae (3 male+3 female)/24 NMJs | | |

| | | | | |
|---|---|---|---|---|
| *CTRL*$^{35785}$- or *NMII*$^{HC}$*/Zip*$^{65947}$ RNAi lines in neurons. | *C155-Gal4/Y;QUAS-mNeonGreenMA(GMA)/UAS-Zip*$^{65947}$*; nSyb-QF2/+* | 6 larvae (3 male+3 female)/20 NMJs | | |
| **Fig.S7B:** Quantification of the fluorescence per µm$^2$ for the mNGMA actin marker in whole NMJs, along with the number of actin assemblies per µm$^2$, and the area and fluorescence intensity of the individual presynaptic actin structure. The actin particles were analyzed without specifying their morphological features as round or linear to account for fragmentation of the presynaptic cytoskeleton as a result of fixation[1]. | *C155-Gal4/Y;QUAS-mNeonGreenMA(GMA)/UAS-CTRL*$^{35785}$*; nSyb-QF2/+*<br><br>*C155-Gal4/Y;QUAS-mNeonGreenMA(GMA)/UAS-Zip*$^{65947}$*; nSyb-QF2/+* | 6 larvae (3 male+3 female)/24 NMJs<br><br>6 larvae (3 male+3 female)/20 NMJs | Actin fluorescence per µm$^2$ | ns – not-significant, ** $p<0.01$, *** $p<0.001$ upon unpaired, non-parametric Mann-Whitney test.<br>N – NMJs |
| | | | Number F-actin assemblies per µm$^2$ | |
| | | 6 larvae (3 male+3 female)/24 NMJs/2184<br><br>6 larvae (3 male+3 female)/20 NMJs/1950 | Area per F-actin assembly (µm$^2$)<br><br>F-actin assemblies' fluorescence per µm$^2$ | |
| **Fig.S7C:** Live-imaged inverted contrast Lifeact::Halo-labeled presynaptic actin in control larvae expressing GFP and larvae expressing NMII$^{HC}$/Zip::GFP.<br>Two independent experiments, multiple crosses. | *C155-Gal4/Y;UAS-Lifeact::Halo/UAS-GFP*<br><br>*C155-Gal4/Y;UAS-Lifeact::Halo/UAS-Zip::GFP* | | | |
| **Fig.S7D-E:** Analyses of the F-actin fluorescence, as well as individual WEKA-segmented round and linear F-actin entities in GFP- and Zip::GFP-expressing animals. | *C155-Gal4/Y;UAS-Lifeact::Halo/UAS-GFP*<br><br>*C155-Gal4/Y;UAS-Lifeact::Halo/UAS-Zip::GFP* | 22 NMJs/8 larvae<br><br>31 NMJs/9 larvae | Actin fluorescence per µm$^2$ | ns – not-significant, upon unpaired, non-parametric Mann-Whitney test.<br>N – NMJs |
| | | 22 NMJs/8 larvae<br>31 NMJs/9 larvae | Number of round F-actin assemblies per µm$^2$ | ns – not-significant, upon unpaired, non-parametric Mann-Whitney test.<br>N – NMJs |

| | | | | |
|---|---|---|---|---|
| | | 813 spots (22 NMJs/8 larvae)<br><br>933 spots (31 NMJs/9 larvae) | Area per round F-actin assembly ($\mu m^2$)<br><br>Round F-actin fluorescence per $\mu m^2$ | * p=0.0148 upon unpaired, non-parametric Mann-Whitney test.<br>N – actin spots |
| | | 22 NMJs/8 larvae<br>31 NMJs/9 larvae | NMJ area covered with linear F-actin (%) | ns – not-significant, upon unpaired, non-parametric Mann-Whitney test.<br>N – NMJs |
| | | 567 structures (22 NMJs/8 larvae)<br><br>903 structures (31 NMJs/9 larvae) | Area per linear F-actin assembly ($\mu m^2$)<br><br>Linear F-actin fluorescence per $\mu m^2$ | ** p=0.0028 upon unpaired, non-parametric Mann-Whitney test.<br><br>N – linear F-actin assemblies |
| **Supplemental Figure 8.** | **Supplemental Figure 8. Neuronal depletion of non-muscle Myosin II rearranges Integrin receptors at the NMJ.** | | | |
| **Fig.S8A:** Pearson's R correlation between Lifeact::Halo and Integrin-β::YFP at NMJs of live larvae.<br>One independent experiment, multiple crosses. | *C155-Gal4/Y;;UAS-Lifeact::Halo/+;ubiMys::YFP/+* | 2 larvae/6 NMJs | Pearson's R correlation | |
| **Fig.S8B-C:** Reorganization of Integrin-β at NMJs of larvae expressing an independent RNAi line *Zip*$^{36727}$ in neurons, compared to controls.<br>One independent experiment, multiple crosses. | *C155-Gal4/Y;;UAS-CTRL*$^{35785}$*/+*<br><br>*C155-Gal4/Y;;UAS- Zip*$^{36727}$*/+* | 10 larvae/20 NMJs<br><br>4 larvae /8 NMJs | Fluorescence of Integrin-β per $\mu m^2$ within the HRP-delineated NMJ area<br><br>Fluorescence of postsynaptic Integrin-β per $\mu m^2$ | Ns – non-significant, *** p<0.001, upon unpaired, non-parametric Mann-Whitney test.<br>N – NMJs |
| **Fig.S8D-F:** Fluorescence intensity of Integrin-β within the HRP-delineated NMJ area and the postsynaptic region in the | *X/Y;;UAS-CTRL*$^{35785}$*/+*<br><br>*C155-Gal4/Y;;UAS-CTRL*$^{35785}$*/+* | 8 larvae/ 16 NMJs<br><br>7 larvae / 14 NMJs | Integrin-β fluorescence per $\mu m^2$ within the HRP- | Ns – non-significant, ** p<0.01, *** p<0.001 upon unpaired, non- |

| | | | | |
|---|---|---|---|---|
| presence and absence of *C155-Gal4*.<br>One independent experiment, multiple crosses. | *X/Y;;UAS- Zip$^{65947}$/+*<br><br>*C155-Gal4/Y;;UAS- Zip$^{65947}$/+* | 7 larvae/ 14 NMJs<br><br>6 larvae / 13 NMJs | delineated NMJ area<br><br>Fluorescence of postsynaptic Integrin-β per μm$^2$ | parametric Mann-Whitney test.<br>N – NMJs |
| **Fig.S8G-H:** Fluorescence intensity of Integrin-β in myotendinous junctions in control and *Zip$^{65947}$* larvae.<br>One independent experiment, multiple crosses. | *C155-Gal4/Y;;UAS-CTRL$^{35785}$/+*<br><br>*C155-Gal4/Y;;UAS- Zip$^{65947}$/+* | 9 larvae/ 18 NMJs<br><br>8 larvae / 16 NMJs<br>The myotendinous junctions at muscles 6/7 imaged | Fluorescence of Integrin-β in myotendinous junctions per μm$^2$ | ns – non-significant, upon unpaired, non-parametric Mann-Whitney test.<br>N – myotendinous junctions |
| **Supplemental Figure 9** | **Supplemental Figure 9. Organization of integrin receptors at the NMJ upon Integrin-β neuronal down-regulation** | | | |
| **Fig. S9A:** Integrin-β (cyan) in axonal bundles (HRP in magenta) in control and larvae expressing the *Integrin-β/Mys$^{33642}$* RNAi in neurons (*C155-Gal4*).<br>Two independent experiments, multiple crosses. | *C155-Gal4/Y;;UAS-CTRL$^{35785}$/+*<br><br>*C155-Gal4/Y;;UAS- Mys$^{33642}$/+* | | | |
| **Fig. S9B:** Quantification of axonal Integrin-β fluorescence in *CTRL$^{35785}$* vs *Mys$^{33642}$* neuronal RNAi. | *C155-Gal4/Y;;UAS-CTRL$^{35785}$/+*<br><br>*C155-Gal4/Y;;UAS- Mys$^{33642}$/+* | 8 axons/8 larvae<br><br>10 axons/10 larvae | Axonal Integrin-β fluorescence per μm$^2$ | *** $p<0.0001$ upon unpaired, non-parametric Mann-Whitney test.<br>N – axons |
| **Fig. S9C:** Integrin-β (cyan) at NMJs (HRP in magenta) in control and *Integrin-β/Mys$^{33642}$* animals. | *C155-Gal4/Y;;UAS-CTRL$^{35785}$/+*<br><br>*C155-Gal4/Y;;UAS- Mys$^{33642}$/+* | | | |
| **Fig. S9D:** Analyses of Integrin-β fluorescence pre- and postsynaptically, as well as in muscles of control and *Integrin-β/Mys$^{33642}$* larvae. | *C155-Gal4/Y;;UAS-CTRL$^{35785}$/+*<br><br>*C155-Gal4/Y;;UAS- Mys$^{33642}$/+* | 9 larvae/17 NMJs<br>8 larvae/16 NMJs | Integrin-β fluorescence per μm$^2$ within the HRP-delineated NMJ area<br><br>Postsynaptic Integrin-β | ns – not significant upon unpaired, non-parametric Mann-Whitney test.<br>N – NMJs |

| | | | | |
|---|---|---|---|---|
| | | | fluorescence per $\mu m^2$ | |
| | | 9 larvae/17 NMJs/63 muscle ROIs<br><br>8 larvae/16 NMJs/50 muscle ROIs | Muscle Integrin-β fluorescence per $\mu m^2$ | ns – not significant upon unpaired, non-parametric Mann-Whitney test.<br>N – muscle ROIs |
| **Fig. S9E:** Integrin-β WEKA-segmented particles in controls and *Integrin-β/Mys$^{33642}$* larvae. | *C155-Gal4/Y;;UAS-CTRL$^{35785}$/+*<br><br>*C155-Gal4/Y;;UAS- Mys$^{33642}$/+* | | | |
| **Fig. S9F:** Analyses of spot-like Integrin-β structures within the postsynaptic ROI | *C155-Gal4/Y;;UAS-CTRL$^{35785}$/+*<br><br>*C155-Gal4/Y;;UAS- Mys$^{33642}$/+* | 9 larvae/18 NMJs<br><br>8 larvae/16 NMJs | Number of Integrin-β spots per $\mu m^2$ | ns – non-significant upon unpaired, non-parametric Mann-Whitney test.<br>N – NMJs |
| | *C155-Gal4/Y;;UAS-CTRL$^{35785}$/+*<br><br>*C155-Gal4/Y;;UAS- Mys$^{33642}$/+* | 9 larvae/18 NMJs/<br>7857 spots<br><br>8 larvae/16 NMJs/<br>7617 spots | Integrin-β spot's area ($\mu m^2$)<br><br>Integrin-β spot's fluorescence per $\mu m^2$ | *** $p<0.0001$ upon unpaired, non-parametric Mann-Whitney test.<br>N – Integrin-β spots |
